# Visualizing Structure and Transitions for Biological Data Exploration

**DOI:** 10.1101/120378

**Authors:** Kevin R. Moon, David van Dijk, Zheng Wang, Scott Gigante, Daniel B. Burkhardt, William S. Chen, Kristina Yim, Antonia van den Elzen, Matthew J. Hirn, Ronald R. Coifman, Natalia B. Ivanova, Guy Wolf, Smita Krishnaswamy

**Author notes:** Corresponding author. Address: 333 Cedar St, New Haven, CT 06510, USA. Correspondence for experiments. These authors contributed equally (Kevin R. Moon, David van Dijk, Zheng Wang, Scott Gigante). These authors contributed equally (Natalia B. Ivanova, Guy Wolf, Smita Krishnaswamy).

## Abstract

With the advent of high-throughput technologies measuring high-dimensional biological data, there is a pressing need for visualization tools that reveal the structure and emergent patterns of data in an intuitive form. We present PHATE, a visualization method that captures both local and global nonlinear structure in data by an information-geometric distance between datapoints. We perform extensive comparison between PHATE and other tools on a variety of artificial and biological datasets, and find that it consistently preserves a range of patterns in data including continual progressions, branches, and clusters. We define a manifold preservation metric DEMaP to show that PHATE produces quantitatively better denoised embeddings than existing visualization methods. We show that PHATE is able to gain unique insight from a newly generated scRNA-seq dataset of human germ layer differentiation. Here, PHATE reveals a dynamic picture of the main developmental branches in unparalleled detail, including the identification of three novel subpopulations. Finally, we show that PHATE is applicable to a wide variety of datatypes including mass cytometry, single-cell RNA-sequencing, Hi-C, and gut microbiome data, where it can generate interpretable insights into the underlying systems.

## 1 Introduction

High dimensional, high-throughput data are accumulating at a staggering rate, especially in biological systems measured using single-cell transcriptomics and other genomic and epigenetic assays. Since humans are visual learners, it is vitally important that these datasets are presented to researchers in intuitive ways to understand both the overall shape and the fine granular structure of the data. This is especially important in biological systems where structure exists at many different scales and is often unknown, for example in phenotypic or response progressions, in which a faithful visualization can lead to hypothesis generation.

There are many dimensionality reduction methods for visualization [1–11], of which the most commonly used are PCA [11] and t-SNE [1–3]. However, these methods suffer from several drawbacks that render them suboptimal for exploration of high-dimensional high-throughput biological data. First, they tend to be **sensitive to noise.** Biomedical data is generally very noisy, and methods like PCA and Isomap [4] fail to explicitly remove this noise for visualization, leading to the smearing of local neighborhoods, rendering fine grained local structure impossible to recognize. Second, nonlinear visualization methods often **scramble the global structure in data.** For example, to faithfully represent local neighborhoods, methods such as t-SNE minimize an objective function that explicitly deprioritizes long-range distances, causing the global structure of the final embedding to be quasi-random. Third, many of them **fail to optimize for two-dimensional visualization.** Dimensionality reduction methods that are not specifically designed for visualization (e.g., PCA and diffusion maps) do not explicitly adapt their embedding (or coordinate mapping) to the number of dimensions required. Therefore, the visualized dimensions (e.g., the first two extracted coordinates) often only represent a small fraction of the structure of the entire dataset while discarding important information captured in additional dimensions.

Furthermore, common implementations of dimensionality reduction methods often **lack computational scalability.** The volume of biomedical data being generated is growing at a scale that far outpaces Moore’s Law. State-of-the-art methods such as MDS and t-SNE were originally presented (e.g., in [1,7]) as proofs-of-concept with somewhat naïve implementations that do not scale well to datasets with hundreds of thousands, let alone millions, of data points due to speed or memory constraints. While some heuristic improvements may be made (see, for example, [3,8]), most available packages still follow the original implementation and thus cannot run on big data, which severely limits the usability of these methods in the medium to long term.

Finally, we note that some methods try to alleviate visualization challenges by directly **imposing a fixed geometry or intrinsic structure on the data.** It is relatively simple to generate a faithful data visualization if one knows in advance the expected structure of the data. However, methods that impose a structure on the data generally have no way of alerting the user whether their predetermined structural assumption is correct. Indeed, any data will be transformed to fit a tree with Monocle2 [12] or clusters with t-SNE, for example. Therefore, while such methods are useful for data that fit their prior assumptions, they tend to generate spurious and misleading results otherwise, and are often ill suited for hypothesis generation, or data exploration that seeks emerging structure in data.

To address the above concerns, we have designed a novel dimensionality reduction method for visualization named Potential of Heat-diffusion for Affinity-based Transition Embedding (PHATE). PHATE generates a low-dimensional embedding specific for visualization which provides an accurate, denoised representation of both local and global structure of a dataset in the required number of dimensions, without imposing any strong assumptions on the structure of the data, and is highly scalable, both in memory and runtime.

To achieve this, we introduce novel mathematical contributions combining ideas from manifold learning, information geometry, and data-driven diffusion geometry. We leverage these together with strengths of current state-of-the-art methods, to result in a dimensionality reduction method carefully optimized for visualization of high-dimensional data. PHATE models the data as a statistical manifold in which each data point is represented by a probability distribution that is constructed via data diffusion using a novel graph construction. The process of data diffusion denoises the data. PHATE then compresses this graph for computational efficiency, and preserves a novel informational distance that we call the *potential distance* between these probability distributions that capture the intrinsic geometry of the dataset. The result is that high-dimensional and nonlinear structures, such as clusters, nonlinear progressions, and branches, become apparent in two or three dimensions and can be extracted for further analysis (Figure 1A).

**Figure 1.**
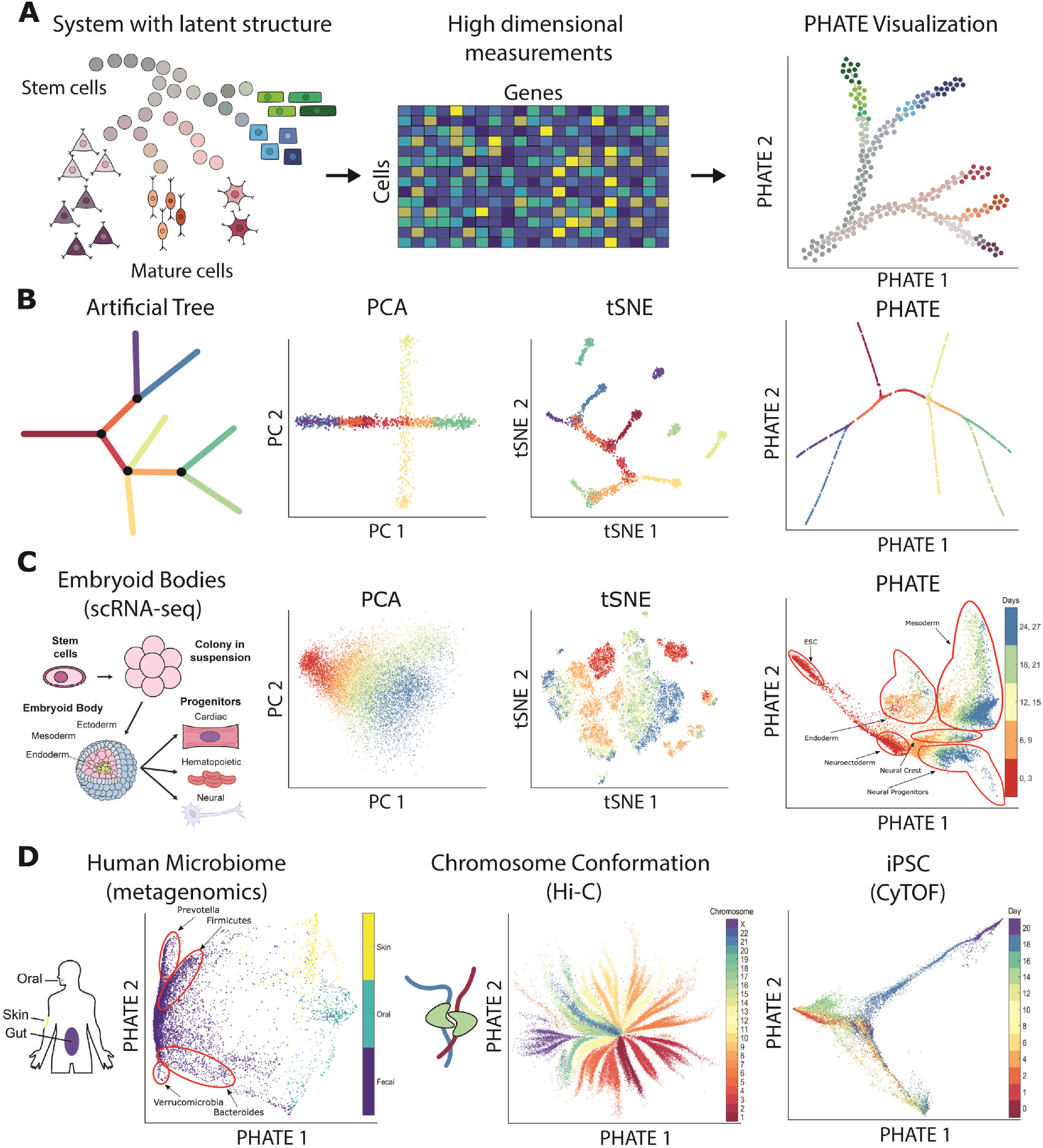
Overview of PHATE and its ability to reveal structure in data. (**A**) Conceptual figure demonstrating the progression of stem cells into different cell types and the corresponding high dimensional single-cell measurements rendered as a visualization by PHATE. (**B**) (Left) A 2D drawing of an artificial tree with color-coded branches. Data is uniformly sampled from each branch in 60 dimensions with Gaussian noise added (see Methods). (Right) Comparison of PCA, t-SNE, and the PHATE visualizations for the high-dimensional artificial tree data. PHATE is best at revealing global and branching structure in the data. In particular, PCA cannot reveal fine-grained local features such as branches while t-SNE breaks the structure apart and shuffles the broken pieces within the visualization. See Figure S3 for more comparisons on artificial data. (**C**) Comparison of PCA, t-SNE, and the PHATE visualizations for new embryoid body data showing similar trends as in (B). (**D**) PHATE applied to various datatypes. Left: PHATE on human microbiome data shows clear distinctions between skin, oral and fecal samples, as well as different enterotypes within the fecal samples. Middle: PHATE on Hi-C chromatin conformation data shows the global structure of chromatin. The embedding is colored by the different chromosomes. Right: PHATE on induced pluripotent stem cell (iPSC) CyTOF data. The embedding is colored by time after induction. See Figures 5, S7, S8, and S9 for more applications to real data.

We show that PHATE consistently outperforms state-of-the-art methods both qualitatively and quantitatively on a wide variety of benchmark test cases where the ground truth is known. We present Denoised Embedding Manifold Preservation (DEMaP), a metric quantifying the ability of an embedding to preserved denoised manifold distances, and show that PHATE consistently outperforms 11 other methods on synthetically generated data with known ground truth. We also use PHATE to visualize several biological and non-biological real world datasets, showing PHATE’s capacity to visualize datasets with many different underlying structures, including trajectories, clusters, disconnected and intersecting manifolds, and more (Figure 1). To demonstrate the ability of PHATE to reveal new biological insights, we apply PHATE to a newly generated single-cell RNA-sequencing dataset of human embryonic stem cells grown as embryoid bodies over a period of 27 days to observe differentiation into diverse cell lineages. PHATE successfully captures all known branches of development within this system as well as numerous novel differentiation pathways, and enables the isolation of rare populations based on surface markers, which we validate experimentally.

## 2 The PHATE Algorithm

Visualizing complex, high-dimensional data in a way that is both easy to understand and faithful to the data is a difficult task. Such a method of visualization needs to preserve local and global structure in the high-dimensional data, denoise the data such that the underlying structure is clearly visible, and preserve as much information as possible in low (2-3) dimensions. In addition to these properties, a visualization method should be robust in the sense that the revealed structure of the data are insensitive to user configurations of the algorithm and scalable to the large sizes of modern data.

Popular dimensionality reduction methods are deficient in one or more of these attributes. For example, t-SNE [1] provides a visualization that focuses on preserving local structure, often at the expense of the global structure (Figure 1B-C). In contrast, PCA focuses on preserving global structure at the expense of the local structure (Figure 1B-C). While PCA is often used for denoising as a preprocessing step, both PCA and t-SNE provide noisy visualizations when the data is noisy, which can obscure the structure of the data (Figure 1B-C). In contrast, diffusion maps [13] effectively denoises data and learns the local and global structure. However, diffusion maps typically encodes this information in higher dimensions [14], which is not amenable to visualization, and can introduce distortions in the visualization under certain conditions (see Figures S1 and S2A). A discussion of other dimensionality reduction methods for visualization is included in Section 2.2.

PHATE is carefully designed to overcome these weaknesses and provide a visualization that preserves the local and global structure of the data, denoises the data, and presents as much information as possible into low dimensions. There are three major steps in the PHATE algorithm.

1. **Encode local data information via local similarities (Figure 2A-C).** Local relationships between data points, even in the presence of noise, are meaningful with respect to the overall structure of the data as they can be chained together to learn global relationships along the manifold. For some data types, such as Hi-C chromatin conformation maps [15], the local relationships are encoded directly in the measurements. In this case, this information can be used directly to encode the local similarities of the data. However, for most data types, the local similarities must be learned. We assume that component-wise, the data are well-modeled as lying on a manifold. Effectively, this means that small Euclidean distances are meaningful while large Euclidean distances may not be. We apply a novel kernel we developed (called the *α*-decay kernel) to Euclidean distances to accurately encode the local structure of the data even when the data is not uniformly sampled along the underlying manifold structure.
2. **Learn global structure and denoise data via diffusion (Figure 2D).** Diffusing through data is a concept that was popularized in the derivation of Diffusion Maps (DM) [13]. Diffusion is performed by first transforming the local similarities into probabilities that measure the probability of transitioning from one data point to another in a single step of a random walk and then powering this operator to *t* steps to give *t*-step walk probabilities. Two points that have a high similarity have a high probability of transition and vice versa. Then, the local single-step transitions are propagated by taking a multi-step random walk. Thus both the local and global structure is encoded in the newly-calculated multi-step transition probabilities, referred to as the diffusion probabilities. For example, two points that have multiple potential, short paths that connect them will have a higher diffusion probability than two points that either have only long paths or relatively few paths connecting them. By considering all possible random walks, the diffusion process also denoises the data by downweighting spurious paths created by noise. Thus at the end of this step each point *x_i_* is a vector of probabilities (in the ith row of the diffused operator) that a random walk starting at *x_i_* would end up at every other vertex within *t* steps.
3. **Embed diffusion probability information into low dimensions for visualization (Figure 2E-F).** Directly embedding the diffusion probabilities into 2 or 3 dimensions via eigenvalue decomposition or distance-preserving methods such as multidimensional scaling (MDS) results in either a loss of information (Figure S1) or an unstable embedding (Figures S2A and S3D), respectively. Since the data are represented at this point by probability distributions, they lie on a statistical manifold. To extract the information from the diffusion probabilities for embedding, we develop a novel informational distance between points on this statistical manifold. We call this distance the *potential distance* (Figure 2E). This information distance is more sensitive to global relationships (between far-away points) and more stable at boundaries of manifolds than straight point-wise comparisons of probabilities (i.e., diffusion distances) as the potential distance is more sensitive to differences between low probabilities, resulting in a stronger emphasis on long-range distances.

**Figure 2.**
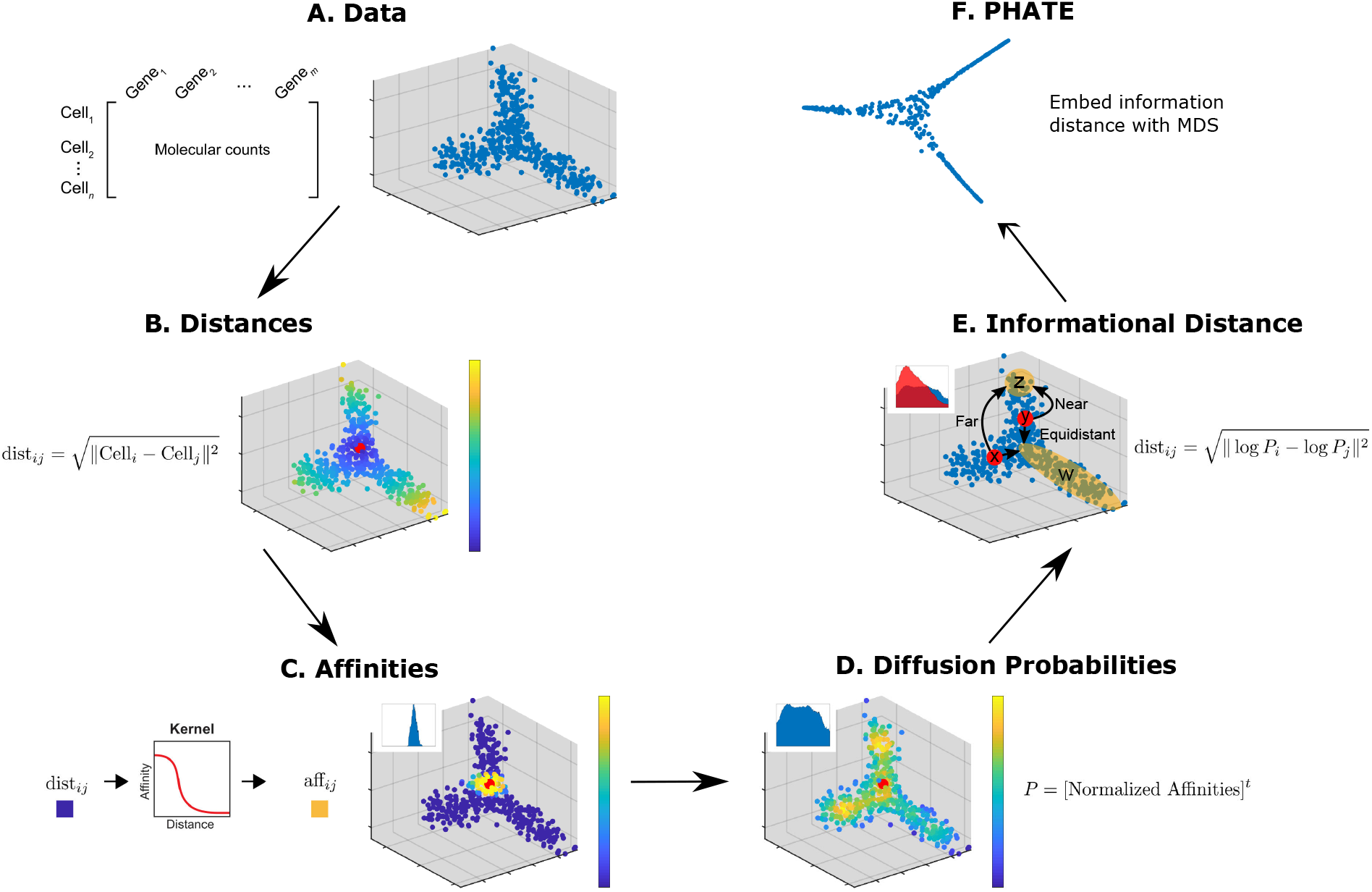
Steps of the PHATE algorithm. (**A**) Data. (**B**) Euclidean distances. Data points are colored by their Euclidean distance to the highlighted point. (**C**) Markov-normalized affinity matrix. Distances are transformed to local affinities via a kernel function and then normalized to a probability distribution. Data points are colored by the probability of transitioning from the highlighted point in a single step random walk. (**D**) Diffusion probabilities. The normalized affinities are diffused to denoise the data and learn long-range relationships between points. Data points are colored by the probability of transitioning from the highlighted point in a *t* step random walk. (**E**) Informational distance. An informational distance (e.g. the potential distance) that measures the dissimilarity between the diffused probabilities is computed. The informational distance is better suited for computing differences between probabilities than the Euclidean distance. See the text for a discussion. (**F**) The final PHATE embedding. The informational distances are embedded into low dimensions using MDS. Note that distances or affinities can be directly input to the appropriate step in cases of connectivity data. Therefore, the Euclidean distance or our constructed affinities can be replaced with distances or affinities that best describe the data. For example, in Figure S9D we replace our affinity matrix with the Facebook connectivity matrix.

To give a more intuitive view, consider two points *x* and *y* that are on different sides of a line of points *W* = {*w*_1_, *w*_2_,…,*w_n_*} (See Figure 2E), suppose that there is a small set of distant points *Z* = {*z*_1_,*z*_2_,…, *z_n_*} that are on the same side of *W* as *y* but opposite side as *x* such that they are twice as far from *x* as from *y*. The representation of each point *x* is as its *t*-step diffusion probability to all other points. So to compute the potential distance between *x* and *y* we compare these probabilities. What is the right type of distance to measure the distinction between these two probability distributions? One solution has been the diffusion distance which is simply the Euclidean distance between these probability distributions. However, in the example mentioned above the diffusion distance would be dominated by larger probabilities and the probabilities to the *Z* points would not affect the distance from *x* to *y* perhaps making them seem close. But instead, we take a divergence between the probabilities from *x* and *y* by first log-scale transforming the probabilities and then taking their Euclidean distance, which makes the distance sensitive to fold-change. Thus, if a probability of 0.01 from *x* to a point *z_i_* is changed to 0.02 from *y* then this has the same effect as if the probabilities had been 0.1 and 0.2. Thus, PHATE is sensitive to small differences in probability distribution corresponding to differences in long-range global structure, which allows PHATE to preserve global manifold relationships using this potential distance.

The information in these distances is then squeezed into low dimensions for visualization via MDS, which creates an embedding by matching the distances in the low-dimensional space to the input distances, which are the potential distances in this case. See Appendix A.1.1 for more details on the advantages of the potential distance.

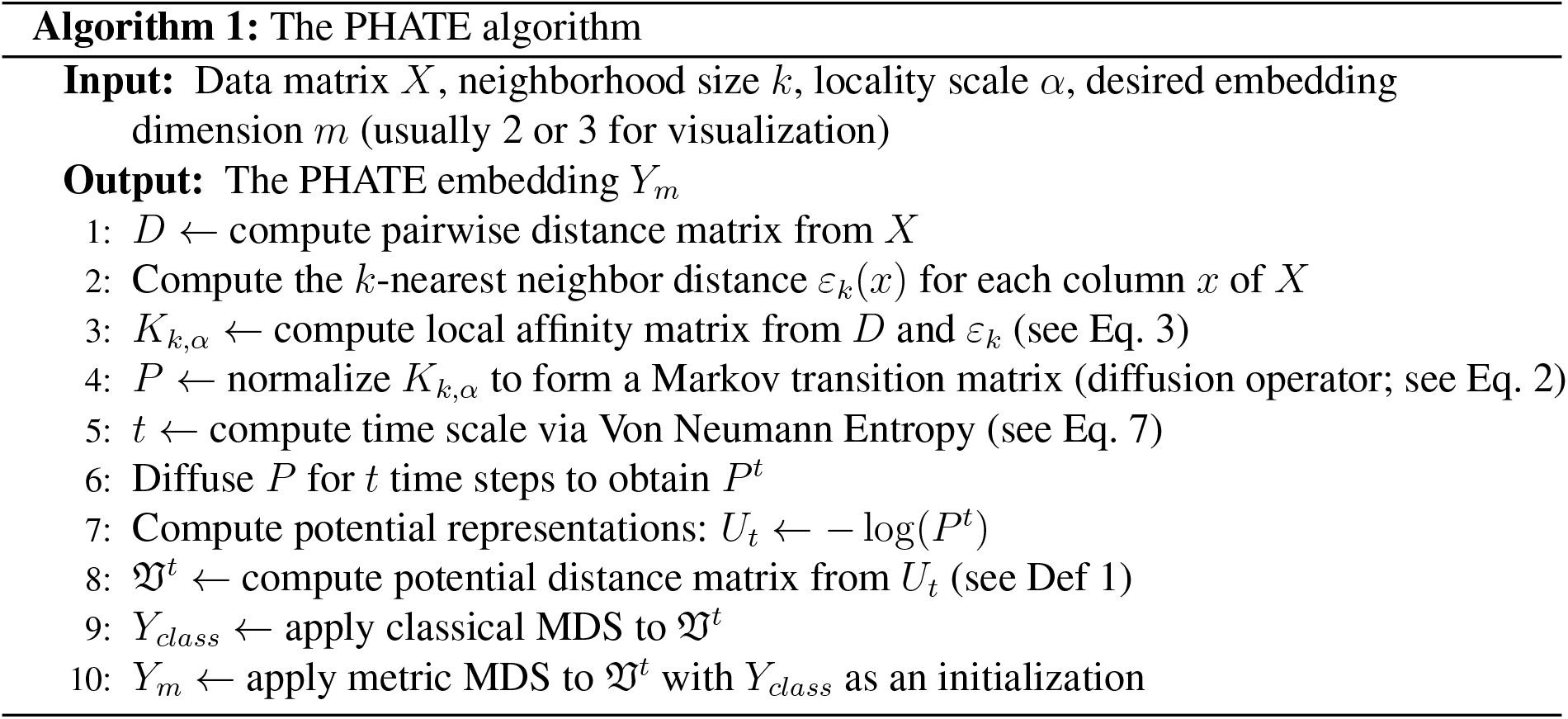

All of these steps are necessary to create a good visualization that preserves local and global structure in the high-dimensional data, denoise the data, and present as much information as possible into low dimensions. Focusing primarily on local relationships can distort global relationships in the data, as evidenced by t-SNE. Focusing primarily on global relationships can distort local relationships, as evidenced by PCA. Additionally, directly embedding both local and global information can result in a loss of information or an unstable embedding, as evidenced by diffusion maps and direct embedding of the diffusion distances. Thus all three steps are necessary. Further details on all of the steps of PHATE are included in Appendix A.1.1 and Algorithm 1. PHATE is also robust to the choice of parameters (Appendix A.1.2 and Figure S4).

In addition to the exact computation of PHATE, we developed a fast version of PHATE that produces near-identical results. In this version, PHATE is implemented in an efficient and scalable manner by using landmark subsampling, sparse matrices, and randomized matrix decompositions. For more details on the scalability of PHATE see Appendix A.1.3, Algorithm 2, and Figure S5, which shows the fast runtime of PHATE on datasets of different sizes, including a dataset of 1.3 million cells (2.5 hours) and a network of 1.8 million nodes (12 minutes).

### 2.1 Extracting Information from PHATE

PHATE embeddings contain a large amount of information on the structure of the data, namely, local transitions, progressions, branches or splits in progressions, and end states of progression. In this section, we present new methods that provide suggested end points, branch points, and branches based on the information from higher dimensional PHATE embeddings. These may not always correspond to real decision points, but provide an annotation to aid the user in interpreting the PHATE visual.

#### Branch Point Identification with Local Intrinsic Dimensionality

Since PHATE emphasizes progressions, PHATE plots can show branch points or divergences in progression. In biological data, branch points often encapsulate switch-like decisions where cells sharply veer towards one of a small number of fates. For example, Figure S6A shows PHATE on a CyTOF dataset of induced pluripotent stem cells (iPSC) [16], with a central branch point identified. This branch point connects the early stages of cells with a branch of cells that are successfully reprogrammed and a branch of cells that are refractory (identified via selected markers including those in Figure S6B) and marks a major decision point between these two cell fates. Identifying branch points in biological data is of critical importance for analyzing such decisions.

We make a key observation that most points in PHATE plots of biological data lie on lowdimensional progressions with some noise as demonstrated in Figure 3Aii. Since branch points lie at the intersections of such progressions, they have higher local intrinsic dimensionality. We can also regard intrinsic dimensionality in terms of degrees of freedom in the progression modeled by PHATE. If there is only one fate possible for a cell (i.e. a cell lies on a branch as in Figure 3Aii) then there are only two directions of transition between data points—forward or backward—and the local intrinsic dimension is low. If on the other hand, there are multiple fates possible, then there are at least three directions of transition possible—a single direction backwards and at least two forward. This cannot be captured by a one dimensional curve and will require a higher dimensional structure such as a plane, as shown in Figure 3Aii. Thus, we can use the concept of local intrinsic dimensionality for identifying branch points.

**Figure 3.**
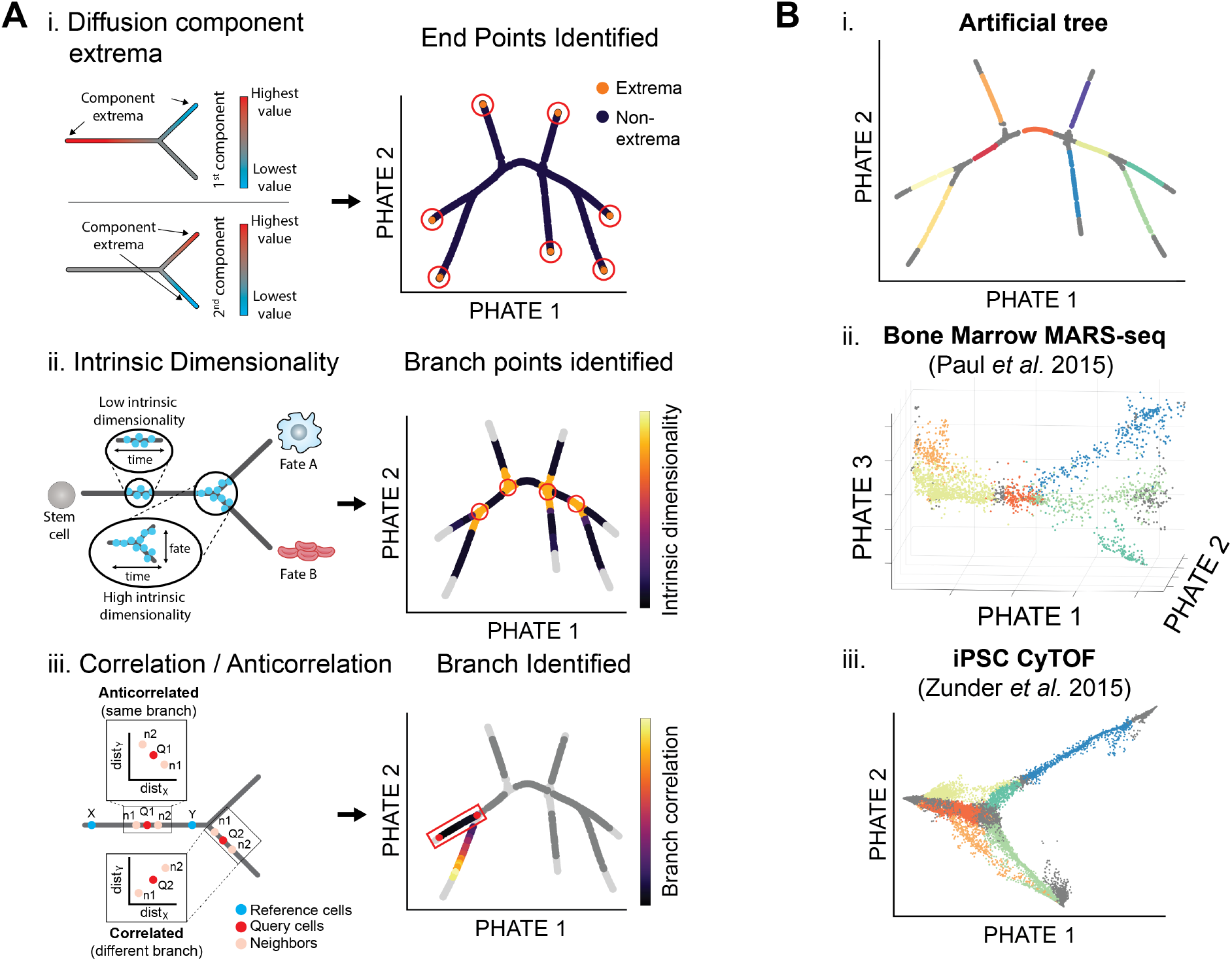
Extracting branches and branchpoints from PHATE. (**A**) Methods for identifying suggested endpoints, branch points, and branches. (i) PHATE computes a specialized diffusion operator as an intermediate step (Figure 2D). We use this diffusion operator to find endpoints. Specifically we use the the extrema of the corresponding diffusion components (eigenvectors of the diffusion operator) to identify endpoints [19]. (ii) Local intrinsic dimensionality is used to find branchpoints in a PHATE visual. As there are more degrees of freedom at branch points, the local intrinsic dimension is higher than through the rest of a branch. (iii) Cells in the PHATE embedding can be assigned to branches by considering the correlation between distances of neighbors to reference cells (e.g. branch points or endpoints). (**B**) Detected branches in the (i) artificial tree data, (ii) bone marrow scRNA-seq data from [21], and (iii) iPSC CyTOF data from [16].

We use a *k*-nn based method for estimating local intrinsic dimensionality [17]. This method uses the relationship between the radius and volume of a *d*-dimensional ball. The volume increases exponentially with the dimensionality of the data. So as the radius increases by *δ*, the volume increases by *δ^d^* where *d* is the dimensionality of the data. Thus the intrinsic dimension can be estimated via the growth rate of a *k*-nn ball with radius equal to the *k*-nn distance of a point. For more details on this approach, see Appendix A.1.4. We note that other local intrinsic dimension estimation methods could be used such as the maximum likelihood estimator in [18]. Figure 3Aii shows that points of intersection in the artificial tree data indeed have higher local intrinsic dimensionality than points on branches.

#### Endpoint Identification

We also identify endpoints in the PHATE embedding. These points can correspond to the beginning or end-states of differentiation processes. For example, Figure S6A shows the PHATE visualization of the iPSC CyTOF dataset from [16] with highlighted endpoints, or end-states, of the reprogrammed and refractory branches. While many major endpoints can be identified by inspecting the PHATE visualization, we provide a method for identifying other endpoints or end-states that may be present in the higher dimensional PHATE embedding. We identify these states using data point centrality and distinctness as described below.

First, we compute the centrality of a data point by quantifying the impact of its removal on the connectivity of the graph representation of the data (as defined using the local affinity matrix *K_k,α_*). Removing a point that is on a one dimensional progression pathway, either branching point or not, breaks the graph into multiple parts and reduces the overall connectivity. However, removing an endpoint does not result in any breaks in the graph. Therefore we expect endpoints to have low centrality, as estimated using the eigenvector centrality measure of *K_k,α_*.

Second, we quantify the distinctness of a cellular state relative to the general data. We expect the beginning or end-states of differentiation processes to have the most distinctive cellular profiles. As shown in [19] we quantify this distinctness by considering the minima and the maxima of diffusion eigenvectors (see Figure 3Ai). Thus we identify endpoints in the embedding as those that are most distinct and least central.

#### Branch Identification

After identifying branch points and endpoints, the remaining points can be assigned to branches between two branch points or between a branch point and endpoint. Due to the smoothly-varying nature of centrality and local intrinsic dimension, the previously described procedures identify regions of points as branch points or endpoints rather than individual points. However, it can be useful to reduce these regions to representative points for analysis such as branch detection and cell ordering. To do this, we reduce these regions to representative points using a “shake and bake” procedure similar to that in [20]. This approach groups collections of branch points or endpoints together into representative points based on their proximity (see Appendix A.1.4 for details). Further, a representative point is labeled an endpoint if the corresponding collection of points contains one or more endpoints as identified using centrality and distinctness. Otherwise, the representative point is labeled a branch point.

After representative points have been selected, the remaining points can be assigned to corresponding branches. We use an approach based on the branch point detection method in [14] that compares the correlation and anticorrelation of neighborhood distances. However, we use higher dimensional PHATE coordinates instead of the diffusion maps coordinates. Figure 3Aiii gives a visual demonstration of this approach and details are given in Appendix A.1.4. Figure 3B shows the results of our approach to identifying branch points, endpoints, and branches on an artificial tree dataset, a scRNA-seq dataset of the bone marrow [21], and an iPSC CyTOF dataset [16]. Our procedure identifies the branches on the artificial tree perfectly and defines biologically meaningful branches on the other two datasets which we will use for data exploration.

### 2.2 Comparison of PHATE to Other Methods

Here we compare PHATE to multiple dimensionality reduction methods. We provide quantitative comparisons on simulated data where the ground truth is known, and provide a qualitative comparison using both simulated and real biological data.

#### Quantitative Comparisons

To compare PHATE to other visualization methods quantitatively, we formulated the Denoised Embedding Manifold Preservation (DEMaP) metric. DEMaP is designed to encapsulate the desirable properties of a dimensionality reduction method that is intended for visualization. These include: 1. the preservation of relationships in the data such that cells close together on the manifold are close together in the embedded space; and cells that are far apart on the manifold are far apart in the embedding, including disconnected manifolds (e.g. clusters) which should be as well separated as possible; and 2. denoising, such that the low-dimensional embedding accurately represents the ground truth data and is as invariant as possible to biological and technical noise. DEMaP encapsulates each of these properties by comparing the geodesic distances on the noiseless data to the Euclidean distances of the embedding extracted from noisy data. An overview of DEMaP is presented in Figure 4A. See Appendix A.3.1 for details.

**Figure 4.**
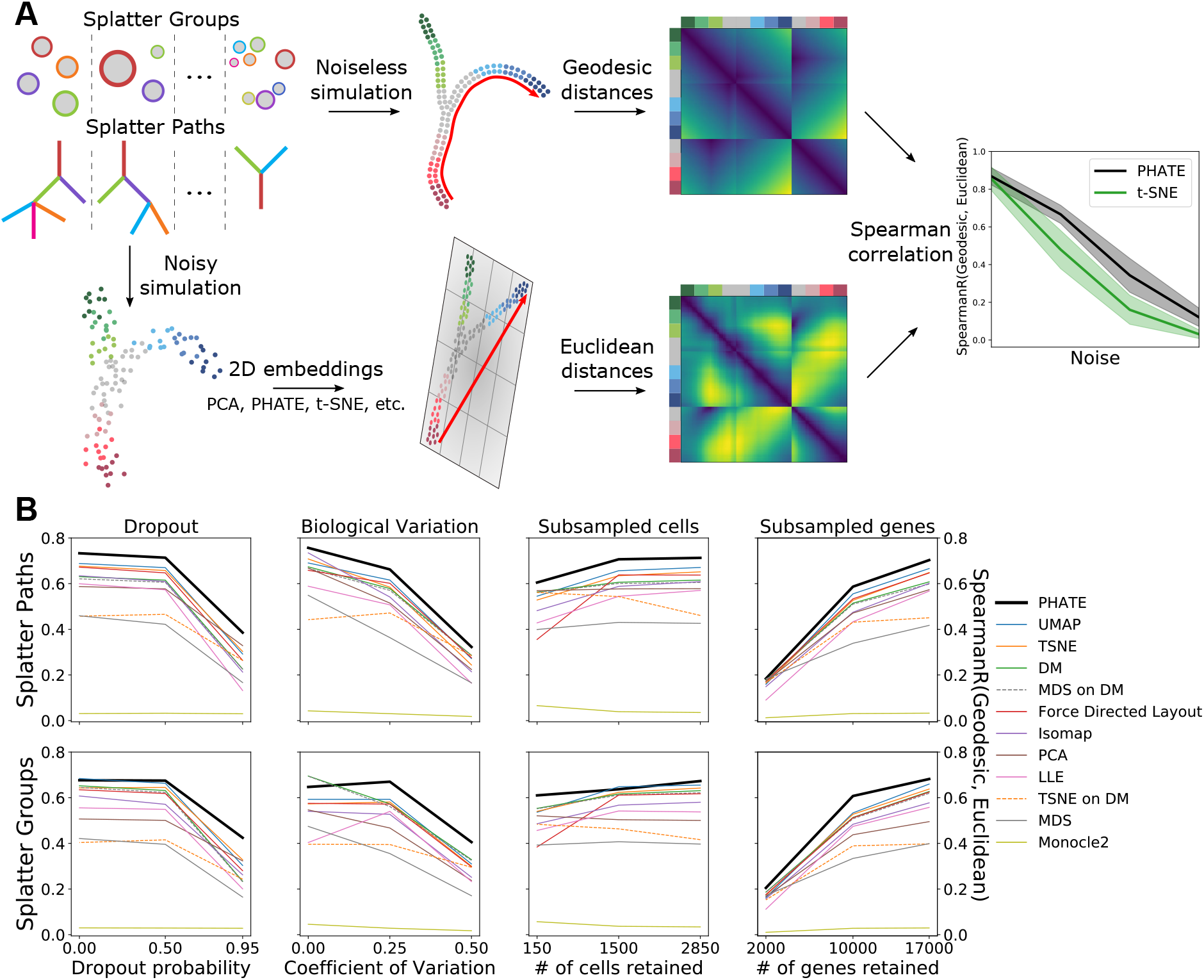
PHATE most accurately represents manifold distances in a 2D embedding. (**A**) Schematic description of performance comparison procedure. For each method and each type of corruption, Euclidean distances in the 2D embedding are compared to geodesic distances in an equivalent noiseless simulation by Spearman correlation. (**B**) Performance of 12 different methods across varying levels of corruption by dropout, decreased signal-to-noise ratio (BCV), randomly subsampled cells (subsample) and randomly subsampled genes (n_genes). Mean correlation of 20 runs for each configuration is shown. For further details see Table 1.

To compare the performance of PHATE to other dimensionality reduction methods, we calculated DEMaP using simulated scRNA-seq data using Splatter [22]. Splatter uses a parametric model to generate data with various structures, such as branches or clusters. This simulated data provides a ground truth reference to which we can add various types of noise. We then use this noisy data as input for each dimensionality reduction algorithm, and quantify the degree to which each representation preserves local and global structures and denoises the data.

To generate a diverse set of ground truth references, we simulated 50 datasets containing clusters and 50 datasets containing branches. In each of these simulated datasets, the number and size of the clusters of branches as well as the global position of the clusters or branches with respect to each other is random. Furthermore, the local relationships between individual cells on these structures is random. Finally, the changes in gene expression within clusters or along branches is random. The output of this simulation is the ground truth reference.

Next, we add biological and technical noise to the reference data. First, to simulate stochastic gene expression we use Splatter’s Biological Coefficient of Variation (BCV) parameter, which controls the level of gene expression in each cell following an inverse gamma distribution. Second, to simulate the inefficient capture of mRNA in single cells, we undersample from the true counts using the default BCV. Third, to demonstrate robustness to varying of total genes measured, we randomly remove genes from the data matrix. Finally, to demonstrate robustness to the number of cells captured, we randomly remove cells from each dataset. We vary each of these parameters, including by default some degree of biological variation and mRNA undersampling to each simulation. See Appendix A.3.3 for the exact parameters used for each simulation.

We compared DEMaP for 12 visualization methods including t-SNE and UMAP. For each method, we used the default parameters and calculated visualization accuracy on each simulated dataset using 12 regimes of biological and technical noise parameters. We then averaged the accuracy across the 50 cluster datasets and 50 branch datasets for a total of 24 comparisons. The results are presented in Figure 4B and Table 1. We found that PHATE had the highest DEMaP score in 22/24 comparisons and was the top-performing method overall. UMAP was the second best performing method overall but had the highest DEMaP score in only two of the comparisons, one of which is equal with PHATE. From these results, we conclude that PHATE captures the true structure of high dimensional data more accurately than existing visualization methods.

**Table 1:**
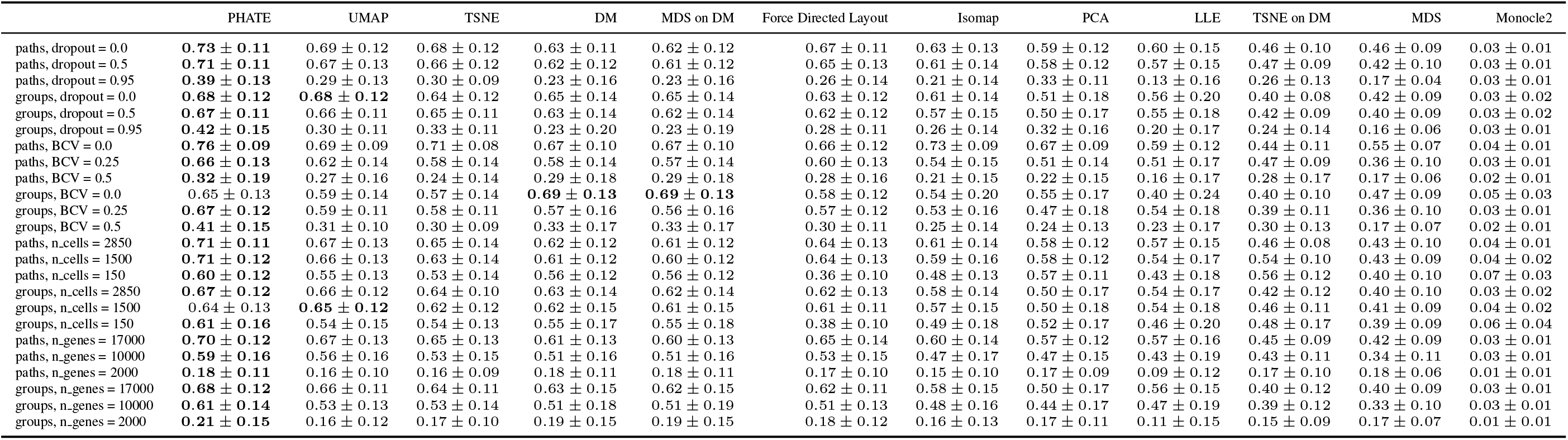
Spearman correlation between Euclidean distances in 2D dimensionality reductions of Splatter simulations and geodesic distances calculated on ground truth data under different noise settings. The best performing method for each test is shown in bold. Results are presented in the form of mean correlation coefficient ± standard deviation. For each category, results are ordered by noise level from least to most. Methods are ordered by mean correlation across all tests, from left to right. The subsample parameter indicates the proportion of cells retained. The n_genes parameter indicates the number of retained genes.

#### Qualitative Comparisons

In addition to the quantitative comparison, we can visually compare the embeddings provided by different methods. Figure 5 shows a comparison of the PHATE visualization to seven other methods on five single-cell datasets with known trajectory (Figure 5A,D,E) and cluster (Figure 5B-C) structures. We see that PHATE provides a clean and relatively denoised visualization of the data that highlights both the local and global structure: local clusters or branches are visually connected to each other in a global structure in each of the PHATE visualizations. Many of these branches are consistent with cell types or clusters validated by the authors [16,21,23,24] and additionally are also present in other visualizations such as force directed layout or t-SNE, suggesting that these structures in the PHATE embedding reflect true structure in the dataset. However, force directed layout tends to give a noisier visualization with fewer clear branches. Additionally, t-SNE [25] tends to shatter trajectories into clusters. Thus t-SNE creates the false impression that data contain natural clusters, which could lead to incorrect analysis. PCA tends to preserve the global structure well, but it gives a noisy visualization with very little visible local structure. We characterize each of these visualizations in detail in Appendix B.

**Figure 5.**
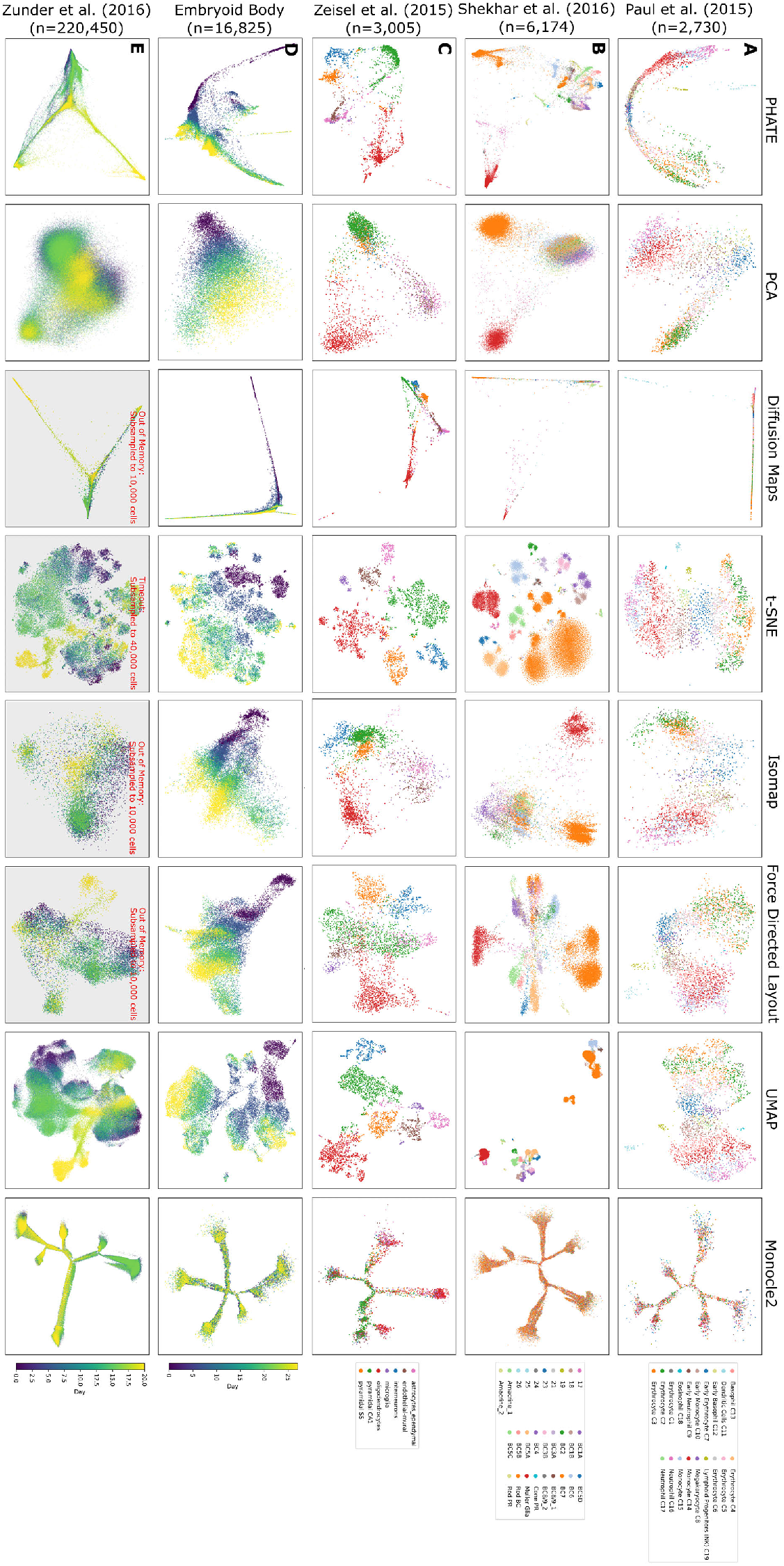
Comparison of PHATE to other visualization methods on biological datasets. Columns represent different visualization methods, rows different datasets.

We obtained similar results by comparing PHATE to eleven methods on nine non-biological datasets, including four artificial datasets where the ground truth is known (Figure S3). Expanded comparisons on single-cell data, including additional datasets and visualization methods, are also included in Figure S7. See Appendix B for a full discussion of each method in all of these comparisons. A discussion of desirable attributes of some of these methods for the purposes of dimensionality reduction can also be found in [25].

## 3 Data Exploration with PHATE

PHATE can reveal the underlying structure of the data for a variety of datatypes. Appendix C discusses PHATE applied to multiple different datasets, including SNP data, microbiome data, Facebook network data, Hi-C chromatin conformation data, and facial images (Figures S8 and S9). In this section, however, we show the insights gained through the PHATE visualization of this structure for single-cell data. See Appendix A.2.2 for details on preprocessing steps.

We show that the identifiable trajectories in the PHATE visualization have biological meaning that can be discerned from the gene expression patterns and the mutual information between gene expression and the ordering of cells along the trajectories. We analyze the mouse bone marrow scRNA-seq [21] and iPSC CyTOF [16] datasets described previously. Our analysis of the iPSC CyTOF data is presented here while the analysis of the mouse bone marrow data is presented in Appendix C. For both of these datasets, we used our new methods for detecting branches and branch points. We then ordered the cells within each trajectory using Wanderlust [26] applied to higher-dimensional PHATE coordinates. Ordering is generally from left to right. We note that ordering could also be based on other pseudotime ordering software such as those in [14,27–30]. To estimate the strength of the relationship between gene expression and cell ordering along branches, we estimated the DREMI score (a conditional-density resampled mutual information that eliminates biases to reveal shape-agnostic relationships between two variables [31]) between gene expression and the Wanderlust-based ordering within each branch. Genes with a high DREMI score within a branch are changing along the branch. We also use PHATE to analyze the transcriptional heterogeneity in rod bipolar cells to demonstrate PHATE’s ability to preserve cluster structure.

### iPSC Mass Cytometry Data

Figure S6C shows the mass cytometry dataset from [16] that shows cellular reprogramming with Oct4 GFP from mouse embryonic fibroblasts (MEFs) to induced pluripotent stem cells (iPSCs) at the single-cell resolution. The protein markers measure pluripotency, differentiation, cell-cycle and signaling status. The cellular embedding (with combined timepoints) by PHATE shows a unified embedding that contains five main branches, further segmented in our visualization, each corresponding to biology identified in [16]. Branch 2 contains early reprogramming intermediates with the correct set of reprogramming factors Sox2^+^/Oct4^+^/Klf4^+^/Nanog^+^ and with relatively low CD73 at the beginning of the branch. Branch 2 splits into two additional branches. Branches 4 and 6 (Figure S6) show the successful reprogramming to ESC-like lineages expressing markers such as Nanog, Oct4, Lin28 and Ssea1, and Epcam that are associated with transition to pluripotency [32]. Branch 5 shows a lineage that is refractory to reprogramming, does not express pluripotency markers, and is referred to as “mesoderm-like” in [16].

Then, branch 3 represents an intermediate, partially reprogrammed state also containing Oct4^+^/Klf4^+^/CD73^+^ but is not yet expressing pluripotency markers like Nanog or Lin28. However, the PHATE embedding indicates that Epcam, which is known to promote reprogramming generally [33], increases along this branch. This is evidenced by the high DREMI score between Epcam and the cell ordering within the branch (Figure S6C). This branch joins into branch 4 at a later stage, showing perhaps an alternative path or timing of reprogramming. Finally, branch 1 shows a lineage that has failed to reprogram, perhaps due to the wrong stoichiometry of the reprogramming factors [34]. Of note, this lineage contains low Klf4 which is an essential reprogramming factor.

Additionally, the PHATE embedding shows a decrease in p53 expression in precursor branches (2 and 3) indicating that these cells are released from cell cycle arrest induced by initial reprogramming factor over expression [35]. However, along the refractory branch (branch 5) we see an increase in cleaved-caspase3, potentially indicating that the failure to reprogram correctly initiates apoptosis in these cells [16].

### PHATE on different views of the data

By default PHATE produces a single low dimensional embedding of a dataset. However, we can obtain variants of this embedding by reweighting the features before computing distances. Such reweightings correspond to specific “views” of the data. For example, in a biological context, we can upweight genes that are involved in a specific process to have PHATE prominently reflect this process. To demonstrate this reweighting scheme, we computed three alternative PHATE embeddings of the iPSC data, by upweighting either cell cycle markers, stem cell markers, or mitotic markers (Figure S10). PHATE, after upweighting cell cycle markers, gives an embedding with a circular structure (Figure S10Aii) that reflects the cyclical nature of the cell cycle. In addition to the circular structure, the embedding shows a small protrusion, with high expression of Ccasp3, suggesting that these cells are apoptotic. Upweighting stem cell markers gives an embedding with a 1-dimensional progression. Expression analysis reveals that stem cell markers such as Sox2 are high at one end of the progression and low on the other end. Moreover, the progression is correlated with time (measurement day), further supporting the idea that the progression that PHATE reveals marks the extent to which the cells are stem-like, with early timepoints being less stem-like. Finally, after upweighting mitotic markers, PHATE shows a different 1-dimensional progression. Here, the progression appears to be correlated with mitotic state, as can be seen by the expression of several mitosis-related genes (Figure S10Biii), such as pAKT, that are high only in one end of the embedding. Thus, PHATE computed after reweighting the genes can be used to obtain a process specific embedding to gain insight into predefined biological processes.

### PHATE reveals transcriptional heterogeneity in Rod Bipolar Cells

Figure S11Ai shows PHATE on scRNA-seq data of mouse retinal bipolar neurons from [23]. Cells were collected from an adult mouse and sorted for transgenic retinal bipolar markers. PHATE visualizes cluster structure while preserving relationships between clusters. The embedding is colored by the clusters described in the original study, which seeks to transcriptionally characterize all subtypes of bipolar cells. In the original characterization, rods bipolar cells (the largest cluster of cells) are shown as a single homogeneous cell type. However, the PHATE embedding in Figure S11Ai reveals a bifurcating trajectory within this cluster. We zoomed in on this bifurcating trajectory, by embedding just those cells, in order to determine their sub-structure.

This embedding of just rod bipolar cells (Figure S11Aii) reveals four distinct sub-clusters (found by k-means clustering) of rod bipolar cells. We characterize the transcriptional profile of these sub-clusters in Figure S11B, showing all genes used for cell type assignment in [23]. Our results show a trajectory between rod bipolar cell types that is consistent with previous work by [36], in which cell types RB1 and RB2 are shown to be a continuum of variants of a single type. Further, we show distinct differences between these in known marker genes distinguishing RB1 and RB2 *Trnp1, Rho* and *Pde6b* [37], indicating that the four clusters we observe may be further subtypes of these two cell types.

## 4 Exploratory Analysis with PHATE on Human ESC Differentiation Data

### 4.1 Embryoid Body Single-Cell RNAseq Time Course

To test the ability of PHATE to provide novel insights in a complex biological system, we generated and analyzed scRNA-seq data from human embryonic stem cells (hESCs) differentiating as embryoid bodies (EB) [38], a system which has never before been extensively analyzed at the single-cell level. EB differentiation is thought to recapitulate key aspects of early embryogenesis and has been successfully used as the first step in differentiation protocols for certain types of neurons, astrocytes and oligodendrocytes [39–42], hematopoietic, endothelial and muscle cells [43–51], hepatocytes and pancreatic cells [52,53], as well as germ cells [54,55]. However, the developmental trajectories through which these early lineage precursors emerge from hESCs as well as their cellular and molecular identities remain largely unknown, particularly in human models.

We measured approximately 31,000 cells, equally distributed over a 27-day differentiation time course (Figure S12A and Appendix A.2.1). Samples were collected at 3-day intervals and pooled for measurement on the 10x Chromium platform. The PHATE embedding of the EB data revealed a highly ordered and clean cellular structure dominated by continuous progressions (Figures 1C and 6A), unlike other methods such as PCA or t-SNE (Figure S7). Exploratory analysis of this system using PHATE uncovered a comprehensive map of four major germ layers with both known and novel differentiation intermediates which were not captured with other visualization methods.

**Figure 6.**
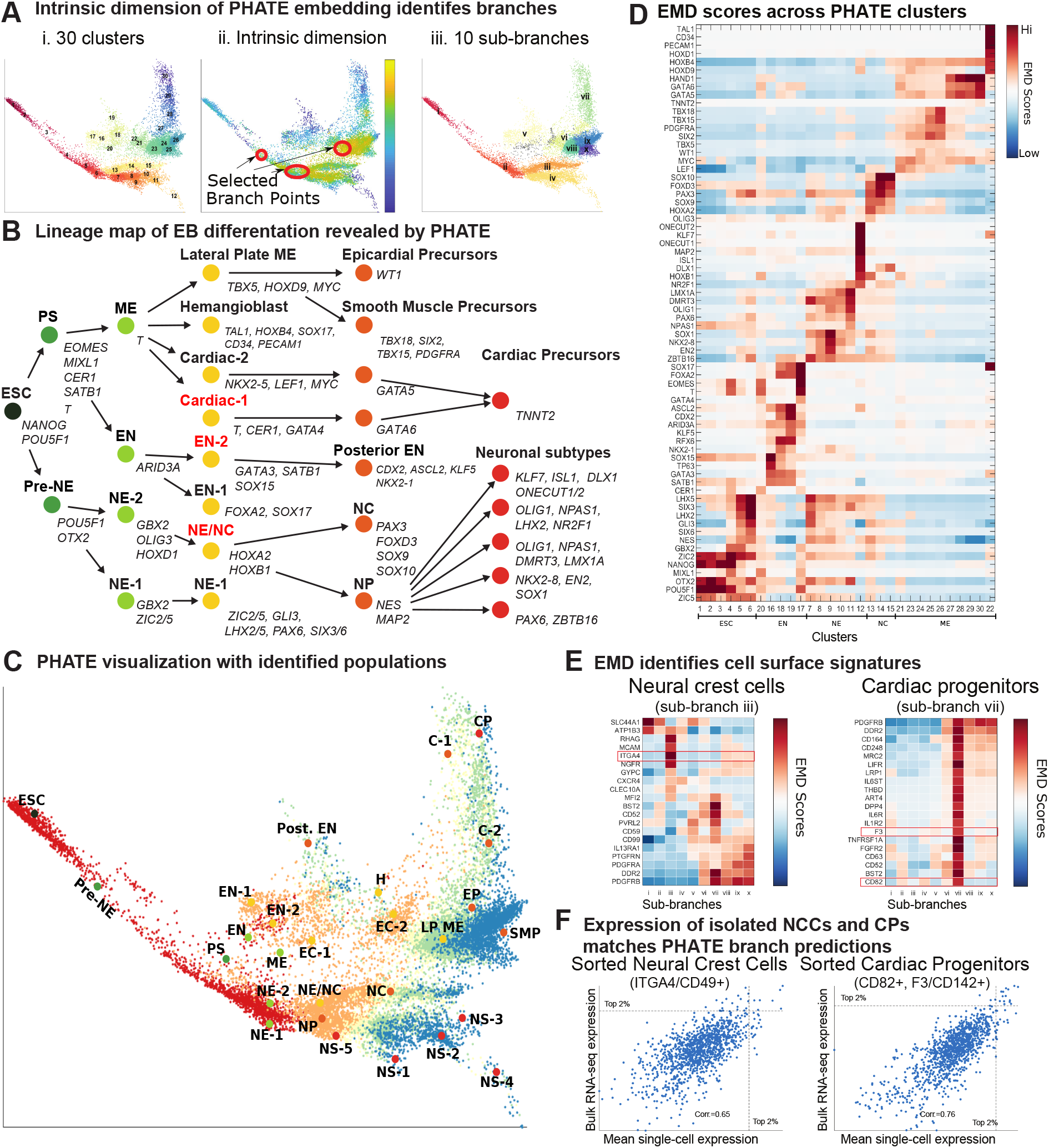
PHATE analysis of embryoid body scRNA-seq data. (**A**) i) The PHATE visualization colored by clusters. Clustering is done on a ten dimensional PHATE embedding. ii) The PHATE visualization colored by estimated local intrinsic dimensionality with selected branch points highlighted. iii) Branches and subbranches chosen from contiguous clusters for analysis.(**B**) Lineage tree of the EB system determined from the PHATE analysis showing embryonic stem cells (ESC), the primitive streak (PS), mesoderm (ME), endoderm (EN), neuroectoderm (NE), neural crest (NC), neural progenitors (NP), and others. Red font indicates novel cell precursors. (**C**) PHATE embedding overlaid with each of the populations in the lineage tree. Other abbreviations include lateral plate ME (LP ME), hemangioblast (H), cardiac (C), epicardial precursors (EP), smooth muscle precursors (SMP), cardiac precursors (CP), and neuronal subtypes (NS). (**D**) Heatmap showing the EMD score between the cluster distribution and the background distribution for each gene. Relevant genes for identifying the main lineages were manually identified. Genes are organized according to their maximum EMD score. (**E**) The EMD scores of the top scoring surface markers in the targeted sub-branches (sub-branches iii and vii). (**F**) Scatter plots of the bulk transcription factor expression vs. the mean single-cell transcription factor expression in sub-branches iii (left) and vii (right).

### 4.2 Deriving a Comprehensive Lineage Map with PHATE

Importantly, PHATE retained global structure and organization of the data as is evidenced by the retention of a strong time trend in the embedding, although sample time was not included in creating the embedding. Further, PHATE revealed greater phenotypic diversity at later time points as seen by the larger space encompassed by the embedding at days 18 to 27 (Figure 1C).

This phenotypic heterogeneity was further analyzed by both an automated analysis described in Sections 2.1 (for identified clusters and branch points, see Appendix D, Figure 6A and Table S1) and by manual examination of the embedding in conjunction with the established literature on germ layer development (Figure S12B). For the manual analyses, we used 80 markers from the literature to identify populations along the PHATE map which gave rise to a detailed germ layer specification map (Figure 6B). These populations are shown on the PHATE visualization in Figure 6C. In the lineage tree, the dots are the populations and the arrows represent transitions between the populations. Our map shows in detail how hESCs give rise to germ layer derivatives via a continuum of defined intermediate states.

### 4.3 Identification of Novel Transitional Populations with PHATE

The comprehensive nature of the lineage map generated from the PHATE embedding allowed us to identify novel transitional populations that have not yet been characterized. Three novel pre-cursor states were identified in both manual and automated analyses: a bi-potent NC and NP pre-cursor, a novel EN precursor, and a novel cardiac precursor.

Within the ectodermal lineage, differentiation begins with the induction of pre-NE state characterized by downregulaton of POU5F1 and induction of OTX2. This state is resolved into two precursors, NE-1 (*GBX2+ZIC2/5+*) and NE-2 (*GBX2+OLIG2+HOXD1+*). While NE-1 neuroectoderm appeared to develop along the canonical NE specification route and expressed a set of well established anterior NE markers (*ZIC2/5, PAX6, GLI3, SIX3/6*), the NE-2 neuroectoderm gave rise to a bi-potent *HOXA2+HOXB1+* precursor that subsequently separated into the NC branch and neural progenitor (NP) branch. Given its potential to generate both NE and NC cell types, the *HOXA2+HOXB1+* precursor could represent the equivalent of the neural plate border cells that have been defined in model organisms [56, 57].

Within the EN branch, the canonical *EOMES+FOXA2+SOX17+* EN precursor was clustered together with the novel *EOMES-FOXA2-GATA3+SATB1+KLF8+* precursor, which further differentiated into cells expressing posterior EN markers *NKX2-1, CDX2, ASCL2*, and *KLF5*. Finally, a novel *T+GATA4+ CER1+PROX1+* cardiac precursor cell was identified within the ME lineage that gave rise to *TNNT2+* cells via a *GATA6+HAND1+* differentiation intermediate.

A more detailed analysis of the novel and canonical cell types derived from the PHATE embedding is given in Appendix D.

### 4.4 Experimental Validation of PHATE-Identified Lineages

We next used the ability of PHATE to extract data on specific regions within the visualization to define a set of surface markers for the isolation and molecular characterization of specific cell populations within the EB differentiation process.

We focused on two specific regions that correspond to the NC branch (sub-branch iii, Figure 6Aiii) and cardiac precursor sub-branch within the ME branch (sub-branch vii, Figure 6Aiii). Differential expression analysis identified a set of candidate markers for each region (Figures 6D-E). We focused on markers with a high Earth Mover’s Distance (EMD) [58] score in the targeted sub-branch, and low EMD scores in all other sub-branches (see Appendix A.1.5 for more details on the EMD). Based on these analyses and the availability of antibodies, *CD49D/ITGA4* was chosen for the neural crest (the highest scoring surface marker for subbranch iii) while *CD142/F3* and *CD82* were chosen for cardiac precursors (among the top 6% of surface markers and the top 3% of all genes by EMD). We FACS-purified *CD49d+CD63-* and *CD82+CD142+* and performed bulk RNA-sequencing (Figure S12F) on these sorted populations.

To verify that we isolated the correct regions, we calculated the Spearman correlation between the gene expression pattern of each cell and the bulk RNA-seq data from the CD49d+CD63-sorted cells (Figures 6F and S12D). The correlation coefficient was the highest in the neural crest branch (branch iii), which corresponds to the highest expression of CD49d. Similar results were obtained for the cardiac precursor cells (Figures 6F and S12E).

Taken together, our analyses show that PHATE has the potential to greatly accelerate the pace of biological discovery by suggesting hypotheses in the form of finely grained populations and identifying markers with which to isolate populations. These populations can be probed further using alternative measurements such as epigenetic or protein expression assays.

## 5 Conclusion

With large amounts of high-dimensional high-throughput biological data being generated in many types of biological systems, there is a growing need for interpretable visualizations that can represent structures in data without strong prior assumptions. However, most existing methods are highly deficient at retaining structures of interest in biology. These include clusters, trajectories or progressions of various dimensionality, hybrids of the two, as well as local and global nonlinear relations in data. Furthermore, existing methods have trouble contending with the sizes of modern datasets and the high degree of noise inherent to biological datasets. PHATE provides a unique solution to these problems by creating a diffusion-based informational geometry from the data, and by preserving a divergence metric between datapoints that is sensitive to near and far manifold-intrinsic distances in the dataspace. Additionally, PHATE is able to offer clean and denoised visualizations because the information geometry created in PHATE is based on data diffusion dynamics which are robust to noise. Thus, PHATE reveals intricate local as well as global structure in a denoised way.

We applied PHATE to a wide variety of datasets, including single-cell CyTOF and RNA-seq data, as well as Gut Microbiome and SNP data, where the datapoints are subjects rather than cells. We also tested PHATE on network data, such as Hi-C and Facebook networks. In each case, PHATE was able to reveal structures of visual interest to humans that other methods entirely miss. Moreover, we have implemented PHATE in a scalable way that enables it to process millions of datapoints in a matter of hours. Hence, PHATE can efficiently handle the datasets that are now being produced using single-cell RNA sequencing technologies, as we demonstrated using the 1.3 million cell dataset released by 10X genomics.

To showcase the ability of PHATE to explore data generated in new systems, we applied PHATE to our newly generated human EB differentiation dataset consisting of roughly 30,000 cells sampled over a differentiation time course. We found that PHATE successfully resolves cellular heterogeneity and correctly maps all germ layer lineages and branches based on scRNA-seq data alone, without any additional assumptions on the data. Through detailed sub-population and gene expression analysis along these branches we identified both canonical and novel differentiation intermediates. The insights obtained with PHATE in this system will be a valuable resource for researchers working on early human development, human ES cells, and their regenerative medicine applications.

We expect numerous biological, but also non-biological, data types to benefit from PHATE, including applications in high-throughput genomics, phenotyping, and many other fields. As such, we believe that PHATE will revolutionize biomedical data exploration by offering a new way of visualizing, exploring and extracting information from large scale high-dimensional data.

## A Methods

Here we present an expanded explanation of our computational methods, experimental methods, and data processing steps.

### A.1 Computational Methods

Here we further discuss the computational methods. We first provide a detailed overview of the PHATE algorithm followed by a robustness analysis of PHATE with respect to the parameters and the number of datapoints. We then provide details on the scalable version of PHATE, identifying branch points and branches, and the EMD score analysis.

#### A.1.1 PHATE Overview

A common way of visualizing high dimensional data is by using dimensionality reduction methods, which map high dimensional data points to low dimensional coordinates (i.e., two or three dimensions in this case) by minimizing some notion of distortion. For example, PCA aims to preserve variance, and thus its notion of distortion is derived from variance lost by linear projection [11]. Similarly, diffusion maps preserve diffusion affinities as inner products [13,59], and Isomap preserves geodesic distances as Euclidean distances [4]. The minimized notion of distortion in these methods can be derived directly from the appropriate inner products or distances, and related to their algorithmic steps. Finally, the popular t-SNE method [1] aims to directly preserve neighborhood structure, and uses KL divergence between high dimensional Gaussian neighborhoods and low dimensional neighborhoods (captured via Student t-distributions) as its distortion.

The effectiveness of a given dimensionality reduction method in a particular application should be considered by ensuring its distortion notion, or resulting embedding, faithfully preserve structures and patterns of interest. Here, we focus on clustering and progression (e.g., trajectory) structures, due to their established significance and importance in biological data. It should be noted that many, if not most, of the common dimensionality reduction methods were not originally designed for visualization purposes, but rather for alleviating the curse of dimensionality [60,61]. While they can be used for visualization by simply setting the dimensionality of the embedding to be two or three, there is no guarantee that the resulting visualization will reliably express clusters, trajectories, or other patterns of interest.

In this work we focus on embedding data into a two dimensional space for visualization purposes. The embedding provided by PHATE is specifically designed to enable visualization of global and local structure in exploratory settings with the following criteria in mind:

**Visualization:** To enable visualization, PHATE captures variance in low (2-3) dimensions.
**Manifold-structure preserving:** To provide an interpretable view of dynamics (e.g., pathways or progressions) in the data, PHATE preserves and emphasizes global nonlinear transitions in the data, in addition to local transitions.
**Denoising:** To enable unsupervised data exploration, PHATE denoises data such that progressions within the data are immediately identifiable and clearly separated.
**Robust:** PHATE produces a robust embedding in the sense that the revealed boundaries and the intersections of progressions within the data are insensitive to user configurations of the algorithm.

PHATE fulfills these criteria with an abstract geometric model based primarily on two properties that we typically observe in high throughput data (biomedical and otherwise). First, transitions between data points tend to be incremental and gradual. There may be many such patches of incremental change but nevertheless these gradual transitions are usually prevalent. Second, there are a limited number of intrinsic directions (or pathways) along which datapoints progress. Therefore, the dynamics captured by collected data are inherently more like a set of rivers, rather than a cloud (expanding outwards in all directions).

Data with such properties can thus be modeled geometrically by a collection of smoothly varying data patches defined by local neighborhoods [62]. This collection essentially fits the manifold learning paradigm, which relies on a mathematical manifold model for the geometry of a progression track, together with analysis tools for characterizing it. Furthermore, data manifolds often have a low intrinsic dimension, even if curvature and noise forces them to span a high dimensional ambient volume in the collected feature space. Finally, progression tracks form trajectories, with a limited number of “branching points”, where progression splits into several directions. Therefore, in this case the underlying data geometry can implicitly be regarded as a collection of intrinsically low-dimensional manifolds (i.e., curves, surfaces) that cross each other in branching points. By exploiting this low-dimensional structure, we avoid the effects of the curse of dimensionality.

It has been shown in several works (e.g., [63,64]) that manifold geometries are closely related to heat diffusion, which is modeled by the heat equation – a differential equation defined in terms of the Laplace-Beltrami operators. Indeed, meta-stable solutions of the heat equation over a manifold capture its intrinsic properties, while providing embeddings, affinities, and distance metrics that capture intrinsic manifold relations. It has further been shown that these can be robustly discretized for empirical observations that correlate with hidden (or latent) manifold models, e.g., by considering diffusion maps embedding of the data [13,65,66]. The embedding obtained by PHATE extends these results by considering this diffusion goemetry as a statistical manifold of diffusion distributions and using tools of information geometry (namely, *α*-representations) to capture its metric stucture and embed it in visualizable (i.e., two or three) dimensions. Further, as we discuss in the following sections, the information distance metric we use also relates to Boltzman energy potentials of the diffusion process, and therefore it combines together both the dyamical systems and information geometry aspects of data-driven diffusion geometries. In particular, for the case of transition structures, this approach enables the consideration of the underlying data geometry consisting of multiple low-dimensional manifolds (such as trajectory curves) that cross each other, while alleviating boundary-condition instabilities to maintain low dimensionality of the embedded space that is better-suited for visualization.

We note that the trajectory structure is not artificially generated in our case, but rather it is expected to be dominant (albeit latent or hidden) in the data. Therefore, the PHATE visualization will only show trajectory structures when data fits such a geometry; otherwise, other (e.g., cluster) patterns will be expressed in the PHATE visualization.

Here we provide further details about each of the steps in PHATE and we explain how each of these steps help us ensure that the provided embedding satisfies the four properties described above.

##### Distances

Consider the common approach of linearly embedding the raw data matrix itself, e.g., with PCA, to preserve the global structure of the data. PCA finds the directions of the data that capture the largest global variance. However, in most cases local transitions are noisy and global transitions are nonlinear. Therefore, linear notions such as global variance maximization are insufficient to capture latent patterns in the data, and they typically result in a noisy visualization (Figure S3, Column 2). To provide reliable *structure preservation* that emphasizes transitions in the data, we need to consider the *intrinsic* structure of the data. This implies and motivates preserving distances between data points (e.g., cells) that consider gradual changes between them along these nonlinear transitions (Figure 2A-B).

##### Affinities and the Diffusion Operator

A standard choice of a distance metric is the Euclidean distance. However, global Euclidean distances are not reflective of transitions in the data, especially in biological datasets that have nonlinear and noisy structures. For instance, cells sampled from a developmental system, such as hematopoiesis or embryonic stem cell differentiation, show gradual changes where adjacent cells are only slightly different from each other. But these changes quickly aggregate into nonlinear transitions in marker expression along each developmental path. Therefore, we transform the global Euclidean distances into local affinities that quantify the similarities between nearby (in the Euclidean space) data points (Figure 2C).

Let 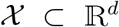 be a dataset with *N* points sampled i.i.d. from a probability distribution *p*: ℝ^*d*^ → [0, ∞) (with ∫ *p*(*x*)*dx* = 1) that is essentially supported on a low dimensional manifold 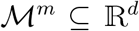, where *m* is the dimension of 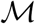 and *m* ≪ *d*. A common approach to transforming global distances to local similarities is to apply some kernel function. A popular kernel function is the Gaussian kernel *k_ε_*(*x,y*) = exp(−||*x* − *y*||^2^/*ε*) that quantifies similarities between points based on Euclidean distances. The bandwidth ε determines the radius (or spread) of neighborhoods captured by this kernel.

Embedding local affinities directly can result in a loss of global structure as is evident in t-SNE (Figures 1, 5, S7, and S3) or kernel PCA embeddings. For example, t-SNE only preserves data clusters, but not transitions between clusters, since it does not enforce any preservation of global structure. In contrast, a faithful structure-preserving embedding (and visualization) needs to go beyond local affinities (or distances), and consider more global relations between parts of the data. To accomplish this, PHATE is based on constructing a diffusion geometry to learn and represent the shape of the data [13,65,66]. This construction is based on computing local similarities between data points, and then *walking* or *diffusing* through the data using a Markovian random-walk diffusion process to infer more global relations (Figure 2D).

The initial probabilities in this random walk are calculated by normalizing the row-sums of the kernel matrix. In the case of the Gaussian kernel described above, we obtain the following:

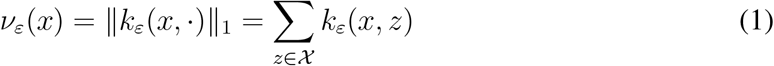

resulting in a *N × N* row-stochastic matrix

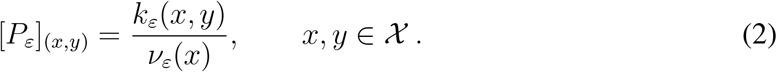

The matrix *P_ε_* is a Markov transition matrix where the probability of moving from *x* to *y* in a single time step is given by Pr[*x* → *y*] = [*P_ε_*]_(*x,y*)_. This matrix is also referred to as the diffusion operator.

##### The **α**-decaying Kernel and Adaptive Bandwidth

When applying the diffusion map framework to data, the choice of the kernel *K* and bandwidth *ε* plays a key role in the results. In particular, choosing the bandwidth corresponds to a tradeoff between encoding global and local information in the probability matrix *P_ε_*. If the bandwidth is small, then single-step transitions in the random walk using *P_ε_* are largely confined to the nearest neighbors of each data point. In biological data, trajectories between major cell types may be relatively sparsely sampled. Thus, if the bandwidth is too small, then the neighbors of points in sparsely sampled regions may be excluded entirely and the trajectory structure in the probability matrix *P_ε_* will not be encoded. Conversely, if the bandwidth is too large, then the resulting probability matrix *P_ε_* loses local information as [*P_ε_*]_(*x*,·)_ becomes more uniform for all 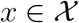, which may result in an inability to resolve different trajectories. Here, we use an adaptive bandwidth that changes with each point to be equal to its kth nearest neighbor distance, along with an **α**-decaying kernel that controls the rate of decay of the kernel.

The original heuristic proposed in [13] suggests setting *ε* to be the smallest distance that still keeps the diffusion process connected. In other words, it is chosen to be the maximal 1-nearest neighbor distance in the dataset. While this approach is useful in some cases, it is greatly affected by outliers and sparse data regions. Furthermore, it relies on a single manifold with constant dimension as the underlying data geometry, which may not be the case when the data is sampled from specific trajectories rather than uniformly from a manifold. Indeed, the intrinsic dimensionality in such cases differs between mid-branch points that mostly capture one-dimensional trajectory geometry, and branching points that capture multiple trajectories crossing each other.

This issue can be mitigated by using a locally adaptive bandwidth that varies based on the local density of the data. A common method for choosing a locally adaptive bandwidth is to use the *k*-nearest neighbor (NN) distance of each point as the bandwidth. A point *x* that is within a densely sampled region will have a small *k*-NN distance. Thus, local information in these regions is still preserved. In contrast, if *x* is on a sparsely sampled trajectory, the *k*-NN distance will be greater and will encode the trajectory structure. We denote the *k*-NN distance of *x* as *ε*_k_(*x*) and the corresponding diffusion operator as *P_k_*.

A weakness of using locally adaptive bandwidths alongside kernels with exponential tails (e.g., the Gaussian kernel) is that the tails become heavier (i.e., decay more slowly) as the bandwidth increases. Thus for a point *x* in a sparsely sampled region where the *k*-NN distance is large, [*P_k_*]_(*x*,·)_ may be close to a fully-supported uniform distribution due to the heavy tails, resulting in a high affinity with many points that are far away. This can be mitigated by using the following kernel

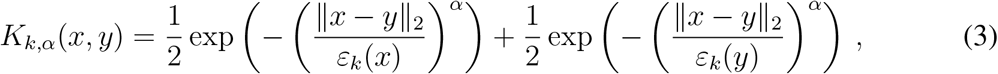

which we call the *α*-decaying kernel. The exponent *α* controls the rate of decay of the tails in the kernel *K_k,α_*. Increasing *α* increases the decay rate while decreasing *α* decreases the decay rate. Since *α* = 2 for the Gaussian kernel, choosing *α* > 2 will result in lighter tails in the kernel *K_k,α_* compared to the Gaussian kernel. We denote the resulting diffusion operator as *P_k,α_*. This is similar to common utilizations of Butterworth filters in signal processing applications [67]. See Figure S2B for a visualization of the effect of different values of *α* on this kernel function.

Our use of a locally adaptive bandwidth and the kernel *K_k,α_* requires the choice of two tuning parameters: *k* and *α*. *k* should be chosen sufficiently small to preserve local information, i.e., to ensure that [*P_k,α_*]_(*x*,·)_ is not a fully-supported uniform distribution. However, *k* should also be chosen sufficiently large to ensure that the underlying graph represented by *P_k,α_* is sufficiently connected, i.e., the probability that we can *walk* from one point to another within the same trajectory in a finite number of steps is nonzero.

The parameter *α* should also be chosen with *k*. *α* should be chosen sufficiently large so that the tails of the kernel *K_k,α_* are not too heavy, especially in sparse regions of the data. However, if *k* is small when *α* is large, then the underlying graph represented by *P_k,α_* may be too sparsely connected, making it difficult to learn long range connections. Thus we recommend that *α* be fixed at a large number (e.g. *α* ≥ 10) and then *k* can be chosen sufficiently large to ensure that points are locally connected. In practice, we find that choosing *k* to be around 5 and *α* to be about 10 works well for all the data sets presented in this work. However, the PHATE embedding is robust to the choice of these parameters as demonstrated in Appendix A.1.2.

In addition to progression or trajectory structures, the recommendations provided in this section work well for visualizing data that naturally separate into distinct clusters. In particular, the *α*-decay kernel ensures that relationships are preserved between distinct clusters that are relatively close to each other.

##### Propagating Affinities via Diffusion

Here we discuss diffusion, i.e., raising the diffusion operator to its *t*-th power as shown in Alg. 1 (Figure 2D). To simplify the discussion we use the notation *P* for the diffusion operator, whether defined with a fixed-bandwidth Gaussian kernel or our adaptive kernel. This matrix is referred to as the diffusion operator, since it defines a Markovian diffusion process that essentially only allows single-step transitions within local data neighborhoods whose sizes depend on the kernel parameters (*ε* or *k* and *α*). In particular, let 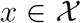 and let *δ_x_* be a Dirac at *x*, i.e., a row vector of length *N* with a one at the entry corresponding to *x* and zeros everywhere else. The *t*-step distribution of *x* is the row in 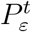 corresponding to *x*:

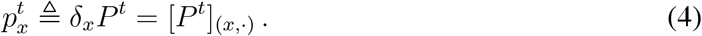

These distributions capture multi-scale (where *t* serves as the scale) local neighborhoods of data points, where locality is considered via random walks that propagate over the intrinsic manifold geometry of the data. This provides a global and robust intrinsic data distance that preserving the overall structure of the data. In addition to learning the global structure, powering the diffusion operator has the effect of low-pass filtering the data such that the main pathways in it are emphasized and small noise dimensions are diminished, thus achieving the denoising objective of our method as well.

For appropriate choices of kernel parameters (as described in previous sections), the diffusion process defined by *P* is ergodic and it thus has a unique stationary distribution *p*^∞^ that is independent of the initial conditions of the process. Thus 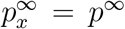 for all 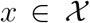. The stationary distribution *p*^∞^ is the left eigenvector of *P* with eigenvalue λ_0_ = 1 and can be written explicitly as *ν*/||*ν*||_1_ with the row-sums from Eq. 1 (possibly adapted to use *K_k,α_* from Eq. 3). It can be shown [66] that for fixed-bandwidth Gaussian-kernel diffusion, *p*^∞^ converges asymptotically to the original distribution *p* of the data as *N* → ∞ and *ε* → 0.

The representation provided by the diffusion distributions 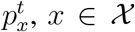, defines a diffusion geometry with the diffusion distance

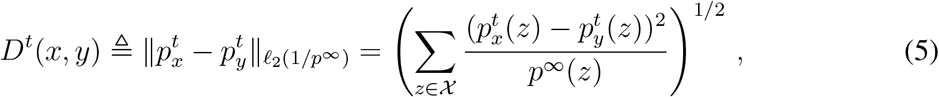

which is given by a weighted *ℓ*_2_ distance between the diffusion distributions originating from the data points *x* and *y*. This distance incorporates a comparison between intrinsic manifold regions of the two data points as well as the concentration of data between them, i.e., the difference between the mass distributions.

The diffusion distance at all time scales can be approximated by the Euclidean distance in the diffusion map embedding, which is defined as follows. If the diffusion process is connected, the eigenvalues of *P* can be indexed as 1 = λ_0_ > λ_1_ ≥ ⋯ λ_*N*−1_ ≥ 0. Let *ψ_i_* and *ϕ_i_* be the corresponding *i*th left and right eigenvectors of *P*, respectively. The diffusion map embedding is defined as

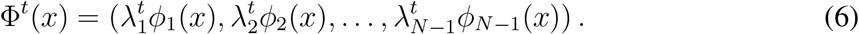

The time scale *t* only impacts the scaling of the embedded coordinates via the powers of the eigenvalues. It can then be shown that *D^t^*(*x, y*) = ||Φ^*t*^(*x*) − Φ^*t*^(*y*)||_2_.

##### Choosing the Diffusion Time Scale *t* with Von Neumann Entropy

The diffusion time scale *t* is an important parameter that affects the embedding. The parameter *t* determines the number of steps taken in a random walk. A larger *t* corresponds to more steps compared to a smaller *t*. Thus, *t* provides a tradeoff between encoding local and global information in the embedding. The diffusion process can also be viewed as a low-pass filter where local noise is smoothed out based on more global structures. The parameter *t* determines the level of smoothing. If *t* is chosen to be too small, then the embedding may be too noisy. On the other hand, if *t* is chosen to be too large, then some of the signal may be smoothed away.

We formulate a new algorithm for choosing the timescale *t*. Our algorithm quantifies the information in the powered diffusion operator with various values of *t*. This is accomplished by computing the spectral or *Von Neumann Entropy* (VNE) [68,69] of the powered diffusion operator. The amount of variability explained by each dimension is equal to its eigenvalue in the eigendecomposition of the related (non-Markov) affinity matrix that is conjugate to the Markov diffusion operator. The VNE is calculated by computing the Shannon entropy on the normalized eigenvalues of this matrix. Due to noise in the data, this value is artificially high for low values of *t*, and rapidly decreases as one powers the matrix. Thus, we choose values that are around the “knee” of this decrease.

More formally, to choose *t*, we first note that its impact on the diffusion geometry can be determined by considering the eigenvalues of the diffusion operator, as the corresponding eigenvectors are not impacted by the time scale. To facilitate spectral considerations and for computational ease, we use a symmetric conjugate

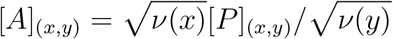

of the diffusion operator *P* with the row-sums *ν*. This symmetric matrix is often called the diffusion affinity matrix. The VNE of this diffusion affinity is used to quantify the amount of variability. It can be verified that the eigenvalues of *A^t^* are the same as those of *P^t^*, and furthermore these eigenvalues are given by the powers 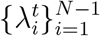 of the spectrum of *P*. Let *η*(*t*) be a probability distribution defined by normalizing these (nonnegative) eigenvalues as 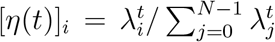. Then, the VNE *H*(*t*) of *A^t^* (and equivalently of *P^t^*) is given by the entropy of *η*(*t*), i.e.,

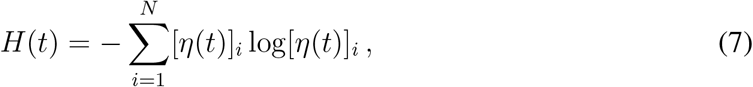

where we use the convention of 0 log(0) ≜ 0. The VNE *H*(*t*) is dominated by the relatively large eigenvalues, while eigenvalues that are relatively small contribute little. Therefore, it provides a measure of the number of the relatively significant eigenvalues.

The VNE generally decreases as *t* increases. As mentioned previously, the initial decrease is primarily due to a denoising of the data as less significant eigenvalues (likely corresponding to noise) decrease rapidly to zero. The more significant eigenvalues (likely corresponding to signal) decrease much more slowly. Thus the overall rate of decrease in *H*(*t*) is high initially as the data is denoised but then low for larger values of *t* as the signal is smoothed. As *t* → ∞, eventually all but the first eigenvalue decrease to zero and so *H*(*t*) → 0.

To choose *t*, we plot *H*(*t*) as a function of *t* as in the first plot of Figure S2C. Choosing *t* from among the values where *H*(*t*) is decreasing rapidly generally results in noisy visualizations and embeddings (second plot in Figure S2C). Very large values of *t* result in a visualization where some of the branches or trajectories are combined together and some of the signal is lost (fourth plot in Figure S2C). Good PHATE visualizations can be obtained by choosing *t* from among the values where the decrease in *H*(*t*) is relatively slow, i.e. the set of values around the “knee” in the plot of *H*(*t*) (third plot in Figure S2C and the PHATE visualizations in Figure 1). This is the set of values for which much of the noise in the data has been smoothed away, and most of the signal is still intact. The PHATE visualization is fairly robust to the choice of *t* in this range, as discussed in Appendix A.1.2.

In the code, we include an automatic method for selecting *t* based on a knee point detection algorithm that finds the knee by fitting two lines to the VNE curve [70]. This algorithm calculates the error between the VNE plot and two lines fitted to the data. The first line has endpoints at the first VNE value and the suggested knee point. The second line has endpoints at the suggested knee point and the last VNE value. The suggested knee point with the minimum error is selected.

##### Potential Distances

To resolve instabilities in diffusion distances and embed the global structure captured by the diffusion geometry in low (2 or 3) dimensions, we instead use a novel diffusion-based informational distance, which we call potential distance (Figure 2E). It is calculated by computing the distance between log-transformed transition probabilities from the powered diffusion operator. The key insight in formulating the potential distance is that an informational distance between probability distributions is more sensitive to global relationships (between far-away points) and more stable at boundaries of manifolds than straight point-wise comparisons of probabilities (i.e., diffusion distances). This is because the diffusion distance is sensitive to differences between the main modes of the diffused probabilities and is largely insensitive to differences in the tails. In contrast, the potential distance, or more generally informational distances, use a submodular function (such as a log) to render distances sensitive to differences in both the main modes and the tails. This gives PHATE the ability to preserve both local and manifold-intrinsic global distances in a way that is optimized for visualization. The resulting metric space also quantifies differences between energy potentials that dominate “heat” propagation along diffusion pathways (i.e., based on the heat-equation diffusion model) between data points, instead of simply considering transition probabilities along them.

The potential distance is inspired by information theory and stochastic dynamics, both fields where probability distributions are compared for different purposes. First, in information theory literature, information divergences are used to measure discrepancies between probability distributions in the information space rather than the probability space, as they are more sensitive to differences between the tails of the distributions as described above. Second, when analyzing dynamical systems of moving particles, it is not the point-wise difference between absolute particle counts that is used to compare states, but rather the ratio between these counts. Indeed, in the latter case the *Boltzmann Distribution Law* directly relates these ratios to differences in the energy of a state in the system. Therefore, similar to the information theory case, dynamical states are differentiated in energy terms, rather than probability terms. We employ the same reasoning in our case by defining our potential distance using localized diffusion energy potentials, rather than diffusion transition probabilities.

To go from the probability space to the energy (or information) space, we log transform the probabilities in the powered diffusion operator and consider an *L*^2^ distance between these localized energy potentials in the data as our intrinsic data distance, which forms an M-divergence between the diffusion probability distributions [71,72].

To provide further mathematical context to the potential distance used in PHATE, we relate it here to the heat propagation dynamics that govern the diffusion geometry and the diffusion process we use to build it. This heat diffusion process can be analyzed by considering two possible scenarios for the origin of the dataset 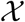 and its distribution *p*, as described in [65,66]. In the first scenario, the data generation process is modeled as an instantiation of a dynamical system that has reached an equilibrium state independent of the initial conditions. Mathematically, let *U*(*x*) be a potential and *w*(*x*) be a d-dimensional Brownian motion process. The data distribution is the steady state solution of the stochastic differential equation (SDE) 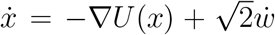, where 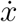 denotes differentiation of *x* with respect to time. The time steps of the system are dominated by the forward and backward Fokker-Planck equations. This steady state solution is given by

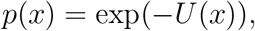

up to normalization in the *L*^1^ norm to form a proper probability distribution.

The distribution of the data in this case is dominated by the potential *U* that models the underlying structure of the data. As an example, if the data is uniformly distributed on or around a manifold, then this potential is minimal on the manifold itself and increases rapidly when deviating from the manifold. The underlying potential also incorporates data densities that are not uniform. For example, data clusters are represented as local wells or pits in the underlying potential, while progression trajectories and transitions between clusters are represented as rivers or branches in the potential. See [65, 66] for more details.

In the second scenario, the data generation process is not modeled as a dynamical system. Instead, we consider the data in this case as generated by drawing *N* i.i.d. samples from the probability distribution *p*(*x*). We then artificially define the underlying potential of the data as

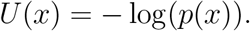

The potential *U* can be used in this scenario since its properties and its relation to the structure of the data are not directly related to the notion of time. Furthermore, in both scenarios, the diffusion-based analysis introduces the notion of diffusion time to reveal intrinsic data geometry. Finally, as shown in [65, 66], in both scenarios the Markov process that defines the diffusion geometry converges asymptotically to a diffusion process governed by Fokker-Planck equations with a potential 2*U*(*x*), whether the original potential is defined naturally or artificially.

Using the same relationship between a potential *U* and an equilibrium distribution *p*, we can define a diffusion potential from the stationary distribution *p*^∞^ as *U*^∞^ = −log(*p*^∞^). This potential corresponds to data generation using the random walk process defined by *P_ε_* with *t* → ∞ with random initial conditions. Similarly, if we consider a data generation process using this random walk process with *t*-steps and a fixed initial condition *δ_x_*, then the generated data is distributed according to 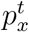 and the corresponding *t*-step potential representation of *x* is 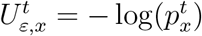.

Given the potential representations 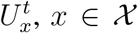 of the data in 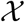, we define the following potential distance metric as an alternative to the distribution-based diffusion distance:

###### Definition 1.

*The t*-step **potential distance** *is defined as* 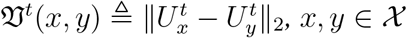.

The following proposition shows a relation between the two metrics by expressing the potential distance in embedded diffusion map coordinates^1^ for fixed-bandwidth Gaussian-based diffusion (i.e., generated by *P_ε_* from Eq. 2):

###### Proposition 1.

*Given a diffusion process defined by a fixed-bandwidth Gaussian kernel, the potential distance from Def 1 can be written as* 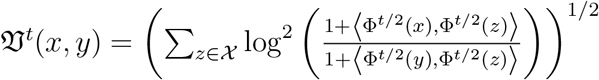

*Proof*. According to the spectral theorem, the entries of 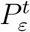 can be written as

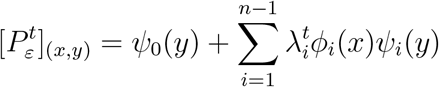

since powers of the operator *P_ε_* only affect the eigenvalues, which are taken to the same power, and since the trivial eigenvalue λ_0_ is one and the corresponding right eigenvector *ϕ*_0_ only consists of ones. Furthermore, it can be verified that the left and right eigenvectors of *P_ε_* are related by *ψ_i_*(*y*) = *ϕ_i_*(*y*)*ψ*_0_(*y*), thus, combined with Eqs. 4 and 6, we get

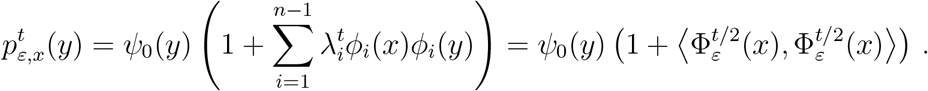

By applying the logarithm to both ends of this equation we express the entries of the potential representation 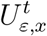 as

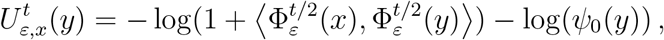

and thus for any *j* = 1,…, *N*,

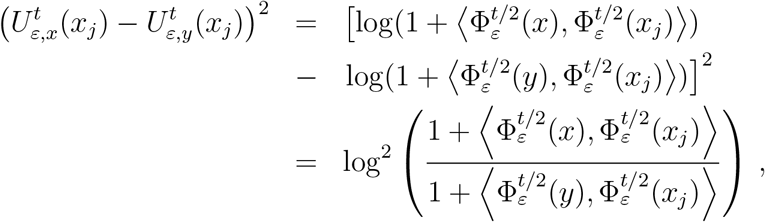

which yields the result in the proposition.

##### Alternative Informational Distances

We note that the potential distance is a particular case of a wider family of diffusion-based informational distances that view the diffusion geometry as a statistical manifold in information geometry. We offer a generalization of these to a family of distances, where the diffusion distance (Eq. 5) is at one extreme of this family and the potential distance (Def. 1) is at the other. Diffusion distances directly compare probability distributions pointwise with no *damping* of high probabilities. Thus changes in the tails of distributions do not contribute much to the distance. The potential distance is damped using the log function to the point where fold changes, even in low probabilities, have impact on the distance. Other distances, including the Hellinger distance [73], are in between these two. We introduce a parameter *γ* as a knob to control the level of damping, with the diffusion distance having *γ* = −1, the potential distance with *γ* =1 and the Hellinger distance at *γ* = 0.

Mathematically, let the diffusion distributions 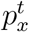 and 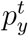 represent data points *x*, 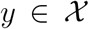. Given an intermediate data point 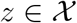, and a meta-parameter −1 ≤ *γ* ≤ 1, let

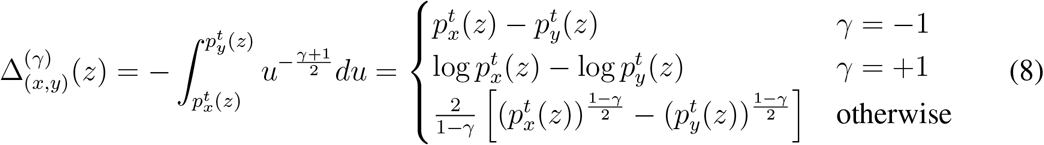

quantify the difference between the transition probabilities 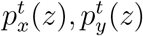 from *x,y* to *z*. Then, diffusion distances and potential distances are given by *L*^2^ norms of 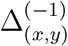 and 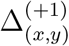 correspondingly (albeit with a different measure over *z*, since diffusion distances are defined over 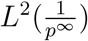 in Eq. 5). Therefore, these distances can be regarded as two extremes of a general family of distances over the diffusion geometry. Moreover, as we show in Prop 2, diffusion dissimilarities of the form 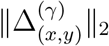 combine both this diffusion-geometry notion of a data-driven distance metric, and an information-theory notion of divergence between diffusion distributions, which is provided in Def. 2 for completeness.

###### Definition 2

(Divergence [Information Theory]). *Let S be a space of probability distributions with common support. A divergence on S is a function D*(·||·): *S × S* → ℝ *s.t. (1) D*(*p||q*) ≥ 0 *for all p,q* ∈ *S, and (2) D*(*p||q*) = 0 *if and only if p* = *q* [74]. *Some specific classes of divergences include*:

i. M-divergence [71,72]: *Let p and q be probability mass functions and let g be a differentiable function with continuous second derivative. Then the M-divergence between p and q is M_g_*(*p, q*) = Σ_*i*_(*g*(*p_i_*) − *g*(*q_i_*))^2^.
ii. *f*-divergence [75,76]: *Let f be a convex function s.t. f*(1) = 0 *and let p and q be probability mass functions. The f-divergence between p and q is* 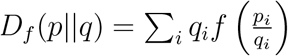.
iii. Bregman divergence [77]: *Let* Ω *be a closed convex set and let F*: Ω → ℝ *be a strictly convex and continuously differentiable function. The Bregman divergence between the points p, q* ∈ Ω *is D_F_*(*p, q*) = *F*(*p*) − *F*(*q*) − 〈∇*F*(*q*),*p* − *q*〉, *where* ∇*F*(*q*) *is the gradient of F evaluated at q*.

Unlike (formal) distance metrics, a divergence is not required to be symmetric nor satisfy the triangle inequality. Examples of *f*-divergences include the Kullback-Leibler divergence [78] and the Hellinger distance [73]. An example of a Bregman divergence is the squared Euclidean distance where *F*(*x*) = *x*^2^. Note that these different types of divergences are not mutually exclusive as is evident in the following proposition.

###### Proposition 2.

*The squared norm* 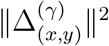 *forms an M-divergence between the diffusion probability distributions* 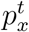 *and* 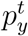 *for all γ* ∈ [−1,1]. *Furthermore*, 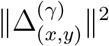 *forms an* f-*divergence for γ* = 0 *and a Bregman divergence for γ* = −1.

*Proof*. For *γ* ∈ (−1,1), it follows that 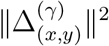 is an M-divergence from Definition 2 with 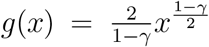. Similarly, *g*(*x*) = *x* and *g*(*x*) = log*x* yield this result for *γ* = −1,1, respectively. For *γ* = 0, we obtain 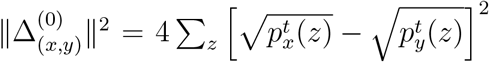, which is indeed an *f*-divergence as it is proportional to the Hellinger divergence. Finally, since *γ* = −1 yields a squared Euclidean distance between the distributions it is indeed a Bregman divergence.

The family of distances (or divergences) formed by 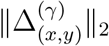 is also directly related to *α*-representations used in information geometries [79] when defining statistical manifolds over probability distributions. Indeed, the *α*-representation of a distribution *p* is defined as

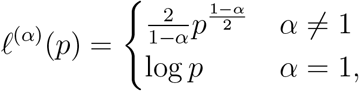

where *ℓ*^(−1)^ and *ℓ*^(+1)^ give rise to the popular mixed family (*m*-family) and exponential family (*e*-family) in information geometry [79,80], correspondingly. We also note at this point that the third popular *α*-family in information geometry is the 0-family, which gives a Fisher geometry as its Riemannian metric as given by Fisher information [80]. Interestingly, this family corresponds to setting *γ* = 0 in our case, which yields a distance 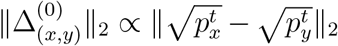 that is proportional to the Hellinger distance between diffusion distributions.

Since, as we discussed here, 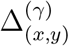 encodes differences between information geometry *ℓ*^(*γ*)^ representations of diffusion distributions, we refer to the distances 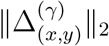 (from Prop. 2) as diffusion-based informational distances. In general, this family of informational distances creates an exciting connection between diffusion geometries and information geometries for exploring emergent structures in data exploration. In particular, in PHATE we focus on the potential distance as an *e*-family distance [79, 80] that combines both the Boltzman distribution law approach and the information geometry approach towards capturing a stable metric structure of the diffusion geometry, and use it for the purpose of visualizing progression by embedding this metric structure in low dimensions.

Figure S2D shows PHATE visualizations of the retinal bipolar data from [23] using different values of *γ*. This figure also indicates the impact of *γ* on the global structure captured by these distances, where indeed the global structure of the potential distance (*γ* = 1) is more similar (as compared to other *γ* values) to the structure captured by PCA, which is known to preserve global structure. Another way to see this is that the structure is unraveled less than when using diffusion distances.

##### Embedding the Potential Distances in Low Dimensions

A popular approach for embedding diffusion geometries is to use the eigendecomposition of the diffusion operator to build a *diffusion map* of the data. However, this approach tends to isolate progression trajectories into numerous diffusion coordinates (i.e., eigenvectors of the diffusion operator; see Figure S1). In fact, this specific property was used in [14] as a heuristic for ordering cells along specific developmental tracks. Therefore, while diffusion maps preserve global structure and denoise the data, their higher intrinsic dimensionality is not amenable for visualization. Instead, we squeeze the variability into low dimensions using metric multidimensional scaling (MDS), a distance embedding method (Figure 2F).

There are multiple approaches to MDS. Classical MDS (CMDS) [7] takes a distance matrix as input and embeds the data into a lower-dimensional space as follows. The squared potential distance matrix is double centered:

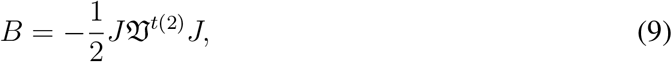

where 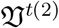 is the squared potential distance matrix (i.e. each entry is squared) and 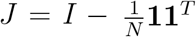 with 1 a vector of ones with length *N*. The CMDS coordinates are then obtained by an eigendecomposition of the matrix *B*. This is equivalent to minimizing the following “strain” function:

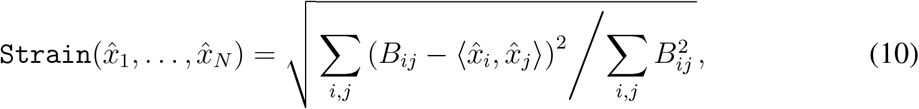

over embedded *m*-dimensional coordinates 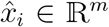 of data points in 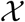. We apply CMDS to the potential distances of the data to obtain an initial configuration of the data in low dimension *m*.

While classical MDS is computationally efficient relative to other MDS approaches, it assumes that the input distances directly correspond to low-dimensional Euclidean distances, which is overly restrictive in our setting. Metric MDS relaxes this assumption by only requiring the input distances to be a distance metric. Metric MDS then embeds the data into lower dimensions by minimizing the following “stress” function:

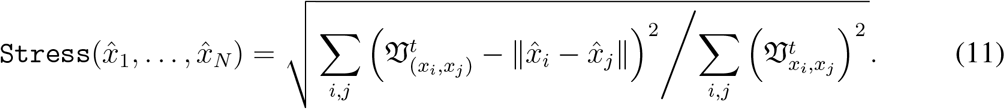

over embedded m-dimensional coordinates 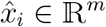 of data points in 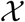.

If the stress of the embedded points is zero, then the input data is faithfully represented in the MDS embedding. The stress may be nonzero due to noise or if the embedded dimension *m* is too small to represent the data without distortion. Thus, by choosing the number of MDS dimensions to be *m* = 2 (or *m* = 3) for visualization purposes, we may trade off distortion in exchange for readily visualizable coordinates. However, some distortion of the distances/dissimilarities is tolerable in many of our applications since precise dissimilarities between points on two different trajectories are not important as long as the trajectories are visually distinguishable. By using metric MDS, we find an embedding of the data with the desired dimension for visualization and the minimum amount of distortion as measured by the stress. When analyzing the PHATE coordinates (e.g. for clustering or branch detection), we use metric MDS with *m* chosen to explain most of the variance in the data as determined by the eigenvalues of the diffusion operator (as is done for von Neumann entropy). In this case, minimal distortion is introduced into the analysis.

A naïve approach towards obtaining a truly low dimensional embedding of diffusion geometries is to directly apply metric MDS, from the diffusion map space to a two dimensional space. However, as seen in Figures S3 (Column 5) and S7, direct embedding of these distances produces distorted visualizations. Embedding the potential distances (defined in Def. 1) is more stable at boundary conditions near end points compared to diffusion maps, even in the case of simple curves that contain no branching points. Figure S2A shows a half circle embedding with diffusion distances versus distances between log-scaled diffusion. We see that points are compressed towards the boundaries of the figure in the former. Additionally, this figure demonstrates that in the case of a full circle (i.e., with no end points or boundary conditions), our potential embedding (PHATE) yields the same representation as diffusion maps.

In some cases, it may be advantageous to relax our assumptions further on the input distances. In this case, non-metric MDS may be used. In contrast with metric MDS, non-metric MDS does not require the input distances to be an actual distance or metric. Non-metric MDS minimizes the differences between a monotonic transformation of the input dissimilarities and the distances in the embedded space. Mathematically, non-metric MDS minimizes the following stress function:

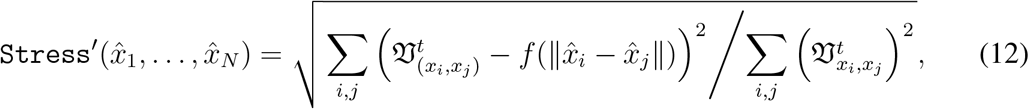

where *f* is a monotonic transformation of the distances between points in the embedded space.

In our experience, the resulting visualizations from metric MDS and non-metric MDS are nearly identical for most datasets. Furthermore, metric MDS is computationally faster than non-metric MDS. Thus, we recommend metric MDS for most problems.

##### Overview Summary

PHATE achieves an embedding that satisfies all four properties delineated above: PHATE preserves and emphasizes the global and local structure of the data via: 1. a localized affinity that is chained via diffusion to form global affinities through the intrinsic geometry of the data, 2. denoises the data by low-pass filtering through diffusion, 3. provides a distance that accounts for local and global relationships in the data and has robust boundary conditions for purposes of visualization, and 4. captures the data in low dimensions, using MDS, for visualization.

We have shown by demonstration in Figures S3 and S7 that all of the steps of PHATE, including the potential transform and MDS, are necessary, as diffusion maps, t-SNE on diffusion maps, and MDS on diffusion maps fail to provide an adequate visualization in several benchmark test cases with known ground truth (even when using the same customized *α*-decaying kernel we developed for PHATE). We have also shown that PHATE is robust to the choice of parameters.

#### A.1.2 Robustness Analysis of PHATE

Here we show that the PHATE embedding is robust to subsampling and the choice of parameters. We demonstrate this both qualitatively and quantitatively. For the quantitative demonstrations, we simulated scRNA-seq data using the Splatter package [22] as in Section 2.2. We first calculated the geodesic pairwise distances for the noiseless data. Then for each setting, we calculated the pairwise Euclidean distances in the 2-dimensional embedding. We then compared the geodesic distances with the embedded distances via the Spearman correlation coefficient as discussed in Section 2.2. We used both the paths and groups options of the Splatter package. Simulation details are found in Appendix A.3.3.

##### Subsampling Robustness

Table 1 shows that PHATE is robust to subsampling on the Splatter datasets. For the paths dataset, the average Spearman correlation is the same when 95% and 50% of the data points are retained. For the groups dataset, the correlation drops slightly when going from 95% retention to 50% retention. Additionally, the correlation coefficient is still quite high (and better than all other methods) when only 5% of the data points are retained. Thus, quantitatively, PHATE is robust to subsampling.

We also demonstrate this visually. We ran PHATE on the iPSC mass cytometry dataset from [16] with varying subsample sizes *N*. Figure S4A shows the PHATE embedding for *N* = 1000, 2500, 5000,10000. Note that the primary branches or trajectories that are visible when *N* = 50000 (Figure S6C) are still visible for all subsamples. Thus, PHATE is robust to the subsampling size. Similar results can be obtained on other datasets.

##### Parameter Robustness

Here, we show that the PHATE embedding is robust to the choice of *t, k*, and *α*. Figure S4B shows the PHATE embedding on the iPSC mass cytometry dataset from [16] with varying scale parameter *t*. This figure shows that the embeddings for 50 ≤ *t* ≤ 200 are nearly identical. Thus, PHATE is very visually robust to the scale parameter *t*. Similar results can be obtained on other datasets and with the *k* and *α* parameters.

The embedding is also quantitatively robust to the parameter choices. Figure S4C-D shows heatmaps of the Spearman correlation coefficient between geodesic distances of the ground truth data and the Euclidean distances of the PHATE visualization applied to the simulated Splatter datasets for different values of *k, t*, and *α*. For *α* ≥ 10, the correlation coefficients are very similar for all values of *k*, *t*, and *α*. This demonstrates that PHATE is robust to the choices of these parameters.

#### A.1.3 Scalability of PHATE

The native form of PHATE is limited in scalability due to the computationally intensive steps of computing potential distances between all pairs of points, computing metric MDS, and storing in memory the diffused operator. Thus, we describe here, and in Algorithm 2, an alternative way to compute a PHATE embedding that is highly scalable and provides a good approximation of the native PHATE described previously. The scalable version of PHATE uses a slight difference in computing t-step diffusion probabilities between points. It requires that every other step that the diffusion takes goes through one of a small number of “landmarks.” Each landmark is selected to be a central point that is representative of a portion of the manifold, selected by spectrally clustering manifold dimensions.

First, we construct the *α*-decaying kernel on the entire dataset. This can be calculated efficiently and stored as a sparse matrix by using radius-based nearest neighbor searches and thresholding (i.e., setting to zero) connections between points below a specified value (e.g., 0.0001), as we regard them numerically insignificant for the constructed diffusion process. The resulting affinity matrix *K_k,α_* will be sparse as long as *α* is sufficiently large (e.g., *α* ≥ 10) to enforce sharp decay of the captured local affinities. The full diffusion operator *P* is constructed from *K_k,α_* by normalizing by row-sums as described previously.

However, powering the sparse diffusion operator would result in a dense matrix, causing memory issues. To avoid this, we instead perform diffusion between points via a series of *M* landmarks where *M* < *N*. We select the landmarks by first applying PCA to the diffusion operator and then using *k*-means clustering on the principal components to partition the data into *M* clusters. This is a variation on spectral clustering. We then calculate the probability of transitioning in a single step from the *i*-th point in 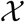 to any point in the *j*-th cluster for all pairs of points and clusters. Mathematically, we can write this as

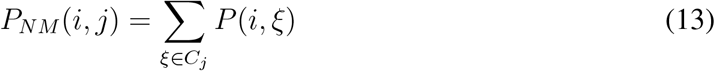

where *C_j_* is the set of points in the *j*th cluster. Thus, we can view each cluster as being represented by a landmark and the (*i, j*)-th entry in *P_NM_* gives the probability of transitioning from the *i*th point in 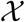 to the *j*-th landmark in a single step. Similarly, we construct the matrix *P_MN_* where the (*j, i*)-th entry contains the probability of transitioning from the *j*-th landmark to the *i*-th point in 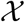. In this case, we cannot simply sum the transition probabilities *P*(*ξ, i*), *ξ* ∈ *C_j_*, since we also have to consider the prior probability *Q*(*j,ξ*) of the *ξ*-th point (with *ξ* ∈ *C_j_*) being the source of a transition from a cluster *C_j_*. For this purpose we use the prior proposed in [81], and write

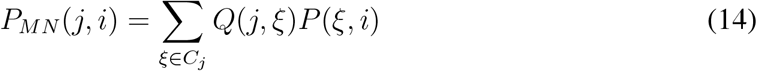

with *Q*(*j,ξ*) = Σ_*i*_ *K_k,α_*(*ξ,i*)/Σ_*ζ∈C_j_*_ Σ_*i*_ *K_k,α_*(*ζ,i*).

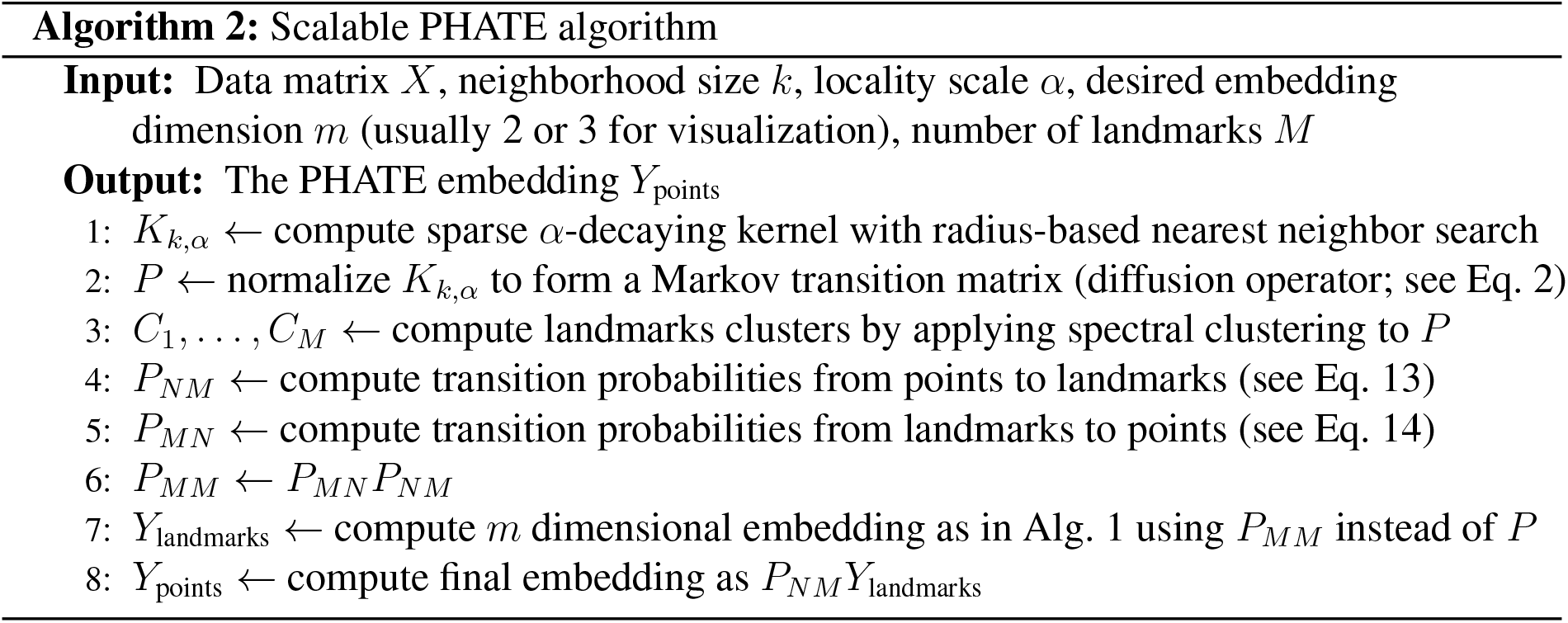

We use the two constructed transition matrices to compute *P_MM_* = *P_MN_P_NM_*, which provides the probability of transitioning from landmark to landmark in a random walk by walking through the full point space. Diffusion is then performed by powering the matrix *P_MM_*. This can be written as

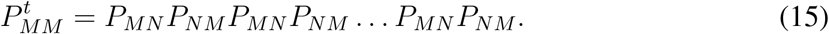

From this expression, we see that powering the matrix *P_MM_* is equivalent to taking a random walk between landmarks by walking from landmarks to points and then back to landmarks *t* times.

We then embed the landmarks into the PHATE space by calculating the potential distances between landmarks and applying metric MDS to the potential distances. Denote the resulting embedding as *Y*_landmarks_. We then perform an out of sample extension to all points from the landmarks by multiplying the point to landmark transition matrix *P_NM_* by *Y*_landmarks_ to get

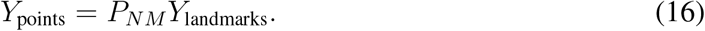

Since *M* is chosen to be vastly less than *N*, the memory requirements and computational demands of the powering the diffusion operator and embedding the potential distances are much lower.

The described steps are summarized in Algorithm 2. In Figure S5A-E we show that this constrained diffusion preserves distances between datapoints in the final PHATE embedding, with the scalable version giving near-identical results to the exact computation of PHATE. Further, in Figure S5B we show that the embedding achieved by this approach is robust to the number of landmarks chosen.

We note that if the only computational bottleneck were in computing MDS, scalable versions of MDS could be used [8,82,83]. However, since storing the entries of the powered diffusion operator in memory is also an issue, we employ the use of landmarks earlier in the process. It has also been shown that “compressing” the process of diffusion through landmarks in the fashion described here performs better than simply applying Nystrom extension (which includes landmark MDS [82]) to diffusion maps [84].

The fast version of PHATE was used in Figures 5, S7, S3, S2D, S5A-E, S10, and S11. All other plots were generated using the exact version of PHATE.

##### PHATE on Large Scale Datasets

To demonstrate the scalability of PHATE for data exploration on large datasets, we applied PHATE to the 1.3 million mouse brain cell dataset from 10x [85]. Figure S5C shows a comparison of PHATE to t-SNE, colored by 10 of the 60 clusters provided by 10x. We see that PHATE retains cluster coherence while t-SNE shatters some of the cluster structure.

#### A.1.4 Branch Point Detection

Here we describe the methods we developed for identifying branch points and selecting representative branch- and endpoints.

##### Branch Point Identification

We used the local intrinsic dimension estimation method derived in [17, 86] to provide suggested branch points. The procedure is as follows. Let **Z**_*n*_ = {**z**_1_,…, **z**_*n*_} be a set of independent and identically distributed random vectors with values in a compact subset of ℝ^*d*^. Let 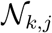 be the *k* nearest neighbors of **z**_*j*_; i.e. 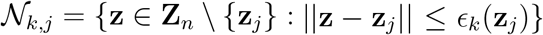. The *k*-nn graph is formed by assigning edges between a point in **Z**_*n*_ and its *k*-nearest neighbors. The power-weighted total edge length of the *k*-nn graph is related to the intrinsic dimension of the data and is defined as

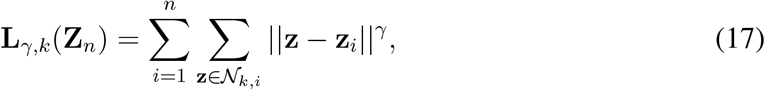

where *γ* > 0 is a power weighting constant. Let *m* be the global intrinsic dimension of all the data points in **Z**_*n*_. It can be shown that for large *n*,

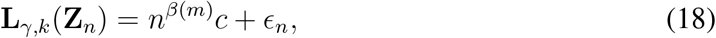

where *β*(*m*) = (*m* − *γ*)/*m, ϵ_n_* is an error term that decreases to 0 as *n* → ∞, and *c* is a constant with respect to *β*(*m*) [86]. A global intrinsic dimension estimator *m* can be defined based on this relationship using non-linear least squares regression over different values of *n* [17,86].

A local estimator of intrinsic dimension 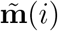 at a point **z**_*i*_ can be defined by running the above procedure in a smaller neighborhood about **z**_*i*_. This approach is demonstrated in Figure 3A, where a *k*-nn graph is grown locally at each point in the data. However, this estimator can have high variance within a neighborhood. To reduce this variance, majority voting within a neighborhood of **z**_*i*_ can be performed:

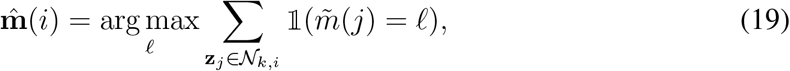

where 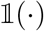 is the indicator function [17].

##### Representative Point Selection with Shake and Bake

To reduce branch points and endpoints to a set of representative points for branch analysis, we use a shake and bake procedure similar to that in [20]. Let 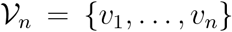 be the set of branch points and endpoints in the high-dimensional PHATE coordinates that we wish to reduce. We create a Voronoi partitioning of these points as follows. We first permute the order of 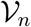, which we denote as 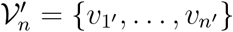. We then take the first point *υ*_1′_ and find all the points in 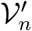 that are within a distance of *h*, where *h* is a scale parameter provided by the user. These points (including *υ*_1′_) are assigned to the first component of the partition and removed from the set 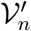. This process is then repeated until all points in *V_n_* are assigned to the partition. To ensure that each point is assigned to the nearest component of the partition (as measured by proximity to the centroid), we next calculate the distance of each point to all centroids of the partition, and reassign the point to the component with the nearest centroid. This reassignment process is repeated until a stable partition is achieved. This completes the process of constructing the Voronoi partition.

The Voronoi partition constructed from this process may be sensitive to the ordering of the points in 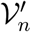. To reduce this sensitivity, we repeat this process multiple times (e.g., 40-100) to create multiple Voronoi partitions. We then construct a distance between points by estimating the probability that two points are not in the same component from this partitioning process. This provides a notion of distance that is robust to noise, random permutations, and the scale parameter *h*. We then partition the data again using the above procedure except we use these probability-based distances. The representative points are then selected from the resulting centroids of this final partition.

##### Branch Detection

We now describe how we assign data points to branches using the correlation and anticorrelation of neighborhood distances (in higher dimensional PHATE coordinates). The approach is demonstrated visually in Figure 3Aiii. Here we consider two reference cells *X* and *Y*. We wish to determine if cells *Q*_1_ and *Q*_2_ belong to the branch between *X* and *Y* or not. Consider *Q*_1_ first which does belong to this branch. If we move from Q1 towards *X*, we also move farther away from *Y*. Thus the distances to *X* and *Y* of a neighborhood of points around *Q*_1_ (which will be located on the branch) are negatively correlated with each other. Now consider *Q*_2_ which does not belong to the branch between *X* and *Y*. In this case, if we move from *Q*_2_ towards *Y*, we also move closer to *X*. Thus the distances to *X* and *Y* of a neighborhood of points around *Q*_2_ are positively correlated with each other. In practice, these distance-based correlations are computed for each possible branch and the point is assigned to the branch with the largest anticorrelation (i.e. the most negative correlation coefficient).

#### A.1.5 EMD Score Analysis

The EMD is measure of dissimilarity between two probability distributions that is particularly popular in computer vision [87]. The EMD was chosen to perform differential expression analysis in the EB scRNA-seq data due to its stability in estimation compared to other divergence measures. Intuitively, if each distribution is viewed as a pile of dirt, the EMD can be thought of as the minimum cost of converting one pile of dirt into the other. If the distributions are identical, then the cost is zero. When comparing univariate distributions (as we do as we only consider a single gene at a time), the EMD simplifies to the *L*^1^ distance between the cumulative distribution functions [58]. That is, if *P* and *Q* are the cumulative distributions of densities *p* and *q*, respectively, then the EMD between *p* and *q* is ∫ |*P*(*x*) − *Q*(*x*)|*dx*. While the EMD is nonnegative, we assign a sign to the EMD score based on the difference between the medians of the distributions.

### A.2 Biological Methods

The processes for generating the EB data and for preprocessing the biological data are described here.

#### A.2.1 Generation of Human Embryoid Body Data

Low passage H1 hESCs were maintained on Matrigel-coated dishes in DMEM/F12-N2B27 media supplemented with FGF2. For EB formation, cells were treated with Dispase, dissociated into small clumps and plated in non-adherent plates in media supplemented with 20% FBS, which was prescreened for EB differentiation. Samples were collected during 3-day intervals during a 27 day-long differentiation timecourse. An undifferentiated hESC sample was also included (Figure S12A). Induction of key germ layer markers in these EB cultures was validated by qPCR (data not shown). For single cell analyses, EB cultures were dissociated, FACS sorted to remove doublets and dead cells and processed on a 10x genomics instrument to generate cDNA libraries, which were then sequenced. Small scale sequencing determined that we have successfully collected data on approximately 31,000 cells equally distributed throughout the timecourse.

#### A.2.2 Data Preprocessing

In this section, we discuss methods we used to preprocess the various datasets.

##### Data Subsampling

The full PHATE implementation scales well for sample sizes up to approximately *N* = 50000. For *N* much larger than 50000, computational complexity can become an issue due to the multiple matrix operations required. All of the scRNAseq datasets considered in this paper have *N* < 50000. Thus, we used the full data and did not subsample these datasets. However, the mass cytometry datasets have much larger sample sizes. To aid in branch analysis, we randomly subsampled these datasets for analysis in Section 3 using uniform subsampling. For the comparison figures (Figures 5, S3, and S7), scalable PHATE was used and subsampling was not performed except as indicated in the figures. The PHATE embedding is robust to the number of samples chosen, which we demonstrated in Appendix A.1.2.

##### Mass Cytometry Data Preprocessing

We process the mass cytometry datasets according to [88].

##### Single-Cell RNA-Sequencing Data Preprocessing

This data was processed from raw reads to molecule counts using the Cell Ranger pipeline [89]. Additionally, to minimize the effects of experimental artifacts on our analysis, we preprocess the scRNAseq data. We first filter out dead cells by removing cells that have high expression levels in mitochondrial DNA. In the case of the EB data which had a wide variation in library size, we then remove cells that are either below the 20th percentile or above the 80th percentile in library size. scRNA-seq data have large cell-to-cell variations in the number of observed molecules in each cell or *library size*. Some cells are highly sampled with many transcripts, while other cells are sampled with fewer. This variation is often caused by technical variations due to enzymatic steps including lysis efficiency, mRNA capture efficiency, and the efficiency of multiple amplification rounds [90]. Removing cells with extreme library size values helps to correct for these technical variations. We then drop genes that are only expressed in a few cells and then perform library size normalization. Normalization is accomplished by dividing the expression level of each gene in a cell by the library size of the corresponding cell.

After normalizing by the library size, we take the square root transform of the data and then perform PCA to improve the robustness and reliability of the constructed affinity matrix *K_k,α_*.

We choose the number of principal components to retain approximately 70% of the variance in the data which results in 20-50 principal components.

##### Gut Microbiome Data Preprocessing

We use the cleaned L6 American Gut data and remove samples that are near duplicates of other samples. We then preprocess the data using a similar approach for scRNA-seq data. We first perform “library size” normalization to account for technical variations in different samples. We then log transform the data and then use PCA to reduce the data to 30 dimensions.

Applying PHATE to this data reveals several outlier samples that are very far from the rest of the data. We remove these samples and then reapply PHATE to the log-transformed data to obtain the results that are shown in Figure 1D.

##### ChIP-seq Processing for Hi-C Visualization

We used narrow peak bed files and took the average peak intensity for each bin at a 10 kb resolution. For visualization, we smoothed the average peak intensity values based on location using a 25 bin moving average.

### A.3 Comparison Details

Here we provide details for the construction of the DEMaP score that we use to compare between dimensionality reduction methods. We also provide details on generating the artificial tree data and Splatter datasets, and for the parameter settings for all PHATE plots in the paper.

#### A.3.1 DEMaP

To quantitatively compare each dimensionality reduction tool, we wish to calculate the degree to which each method preserves the underlying structure of the reference dataset and removes noise. Since single-cell RNA-sequencing and other biological types of data are highly noisy, visual renderings of the data that can offer denoised embeddings that reveal the underlying structure of the data are desirable. Therefore, the goal of our accuracy metric is to quantify the correspondence between distances in the low-dimensional embedding and manifold distances in the ground truth reference.

To define a quantitative notion of manifold distance we use geodesic distances. Geodesic distances are shortest path distances on a nearest-neighbor graph of the data weighted by the Euclidean distances between connected points [4]. In cases where points are sampled noiselessly from a manifold, such as in our ground truth reference, geodesic distances converge exactly to distances along the manifold of the data [4,91]. Thus we reason that if geodesic distances between points on the noiseless manifold are preserved by an embedding computed on the noisy data then the data is sufficiently denoised and the manifold structure is also preserved.

We take this approach to formulate our ground-truth manifold distance as a quantification of the degree to which each dimensionality reduction method preserves the pairwise geodesic distances of the noiseless data after low-dimensional embedding of the corresponding noisy data. Since the low dimensional embedding is often a result of a non-linear dimensionality reduction, curves and major paths in the data are “straightened” such that Euclidean distances in the embedding space correspond to manifold distance in the high dimensional space [7]. Thus we quantify the preservation of manifold distances as the correlation between geodesic distance in the noiseless reference dataset and Euclidean distances in the embedding space as a measure of structure preservation which we call *Denoised Embedding Manifold Preservation (DEMaP)*. An overview of DEMaP is presented in Figure 4A.

#### A.3.2 Construction of the Artificial Tree Test Case

The artificial tree data shown in Figure 1B is constructed as follows. The first branch consists of 100 linearly spaced points that progress in the first four dimensions. All other dimensions are set to zero. The 100 points in the second branch are constant in the first four dimensions with a constant value equal to the endpoint of the first branch. The next four dimensions then progress linearly in this branch while all other dimensions are set to zero. The third branch is constructed similarly except the progression occurs in dimensions 9-12 instead of dimensions 5-8. All remaining branches are constructed similarly with some variation in the length of the branches. We then add 40 points at each endpoint and branch point and add zero mean Gaussian noise with a standard deviation of 7. This construction models a system where progression along a branch corresponds to an increase in gene expression in several genes. Prior to adding noise, we also constructed a small gap between the first branch point and the orange branch that splits into a blue and purple branch (see the top set of branches in the left part of Figure 1B). This simulates gaps that are often present in measured biological data. We also added additional noise dimensions, bringing the total dimensionality of the data to 60.

#### A.3.3 Splatter Simulation Details

Splatter is a scRNA-seq simulation package that uses a parametric model to generate data with various structures, such as branches or clusters [22]. We use Splatter to simulate multiple ground truth datasets for multiple experiments. To select parameters for the simulation, we fit the Splatter simulation to the EB data, and then modified the resulting dataset from both the Splatter “paths” and the Splatter “groups” simulations as described in Section 2.2. Note that we do not make use of Splatter’s built-in dropout function, since it uses a zero-inflated model; multiple studies have shown that an undersampling (binomial) model is more appropriate [92–96]. Each simulation is performed with 3000 simulated cells. The mean correlation coefficient and standard deviations are calculated from 20 trials.

The default parameters used in the simulation are the following: batchCells = 3000, nGenes=17580, mean.shape=6.6, mean.rate=0.45, lib.loc=9.1, lib.scale=0.33, out.prob=0.016, out.facLoc=5.4, out.facScale=0.90, bcv.common=0.18, bcv.df=21.6, de.prob=0.2, dropout.type=“none”, with a post-hoc binomial dropout of 50%. For the groups simulation we drew the number of groups *n* from a Poisson distribution with rate λ = 10, and then drew the `group.prob` parameter from a Dirich-let distribution with *n* categories and a uniform concentration *α*_1_ = ⋯ = *α_n_* =1. For the paths simulation, we set `group.prob` as above, and additionally set the *i*th entry in the parameter `path.from` as a random integer between 0 and *i* − 1, drew the parameter `path.nonlinearProb` from a uniform distribution on the interval (0,1), and drew the parameter `path.skew` from a beta distribution with shape *α* = 10, *β* = 10. Note that here the library size was doubled from the fit value, since the EB data itself suffers from dropout. To reduce the number of genes for the *n_genes* simulation, we randomly removed genes post-hoc in order to avoid changing the state of the random number generator in building the simulation.

For the ground truth simulations, we set `bcv.common` to 0, did not perform binomial dropout, and did not remove genes or cells. For the *BCV* simulation, we performed 50% post-hoc binomial dropout, did not remove genes or cells, and set `bcv.common` to 0, 0.25, and 0.5. For the *dropout* simulation, we set `bcv.common` to 0.18, did not remove genes or cells, and performed 0%, 50%, and 95% post-hoc binomial dropout. For the *subsample* simulation, we set `bcv.common` to 0.18, performed 50% post-hoc binomial dropout, did not remove genes, and subsampled rows of the matrix to retain 95%, 50%, and 5% of the total cells. For the *n-genes* simulation, we set `bcv.common` to 0.18, performed 50% post-hoc binomial dropout, did not remove cells, and subsampled columns of the matrix to retain 17000, 10000, and 2000 genes.

#### A.3.4 PHATE Experimental Details

For all of the quantitative comparisons, we have used the default parameter settings for the PHATE plots. For the majority of the qualitative comparisons in Figures 5, S7, and S3, we also used the default parameter settings for all methods. Exceptions to this are the artificial tree (Figure S3A), the intersecting circles (Figure S3D), and the MNIST dataset (Figure S3I). In these cases, the PHATE parameters have been tuned to give a clearer separation of the branches. However, in general, the default PHATE settings give good results on most datasets, especially those that are complex, high-dimensional, and noisy as demonstrated empirically in Appendix A.1.2. The default settings are also used in Figures S2D, S5A-E, S10, and S11. For all other PHATE plots, the parameters were tuned slightly to better highlight the structure of the data.

### A.4 Data Availability

The embryoid body scRNA-seq and bulk RNA-seq datasets generated and analyzed during the current study are available in the Mendeley Data repository at: http://dx.doi.org/10.17632/v6n743h5ng.1 Figure S12A contains images of the raw single cells while Figure S12F contains scatter plots showing the gating procedure for FACS sorting cell populations for the bulk RNA-seq data.

### A.5 Code Availability

Python, R, and Matlab implementations of PHATE are available on GitHub, for academic use: https://github.com/KrishnaswamyLab/PHATE An interactive tool for exploring PHATE on the EB data is available at: https://www.krishnaswamylab.org/phatewebtool

## B Extended Comparison of PHATE to Other Methods

Here we provide an extended qualitative comparison of PHATE to methods of dimensionality reduction, graph rendering, and visualization on multiple datasets. These comparisons are found in Figures S3 and S7. Details on parameter selection are included in Appendix A.3.4.

Figure S3 shows comparisons on non-biological datasets. These include (in the order shown) A. Artificial tree data with 7 branches in 60 dimensions; B. Simulated Splatter path data [22]; C. Simulated Splatter groups data [22]; D. Three intersecting curves in 3 dimensions; E. Video of a rotating teapot [97]; F. Swiss roll data used in [4]; G. Frey faces video [6]; H. Columbia Object Image Library (COIL-20), 20 videos of rotating objects [98]; I. MNIST, 70,000 images of handwritten digits from 0 to 9 [99].

Figure S7 shows comparisons on biological datasets including (in the order shown) A. Developing mouse bone marrow cells, enriched for the myeloid and erythroid lineages, which were measured with the MARS-seq single cell RNA-sequencing technology [21]; B. Single cell RNA-sequencing of epithelial cells from mouse small intestine and organoids [100]; C. Mass cytometry data measuring T cell development into CD8+ and CD4+ T cells in mouse thymus [27]; D. New embryoid body data from a 27-day timecourse; E. Mass cytometry data showing iPSC reprogramming of mouse embryonic fibroblasts [16]; F. Single cell RNA-sequencing of mouse retinal bipolar cells [23]; G. Single cell RNA-sequencing of mouse cortical cells from the somatosensory cortex and hippocampal CA1 region [24].

Some of these biological datasets represent differentiating processes within the body, and hence visualizing progression is key to understanding the structure of these datasets. Other artificial datasets show a combination of clusters and trajectories, while the artificial datasets give a plausible range of manifold structures that could be found in biological data.

PHATE is primarily a dimensionality reduction method that takes high dimensional raw data and embeds it, via a metric preserving embedding, into low dimensions that naturally show trajectory structure. Thus, we focus our comparisons of PHATE to existing dimensionality reduction methods such as PCA, t-SNE, and diffusion maps.

We note that several methods exist that focus on finding *pseudotime* orderings of cells, such as Wanderlust [26], Wishbone [27], and diffusion pseudotime [14]. Wanderlust can find single non-branching progressions. Wishbone recognizes a single branch, while diffusion pseudotimes provides potentially multiple branches. However, pseudotime approaches do not naturally provide a dimensionality reduction method to visualize such structure. Therefore, the resulting cell orderings can be difficult to interpret and verify, especially in the context of the entire data set. Since PHATE reveals the entire branching structure in low dimensions, it can be beneficial to visualize, verify and interpret the results of any pseudotime algorithm displayed on PHATE.

These pseudotime methods can also be used alongside PHATE to order parts of the branching progressions. Indeed, we use Wanderlust on the EB data in Section 4 to extract ordering from the branches identified from PHATE. A comprehensive overview and comparison of state of the art pseudotime and trajectory inference methods for single-cell RNA sequencing is given in [101].

A few pseudotime methods can create a visualization based on finding and rendering a differentiation tree structure, including SPADE [102,103] and Monocle2 [12]. However, SPADE is not well-suited for analyzing scRNA-seq data without prior knowledge (see [103] and Section B.2). Thus we only compare to Monocle2.

Note that we do not include any metadata, such as sample time or clusters, in any of the analyses. Therefore, we are focusing on the performance of these methods in an unsupervised setting. Instead, we use this metadata as a tool for comparing the results of the various methods.

### B.1 Comparison of PHATE to Dimensionality Reduction Methods

Figures S3 and S7 compare the PHATE visualization to the dimensionality reduction methods of principal components analysis (PCA), Diffusion Maps (DM), Metric Multidimensional Scaling (MDS), MDS performed on Diffusion Maps, t=SNE, t-SNE performed on Diffusion Maps, Local Linear Embedding (LLE), Isomap, Force Directed Layout, UMAP and Monocle2 on nine non-biological datasets and seven biological datasets, including the five biological datasets shown on a subset of these methods in Figure 5. The datasets show a combination of intersecting and distinct manifolds, clusters, and branching trajectories, examining a range of both real biological manifolds and plausible challenging structures for visualization. The PHATE visualization most consistently distinguishes branches, trajectories and clusters in order to give a denoised representation of the underlying structure of the data. We focus on each method individually.

#### Comparison of PHATE to PCA

PCA is a popular method of data analysis that uses eigen-decomposition of the covariance matrix to learn axes within the high-dimensional data that account for the largest amount of variance within the data [11]. However, PCA assumes a linear structure on the data, the visualization amounts to projecting the data onto a slicing plane, which creates a noisy visualization. Also, since biological data are rarely linear, PCA is unable to optimally reduce non-linear noise along the manifold and reveal progression structure in low dimensions. This is evident in Figure S3A where we compare PCA to PHATE on artificial tree data. This data contains seven distinct branches uniformly sampled in 60 dimensions. See Appendix A.3.2 for details. PCA does capture some of the global structure in this relatively low-noise data. However, many branches are not visible in the first two PCA dimensions and the trajectories in the PCA visualization are noisy compared to the PHATE visualization, in which all seven branches are easily identifiable.

For the other datasets in Figures S3 and S7, PCA captures some of the overall global structure of the datasets. For example, the PCA dimensions in Figure S7D encode the overall time progression of the noisy EB scRNA-seq data. Thus, PCA captures some of the global structure. However, PCA presents mostly a cloud of cells in this case and any finer branching structure is not visible. This contrasts with PHATE which shows multiple branches and trajectories. Similar results are obtained from the mouse bone marrow scRNA-seq and iPSC CyTOF datasets in Figures S7A and E respectively, demonstrating that PCA is unable to accurately visualize the global and local structure of the data simultaneously.

#### Comparison of PHATE to t-SNE

t-SNE (t-distributed stochastic neighbor embedding) [1] is a visualization method that emphasizes local neighborhood structure within data. Recently, t-SNE has become popular for revealing cluster structure or separations in single cell data [2]. However, due to its emphasis on preserving local neighborhoods, t-SNE tends to shatter trajectories into clusters as seen in the artificial tree and intersecting curves in Figures S3A and D as well as the EB data and the iPSC data in Figures S7D and E. In all of these cases, the data naturally have a strong trajectory structure either by design (the artificial trees and intersecting curves) or due to the developmental nature of the data (the EB and iPSC datasets). Thus t-SNE creates the false impression that the data contain natural clusters, which could lead to incorrect analysis.

Furthermore, the adaptive kernel used in t-SNE for calculating neighborhood probabilities tends to spread out neighbors such that dense clusters occupy proportionally more space in the visualization compared to sparse clusters [104]. Thus, the relative location of data points within the t-SNE embedding often does not accurately reflect the relationships between them. This is clearly visible in the t-SNE plot in Figure S3A where the shattered branches are located far away from where they originated in the main structure in the artificial tree data. Similarly, t-SNE creates clusters in the EB and iPSC data in Figures S7D and E which split the time samples into different components. Since the relative position of clusters in t-SNE is generally meaningless the overall progression of the data is destroyed.

Even in the case where the data are more naturally separated into clusters, t-SNE can destroy the global information about the relative relationships between clusters due to this weakness. In contrast, PHATE separates clusters that are sufficiently separated from each other (see Figure S3C) while maintaining the relative relationships of clusters based on the relative positions of the clusters in the PHATE embedding. In other words, PHATE preserves both the global and local structure while t-SNE only preserves local structure.

One proposed solution to the failure of t-SNE to retain global structure is to use a random walk to learn the global structure, and then apply t-SNE to the resulting kernel [1]. One approach to do this is to apply t-SNE to DM. However, our experiments show that this fails to capture the global structure of the data. In the artificial tree data (Figure S3A) the intersecting curves (Figure S3D), as well as the new EB data (Figure S7D) t-SNE on DM shatters trajectories. Furthermore, in the retinal bipolar dataset (Figure S7F) t-SNE on DM shatters clusters and creates misleading trajectory-like structures. Hence, performing t-SNE on diffusion maps suffers from the same shortcomings as t-SNE, with additional distortion from the denoising aspect of diffusion maps, as t-SNE tends to shatter trajectories less frequently on noisier data (see Figures S3A and B). Due to the nature of the t-SNE penalty function, global distances encoded in the diffusion distances are ignored, and the resulting embedding is a denoised equivalent of t-SNE, which is more prone to shattering trajectories than t-SNE and lacks the global structure benefits from Diffusion Maps.

#### Comparison of PHATE to Diffusion Maps

Diffusion maps effectively encode continuous relationships between cells. However, different trajectories are often encoded in different dimensions (i.e., since they represent different meta-stable states of the diffusion process) as seen in Figure S1, which is unsuitable for visualization. In contrast, PHATE effectively encodes trajectories in lower dimensions for visualization. This is also seen clearly in Figures S3A and D in the comparison of PHATE to diffusion maps on the artificial tree and intersecting curves data (for each data set, the same kernel and diffusion scale *t* is used for both diffusion maps and PHATE). In this case, the diffusion maps visualization is denoised and the global structure is visible. However, multiple branches are not visible in the low-dimensional visualization of diffusion maps. In fact, approximately six diffusion maps coordinates are needed to separate all ten branches (see Figure S1). In contrast, all of the ten branches of the artificial tree data are clearly visible in the PHATE visualization. Similarly, multiple branches that are visible in the PHATE visualization are not visible in the diffusion maps visualization for the bone marrow and EB scRNA-seq datasets (Figures S7A and D, respectively). Additionally, the diffusion maps instabilities mentioned previously appear to cause very noisy data (e.g. the scRNA-seq data in Figures S7A, B, and D) to contract too much into thin trajectories, which can distort some of the underlying progression structure. In summary, while diffusion maps works well for nonlinear dimensionality reduction, it is not well-suited for visualizing data with multiple trajectories due to its instabilities and its propensity to encode different trajectories in different dimensions.

A logical question is whether applying MDS on diffusion distances would be sufficient for encoding the high dimensional spatial information from diffusion maps in low dimensions for visualization. However, in Figures S3D and H on the intersecting curves and COIL20 we show that MDS on DM suffers equivalently from instability at boundary conditions and intersections of manifolds, producing a totally structureless embedding. Additionally, MDS on DM collapses trajectories into thin trajectories as shown on the mouse bone marrow scRNAseq, EB data, and iPSC CyTOF data (Figures S7A, D, and E respectively), masking the intricate structure visible in PHATE. Thus, performing MDS on diffusion maps without applying the informational potential transformation of PHATE is insufficient for high quality visualization.

#### Comparison of PHATE to MDS

Multidimensional scaling (MDS) [105] aims to preserve or approximate the metric structure of the data by optimizing a stress loss between original (Euclidean) distances and embedded ones. While MDS ostensibly preserves both local and global distances, it does not rely on or infer any particular intrinsic structure from the data. Therefore, it cannot separate meaningful relations in complex high dimensional data from superfluous ones. In particular, this causes MDS to be strongly affected by noise, as shown in Splatter simulations in Figures S3B and C where the embeddings show no structure at all. Further, this causes problems in visualizing noisy biological datasets, such as the new EB scRNAseq data (Figure S7D) where local trajectories are lost, and the iPSC CyTOF data (Figure S7E) where the branching structure is entirely masked by noise. MDS also fails to separate clusters in noisy data, as shown in biological datasets from [24] and [23] (Figures S7F and G) as well as in MNIST (Figure S3I).

#### Comparison of PHATE to Isomap

Isomap [4] embeds the intrinsic metric structure of the data by applying MDS to geodesic distances, which are obtained by constructing a *k*-nearest neighbor graph over the data, and then applying all-pairs shortest path search (e.g., Dijkstra [106] or Floyd [107]) to compute distances. Like other manifold learning methods, Isomap works under the assumption that the data is sampled from an underlying manifold, and thus the geodesic distances approximate intrinsic manifold distances and the coordinate assigned by MDS should provide a global intrinsic coordinate system. However, this assumption is mainly valid when the manifold itself is convex, with no holes, and the data is sampled uniformly from it with only small amount of noise away from the manifold. Indeed, the main weaknesses of Isomap is its topological instability when such assumptions are not satisfied in practice [25,91,108], as we also show here. These instabilities render Isomap susceptible to spurious connections created in noisy datasets, as shown by the failure to separate branches on the artificial tree and Splatter paths (Figure S3A and B). Further, Isomap is incapable of embedding clusters, such as the Splatter groups (Figure S3D), the mouse retinal bipolar scRNAseq (Figure S7F), and MNIST (Figure S3I), in which many clusters are merged together since geodesic distances do not consider the data distribution and do not quantify relations between disconnected clusters. Additionally, Isomap is also unstable to intersecting manifolds, as shown by the Intersecting Curves dataset, where Isomap fails to distinguish between two of the curves and shows no intersections (Figure S3D), and does not clearly display branching points, as shown on the iPSC data where Isomap only separates points by timepoint (Figure S7E).

#### Comparison of PHATE to Locally Linear Embedding (LLE)

LLE [6] is a manifold learning algorithm that is similar to Isomap in that it assumes the data is sampled from a single smooth manifold, approximated by a *k*-NN graph, and tries to use its geometric properties to embed the data. However, unlike Isomap, it only considers local information - namely, it uses the low dimensional coordinate neighborhoods, which are (independently) linearly related to local manifold patches, and attempts to tile these local coordinates into a consistent global embedding in low dimensions. This is done by first optimizing weights that allow the approximation of each data point as a linear combination of its neighbors, and then optimizing low dimensional coordinates that preserve the same linear relations encoded by these weights.

While LLE is less susceptible to shortcut connections, it heavily relies on a smooth well-connected manifold structure to provide global connections through the data. Therefore, LLE is ill-suited for embedding separate clusters, as shown by its failure to separate the distinct digits in the MNIST dataset (Figure S3I). Further, due to its reliance on local interpolation for encoding relations between each point and its neighbors, it strongly affected by boundary conditions that make such interpolation unstable, as shown by its complete failure to embed the Gaussian Mixture Model (Figure S3C). Additionally, LLE does not handle intersecting manifolds and high curvatures, due its implicit assumption that local data patches correspond to manifold coordinate neighborhoods, which can be linearly approximated by a tangent space of the same intrinsic dimension of the manifold. For example, on the Intersecting Curves dataset, while PHATE shows a clean intersection between the curves, LLE treats one of the intersections as a noisy subcluster, and the other intersection is not shown, with the orange curve shown simply as a continuation of the blue curve (Figure S3D). Finally, we note that some additional weaknesses of LLE for visualization of even simple synthetic biomedical data were reported previously, e.g., in [25,109].

#### Comparison of PHATE to UMAP

Like t-SNE, upon which the UMAP algorithm is modeled, UMAP encourages formation of clusters even when cluster structures do not exist. As such, UMAP shatters trajectories, as shown in the artificial trees and intersecting circles (Figures S3A and D respectively) as well as in the new EB data (Figure S7D). Additionally, although UMAP’s algorithm is designed to respect global relationships between the clusters that it forms, this does not always hold between distant clusters; take for example the artificial tree, in which UMAP places the shattered cyan branch at the opposite end of the plot to the green and yellow branches to which it is connected, and does the same for the purple branch, which should be connected to be the brown, orange and red branches, but is placed far from all of these (Figure S3A).

### B.2 Comparison of PHATE to Graph-Rendering Methods

Graph-rendering methods differ fundamentally from dimensionality reduction methods in that they do not produce a reduced dimension representation of the data and instead focus only on providing a specific rendering of the data. However, these renderings are often limited by structural assumptions on the data (e.g. a tree) that may be inaccurate. Figures S3 and S7 compare the PHATE visualization to graph-rending methods including Force Directed Layout (FDL) and Monocle2.

#### Comparison of PHATE to Force-Directed Methods

FDL algorithms attempt to draw a weighted graph in a two-dimensional space so that all edge weights are approximately preserved as distances and relatively few edges cross each other. To this end, such methods solve a *n*-body problem modeled by attractive and repulsive forces that resemble physical systems, such as elastic (e.g., Hooke’s law), electric (e.g., Coulomb’s law), or nuclear forces. Typically, repulsive forces are used to separate and spread out all pairs of nodes while attractive ones are used to keep neighboring nodes on the graph close to each other in the embedding. These attractive forces are scaled based on the strength of the connections within the graph. We specifically apply the Spring Layout method within the NetworkX Python package [110] to the graph defined by the *α*-decaying kernel produced for PHATE. This method uses the Fruchterman-Reingold algorithm [111], which is motivated by the aesthetics of rendering a planar graph, and models the attractive and repulsive forces based on a combination of notions elastic attraction, nuclear repulsion, and global stability criteria (e.g., ideal uniform distance and decaying temperature of the system).

FDL algorithms are generally computationally expensive, making it difficult to scale them to larger datasets (Figure S5E). FDL algorithms also do not denoise the connections between data points, but rather assume that attractive and repulsive forces will eventually balance each other. Thus, they suffer from the same sensitivity to graph construction as Isomap, where spurious connections in noisy data can dominate the resulting force-dynamics and strongly affect the embedding (see the artificial tree and intersecting curves in Figures S3A and D respectively). Additionally, exceedingly weak forces between distant clusters, which are clearly not well-connected in the graph, lead to a failure to retain long-range global distances, as shown in Figure S7F, in which rod bipolar cells and cone bipolar cells do not appear substantially distinct. These shortcomings can lead to the loss of information, such as a lack of branching structure in the iPSC CyTOF data, or a merging of cell types in the neuronal scRNAseq from [24] (Figures S7E and G respectively.)

#### Comparison of PHATE to Tree-Rendering Methods (SPADE and Monocle2)

SPADE [102,103] and Monocle2 [12] are popular methods that fit the data to a predetermined structure such as a tree. These methods first attempt to do data reduction by clustering the data. Clustering methods tend to make less restrictive assumptions on the structure of the data compared to PCA. However, clustering methods assume that the underlying data can be partitioned into discrete separate regions. In reality, biological data are often continuous, and the apparent cluster structure given by clustering methods is only a result of non-uniform density and finite sampling of the continuous underlying state space. Additionally, the results from these methods will be incorrect if the underlying data does not lie on a tree. In contrast, PHATE does not make any assumptions on the data and instead learns the underlying structure.

SPADE fits a minimal spanning tree to the clusters and was originally designed for mass cytometry data [102]. In Anchang et al. [103], the authors applied SPADE to scRNA-seq data by selecting relevant genes to perform dimensionality reduction. This makes it difficult to do data exploration as gene selection must be performed first. In contrast, PHATE does not require any gene selection procedure although PHATE can be used to analyze specific genes of interest by including only the relevant genes. SPADE has several other limitations according to Anchang et al. [103]. First, the SPADE results can be sensitive to the number of clusters which must be specified by the user. Second, down-sampling is required to visualize large datasets. SPADE is very sensitive to random down-sampling and will produce very different trees even when only down-sampling to 99% [103]. Thus, the random nature of the SPADE results make it difficult to discover the right structure of the data with SPADE. Given these limitations, we do not make a direct comparison between PHATE and SPADE.

Monocle2 also fits a tree to cell clusters using the DDRTree algorithm [112,113] as a default. We compare Monocle2 to PHATE in Figures S7 and S3. For the artificial tree in Figure S3A, Monocle2 successfully identifies the branches of the tree. However, for the bone marrow scRNA-seq data in Figure S7A, Monocle2 fails to detect several branches that are visible in the PHATE visualization. At the same time, Monocle2 shows several branches that are not detected by PHATE. The number and location of these branches vary from run to run on the same data with the same settings. Thus it is difficult to determine if the branches shown by Monocle2 are spurious or not.

Similar results are obtained when applying Monocle2 to the EB scRNA-seq data where the number and location of the branches within the Monocle2 visualization differ drastically from run to run. Thus it is difficult to determine the underlying structure of this data using Monocle2. In contrast, for the same set of parameters, PHATE produces the same results with each run while preserving the relative relationships between different branches directly in the visualization based on their proximity.

Furthermore, since Monocle2 enforces a tree structure on the data, the application of Monocle2 to data without a tree structure produces unintelligible results, giving tree-shaped visualizations for clustered (Figures S3C, H, I and S7F), circular (Figure S3E) and planar (Figure S3F) data.

## C PHATE for Data Exploration on Various Data Types

PHATE can be used for exploratory data analysis on a variety of data types. Here we present some examples of PHATE applied to other datasets including non-single-cell data such as microbiome data, SNP data, facial images, Hi-C chromatin conformation maps, and Facebook network data.

### C.1 PHATE on High-dimensional High-throughput Data

As a general dimensionality reduction method, PHATE is applicable to many datatypes. Here we show that PHATE reveals and preserves global transitional structure in mouse bone marrow scRNA-seq data, microbiome data, human SNP data, and (non-biological) image data.

#### Bone Marrow scRNA-seq Data Reveals New Structure

Figure S6D shows the color-coded 3D PHATE embedding and gene expression matrix for scRNA-seq data from mouse bone marrow. This data is enriched for myeloid and erythroid lineages and was organized into clusters in [21], as shown in Figure 5A. Here, we show that PHATE reveals a continuous progression structure instead of cluster structure and illustrates the connections between clusters. The PHATE embedding shows a continuous progression from progenitor cell types in the center to erythroid lineages towards the right and myeloid lineages towards the left. The expression matrix shows increasing expression of erythroid markers in the rightmost branches (branches 4, 5, and 6) such as hemoglobin subunits Hba-a2 and Hbb-b1 as well as heme synthesis pathway enzyme Cpox as the lineage progresses to the right. Towards the left in branches 1 and 2, we see an enrichment for myeloid markers, including CD14 and Elane, which are primarily monocyte and neutrophil markers, respectively.

In addition, PHATE splits the erythrocytes into three branches not distinguished by the authors of [21]. These branches show differential expression of several genes. Branch 6 is more highly expressed in Gata1 and Gfi1B, both of which are involved in erythrocyte maturation. Branch 4 is also more highly expressed in Zfpm1 which is involved in erythroid and megakaryocytic cell differentiation. Additionally, branches 4 and 5 are more highly expressed in Car2, which is associated with the release of oxygen. Given these differential expression levels, it is likely that the different branches correspond to erythrocytes at different levels of maturity and in different states [114–120]. In addition, the branches at the right have high mutual information with CD235a, which is an erythroid marker that progressively increases in those lineages.

PHATE in 3 dimensions more clearly reveals a separation of the myeloid lineages to the left into two branches. Cd14 and Sfpi1 are both more highly expressed at the beginning of branch 2 than in branch 1, suggesting that branch 2 is associated with monocytes while branch 1 is associated with neutrophils.

We note that due to the lack of common myeloid progenitors in this sample, one may expect to see a gap between the monocytes and megakaryocyte lineage, since PHATE does not artificially connect separable data clusters (see Figure S3C). However, we note that all but one of the 12 embeddings of this data in Figure S7A also lack a gap between these trajectories, suggesting there is a genuine similarity between cells in the monocyte and megakaryocyte lineages, whether due to biological similarities or a technical artifact of sequencing (e.g., contamination.)

#### PHATE on Facial Images

To demonstrate that PHATE can also be used to learn and visualize the underlying structure of nonbiological data, we applied PHATE to the Frey Face dataset used in [6]. This dataset consists of nearly 2000 video frames of a single subject’s face in various poses. Figure S8A shows a 3D visualization of this dataset using PHATE, colored by time. Multiple branches are clearly visible in the visualization and each branch corresponds to the progression of a different pose. A short video highlighting two of these branches is available at https://www.youtube.com/watch?v=QMCWsKNgvHI. The video highlights the continuous nature of the data as points along branches correspond to transitions from pose to pose.

#### PHATE on Microbiome Data Reveals Archetypal Structure

Recently there have been many studies of bacterial species abundance in the human intestinal tract, saliva, vagina and other membranes as measured by sequencing of the 16S ribosomal-RNA-encoding gene (16s sequencing) or by whole genome shotgun sequencing (metagenomics). However, most analysis of microbiome data has been limited to clustering and PCA. Here we use PHATE to analyze microbiome data from the American Gut Project.

First we note that PCA applied to 9660 fecal, oral, and skin samples (Figure S8B left) results in an undifferentiated cloud with two density centers corresponding to fecal samples on the right and oral/skin samples on the left. In contrast, PHATE shows branching structures with 4 branches emanating from a point of origin for fecal sample, and additional structures on the right that differentiates between skin samples, which form their own progression, and oral samples, which again result in several branches. Figure S8C shows the PHATE embedding colored by two genera (bacteroides and prevotella) and a phylum (actinobacteria) of bacteria on the same 9660 samples. These figures show that the Bacteroides genus of bacteria is almost exclusively found in the fecal samples. The Prevotella genus of bacteria is found in certain stool and oral samples while the Actinobacteria phylum is primarily found in the oral and skin samples. This is consistent with the work in [121] which showed that different genera and phyla of bacteria are prevalent in the different body sites.

Upon “zooming in” to the 8596 fecal samples in Figure S8D, we see 4 major branches, instead of the three enterotypes reported in previous literature [122], with highly expressed Firmicutes, Prevotella, Bacteroides and Verrucomicrobia respectively. Furthermore, the Fermicutes/Bacteroides branches seem to form a smooth continuum with samples falling into various parts of a triangular simplex shape typically seen in archetypal analysis [123,124]. This shows that individuals can exist as mixed phenotypes between archetypal bacterial states as well as in a continuum with more or less prevalence for each of these states.

#### PHATE on SNP Data Reveals Geographic Structure

To demonstrate PHATE on population data, we examined a dataset containing 2345 present-day humans from 203 populations genotyped at 594,924 autosomal SNPs with the Human Origins array [125]. In Figure S8E, we see that as compared to PCA, the PHATE embedding shows clear population structures, such as the near eastern Jewish populations near the bottom (Iranian and Iraqi Jews, Jordanians), with further branches showing progression within the same population, such as the Jordanian population show as orange diamonds (see Figure S8F for population labels.) Further, PHATE shows a global structure that mimics geography, with European populations generally towards the top and Near Eastern populations towards the bottom. Thus PHATE shows that the occurrence and structure of these SNPs follows a progression based on geography and population divergence. PCA tends to crowd populations together into two linear branches, without clearly distinguishing between population groups or showing population divergence.

### C.2 PHATE on Connectivity Data

Thus far we have used PHATE to embed high-dimensional data. That is, we have mapped the original feature space of the data to the PHATE dimensions. However, the PHATE algorithm also allows for embedding any data that exists in either an inner product space or a metric space. In other words, we can apply PHATE to data that is naturally described by distances or affinities instead of features. For example, network data (i.e. graphs) can be embedded using PHATE simply by skipping the initial steps of the algorithm that go from the original feature space to an affinity matrix. We can thus replace the affinity matrix with the network (as represented by an affinity matrix) and proceed with the rest of the PHATE algorithm.

There are abundant examples of natively networked data in biology, such as chromatin conformation contact maps (Hi-C), gene/protein interaction networks, and neural connectivity data such as fMRI. Outside of biology, network data is also prevalent. A common example is social network data that contain information of friend-communities (e.g. Facebook) or interest-communities (e.g. Twitter). We show that PHATE provides a visualization of network data that emphasizes major structure (i.e. pathways) in the network, better than typical graph layout methods.

#### PHATE on Hi-C Data Reveals Spatial Chromatin Structure

We use PHATE to visualize human Hi-C data from [15] by using the Hi-C contact map as the affinity matrix in the PHATE algorithm. The contact map gives the frequency with which genomic locations are observed in spatial proximity. Hi-C contact maps are typically visualized using the matrix directly – often using the 45 degree counterclockwise rotated upper triangle part of the matrix [15]. While this depiction can show chromosomal domains, it is not a reconstruction of the actual spatial structure of the chromatin. In contrast, the PHATE embedding reconstructs the relative positions of the genomic locations both locally and globally in such a way that the embedding represents an actual projection of the spatial structure within and between chromosomes. As a result, we get an intuitive visual of the Hi-C contact map that not only shows the topological domains present in the data but also how they are connected to one another. Figure S9A shows a 3D PHATE embedding of the chromatin, colored by chromosome. We see that the embedding resembles the fractal globule structure proposed in [126], with the total chromatin organized in a spherical shape and individual chromosomes mostly connected within themselves.

PHATE can also effectively visualize a single chromosome. Figure S9B shows PHATE just on chromosome 1 contact map at 10 kilobase (kb) resolution. Each point corresponds to a genomic fragment and is colored by its location within the genome. In this visualization, multiple “folds” are clearly visible.

To validate that the PHATE embedding of Hi-C data is meaningful we color the embedding by the ChIP-seq signals of several chromatin modification markers. Figure S9C shows a 2D PHATE embedding of chromosome 1 colored by various methylation and acetylation markers (ChIP-seq [127], dataset ENCSR977QPF). Histone methylation and acetylation play an important role in global gene regulation via control of chromatin organization. They are often used in combination with Hi-C data to investigate the open or closed structure of chromatin [15], where the ChIP-seq signal of various histone modification markers correlates with so called topologically associated domains or TADs. Figure S9C shows an organized methylation (H3K27me3 and H3K4me2) and acetylation (H3K9ac and H3K27ac) signal on PHATE, with clusters of similar intensity of the marker, suggesting that the PHATE embedding has biological meaning with respect to the spatial organization of the chromatin.

These results indicate that PHATE provides a visualization of Hi-C data that captures more spatial information compared to directly looking at the contact map.

#### PHATE on Facebook Data Reveals Super Connectors

We visualize Facebook network data using PHATE. The data [128] consist of networks of friends. This network graph is directly converted into a 0-1 affinity matrix and fed to PHATE. Figure S9D compares the PHATE visualization of the network to a force-directed layout, a common method for visualizing network data. In both plots, edges (friends) are shown and each node/person is colored by degree (the number of friends a person has). The PHATE embedding clearly shows multiple branching structures in the network, that are not visible in the force-directed layout which shows a T shape.

Several subnetworks and important nodes (super-connectors) between subnetworks are visible in the PHATE embedding that are not visible in the force-directed layout visualization. For example, in the top-left corner of the PHATE visualization of the entire network, a single node connects the top-left group of people to the remainder of the network. Several other important nodes can be identified that bridge the gap between the left section of the network and the center region. Thus PHATE can be used for visualizing network data and identifying important features of the network.

We also show that PHATE can be used to find more structure within subnetworks as identified by the friend networks of selected individuals (referred to as ego nodes in [128]) in Figure S9D. Again, we find that PHATE finds more structure in subnetworks.

Therefore, we see that PHATE can be used to visualize any type of data, high-dimensional or featureless. Further, we see that PHATE will emphasize continuous transitional structure, while maintaining separation between clusters in these datasets.

## D Detailed EB Data Results

Here we provide a more detailed analysis of the EB data based on the PHATE visualization. We examined markers of cell proliferation as they are known to be correlated with the time trend. Undifferentiated hESCs have very fast cell cycles with prominent S and M phases and very short G1 phase, while differentiating cells have longer G1 phase and cycle slower [129]. Indeed, we found that S-phase cyclins *CCNE1* and *CCNE2* had the highest level of expression in the earlier time points while G1-specific CDK inhibitors *CDKN1A, CDKN1B* and *CDKN1C* were induced at later time points as shown in Figure S12B.

To examine localized lineages and branching structures, we visualized the expression of known germ layer markers on PHATE (as in Figure S12B), resulting in the germ layer specification chart shown in Figure 6B. Note that in the remaining description we refer to the branches based on the labels shown in Figure 6Aiii. These branches are identified as follows. First, we performed clustering of the data using the *k*-means algorithm on ten-dimensional PHATE (Figure 6Ai) to identify differentiation intermediates within each germ layer branch. This can be viewed as a variant on spectral clustering using PHATE dimensions instead of the eigenvectors of a Laplacian matrix [130,131]. We note that clustering could also be applied directly to the potential distances and other clustering approaches can be used including hierarchical clustering, Louvain clustering [132,133], and others [134,135]. For each cluster, we performed differential expression analysis by comparing, for each gene, its expression distribution within the cluster to the expression distribution in all other cells (i.e., the “background” expression distribution) using the earth mover’s distance or EMD (Figure 6D), a measure of dissimilarity between probability distributions [87] (see Appendix A.1.5 for details). We confirmed that the genes expressed in several differentiation intermediates, which we identified by manual inspection, can also be identified based on the EMD scores of the corresponding cluster. For example, *SOX10* has a high EMD score in clusters 14 and 15, which corresponds with neural crest cells. Similarly, *EOMES* has a high EMD score in clusters 4, 17, and 20, which corresponds with the primitive streak cells. We also observed several canonical differentiation intermediates that were previously defined based on studies in mice. These include anterior NE and NC branches in the ectodermal lineage, anterior EN, cardiac and lateral plate ME, and hemangioblasts in the mesendoderm lineage.

The advantage of clustering over manual population extraction is that the resolution of the clustering can be chosen by setting *k* (the number of clusters), and very finely grained populations can be extracted and characterized. Here, we characterized thirty clusters, at finer resolution than the manual analysis above. However, a larger *k* can be used for even finer exploration of the data. These clusters can then be characterized on the basis of gene expression for FACS sorting and further experimentation as we show in the next section.

ESC-specific transcripts *POU5F1, NANOG*, and *DPPA2/4* were highly expressed in cells located at the left-most part of sub-branch i, indicating that this is the starting point of the data (Figure S12B). As cells traveled along sub-branch i into sub-branch ii, the epiblast marker *OTX2* was upregulated and then downregulated, followed by a sharp increase in markers associated with primitive streak (*CER1, MIXL1, EOMES*, and *T*), indicating that mesoderm (ME) and endoderm (EN) differentiation begins in this region. These markers continued to be highly expressed in cells located within the sub-region of branch v, which also exhibits high expression of the anterior EN markers *FOXA2* and *SOX17*, while the posterior EN transcripts *CDX2, NKX2-1*, and *KLF5* were expressed in the adjacent region of the same branch. ME transcripts *NKX2-5, TNNT2, TAL1, TBX5*, and *TBX18* were specifically expressed in different sub-branches vi-x.

Further along sub-branch ii, past the ME/EN initiation region, the neuroectoderm (NE) markers *PAX6, ZBTB16, SOX1, SOX2, NEUROG1*, and *DCX* were expressed throughout branch iv, while branch iii was positive for the neural crest (NC) markers *PAX3, FOXD3, SOX9*, and *SOX10*. Figures S6E and S12C further show how the expression of these and other selected genes changes at different points within the branches. From these analyses, it is clear that PHATE successfully resolved all known germ layer branches based solely on the scRNA-seq data without any prior assumption about the structure of the data.

Furthermore, the PHATE embedding suggested three novel differentiation trajectories. Within the ectodermal lineage, a distinct *GBX2+ZIC2/5-HOXD1+GLI3+* NE progenitor gave rise to a bi-potent *HOXA2+HOXB1+* precursor that separated into the NC branch and neural progenitor (NP) branch. The latter, in turn, split into at least five specialized neuronal subtypes. Within the EN branch, the canonical *EOMES+FOXA2+SOX17+* EN precursor was clustered together with the novel *EOMES-FOXA2-GATA3+SATB1+KLF8+* precursor, which further differentiated into cells expressing posterior EN markers *NKX2-1, CDX2, ASCL2*, and *KLF5*. Importantly, both the canonical and non-canonical EN cells expressed *ARID3A*, which can thus be defined as a pan-EN marker in humans. Moreover, we have identified a novel *T+GATA4+ CER1+PROX1+* cardiac precursor cell type within the ME lineage that gives rise to a *TNNT2+* cell via a *GATA6+HAND1+* differentiation intermediate.

---

**Supplemental Table S1:** Branches of the EB data identified for analysis and the corresponding clusters (Figure 6Bi) and sub-branches (Figure 6Biii) that comprise them.

| Branch | Sub-branches | Clusters |
| --- | --- | --- |
| ESC | i, ii | 1-6 |
| Neural Crest | iii | 13-15 |
| Neuroectoderm | iv | 7-12 |
| Endoderm | v | 16-20 |
| Mesoderm | vi-vii | 20-30 |

**Supplemental Figure S1.**
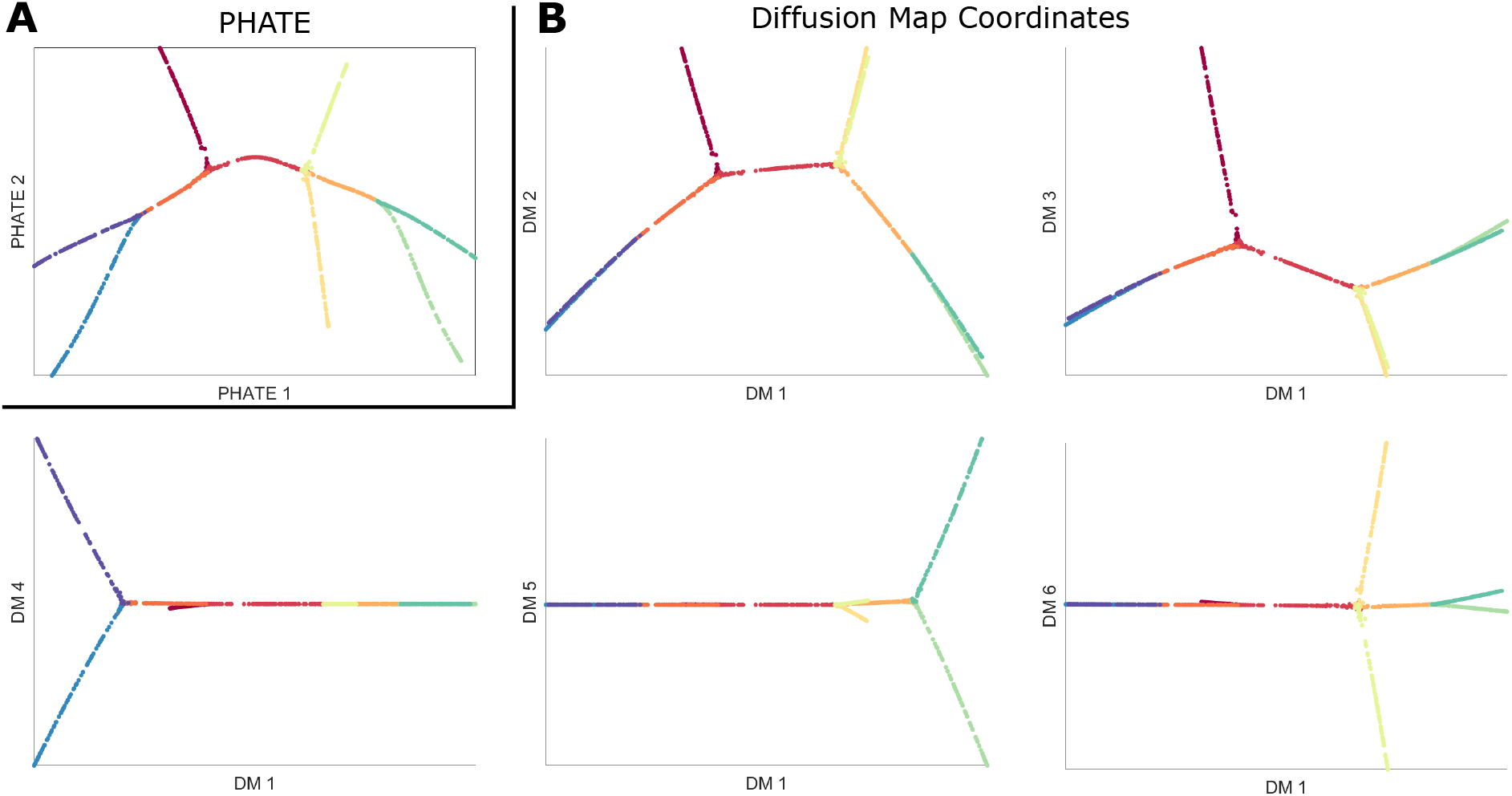
Comparison of PHATE to DM. (**A**) PHATE applied to the artificial tree data. Only two PHATE coordinates are needed to separate all branches. (**B**) The first six diffusion map coordinates of the artificial tree data. At least five of these coordinates are necessary to separate all of the branches.

**Supplemental Figure S2.**
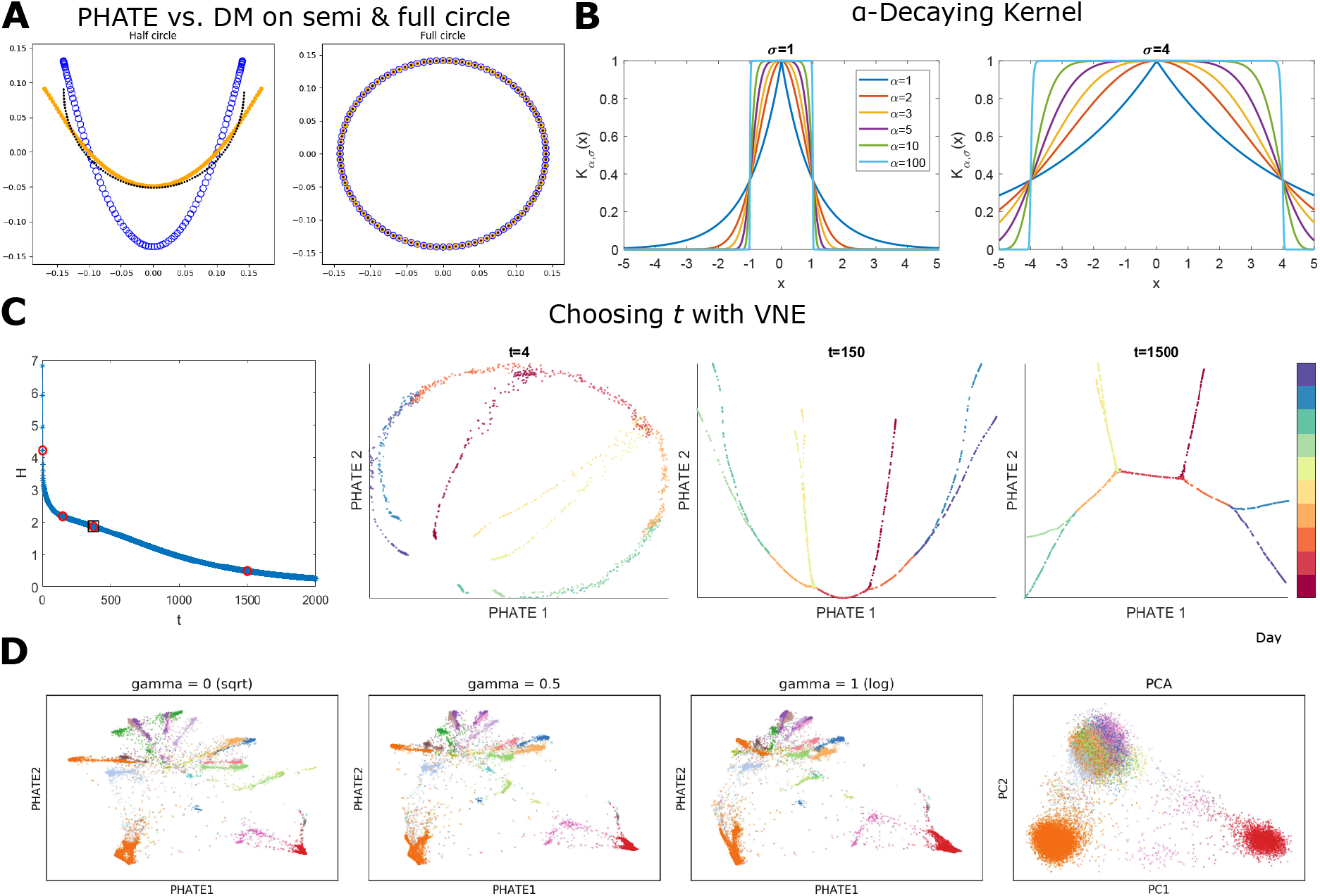
Impact of potential distances and PHATE parameters on the resulting visualization. (**A**) Comparison of Diffusion Maps (blue) and PHATE (orange) embeddings on data (black) from a half circle (left) and a full circle (right). Both the data and the embeddings have been centered about the mean and rescaled by the max Euclidean norm. For the full circle, both embeddings are identical (up to centering & scaling) to the original circle. However, for the half circle, the Diffusion Maps embedding (blue) suffers from instabilities that generate significantly higher densities near the two end points. The PHATE embedding (orange) does not exhibit these instabilities. (**B**) The *α*-decaying kernel 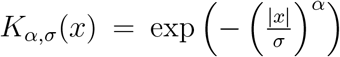 as a function of *x* for different values of *α* and *σ* = 1 (left) and *σ* = 4 (right). As *α* increases, *K_α,σ_*(*x*) becomes more constant for *x* ∈ (−*σ, σ*) and the tails of the kernel become lighter (i.e., decay to zero more quickly) for *x* ∉ (−*σ, σ*). (**C**) Demonstration of the effect of the scale *t* on the PHATE visualization for the artificial tree data colored by branch. The first column shows the VNE **H*(*t*)* (see Eq. 7) of the diffusion affinities as a function of the time scale *t*. The other columns give the PHATE visualization with different values of *t*. The red dots in the first column indicate the values of *t* chosen for the plots. The red dot surrounded by a black box indicate the chosen value of *t* for the visualization in Figure 1B of the artificial tree data. Values of *t* that are too low can give noisy visualizations while very high values of *t* can result in a loss of information in the visualization. (**D**) Visualization of scRNAseq on mouse retinal bipolar neurons data using different informational distances defined via the parameter *γ*.

**Supplemental Figure S3.**
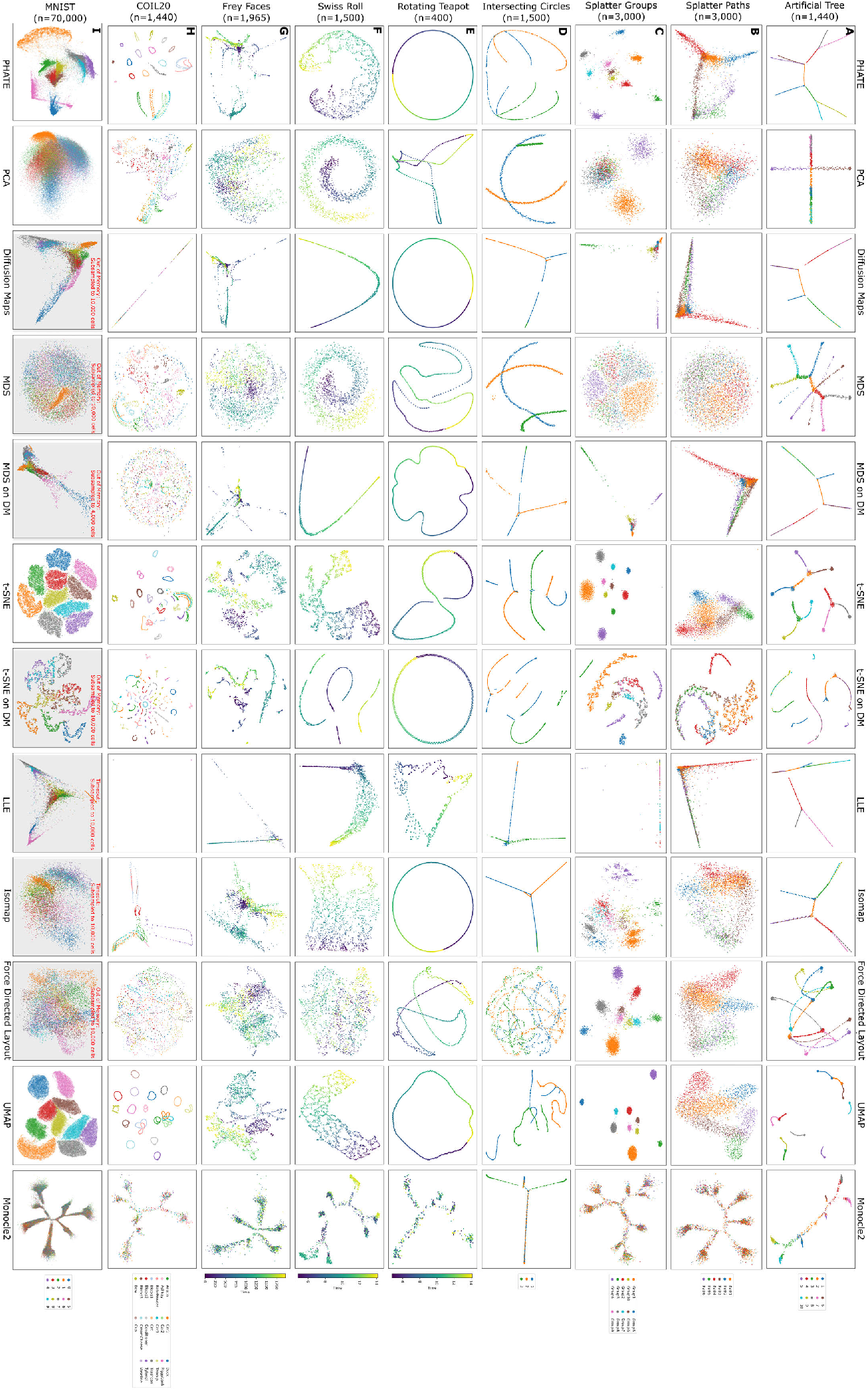
Comparison of PHATE to various methods on multiple artificial and non-biological datasets. See B for discussion.

**Supplemental Figure S4.**
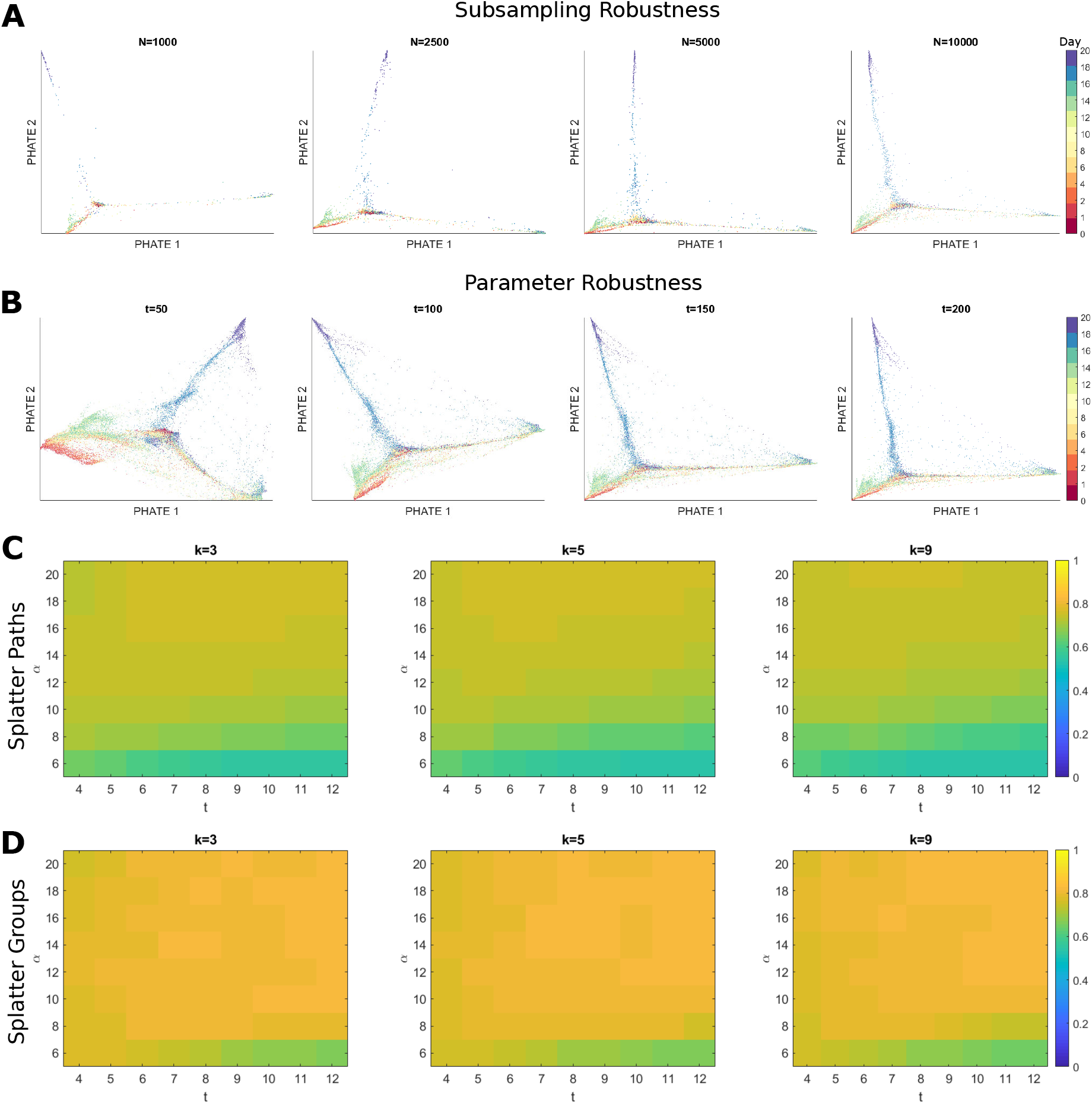
Visual and quantitative demonstrations of the robustness of PHATE to subsampling and the choice of parameters. (**A**) The PHATE visualization for the iPSC mass cytometry dataset from [16] with varying number of subsample sizes *N*. The main branches present for *N* = 10000 are also visible for the other values of *N*, demonstrating that the PHATE embedding is robust to the size of the subsample. (**B**) The PHATE visualization for the iPSC CyTOF dataset from [16] with varying scale parameter *t*. The embeddings for all *t* preserve the branching structure and the visualizations are very similar to each other, demonstrating that the embedding is robust to the choice of *t*. (**C**) Heatmap of the Spearman correlation coefficient between geodesic distances of the ground truth data and the Euclidean distances of the PHATE visualization applied to the simulated paths dataset using Splatter [22]. The results are presented using different values for *k, t*, and *α*. The value of *t* selected using the kneepoint method in this case is 8. (**D**) Heatmap of the Spearman correlation coefficient between geodesic distances of the ground truth data and the Euclidean distances of the PHATE visualization applied to the simulated groups dataset using Splatter [22]. The results are presented using different values for *k, t*, and *α*. For both the groups and paths datasets, the results are very stable for *α* ≥ 10. The value of *t* selected using the kneepoint method in this case is 8.

**Supplemental Figure S5.**
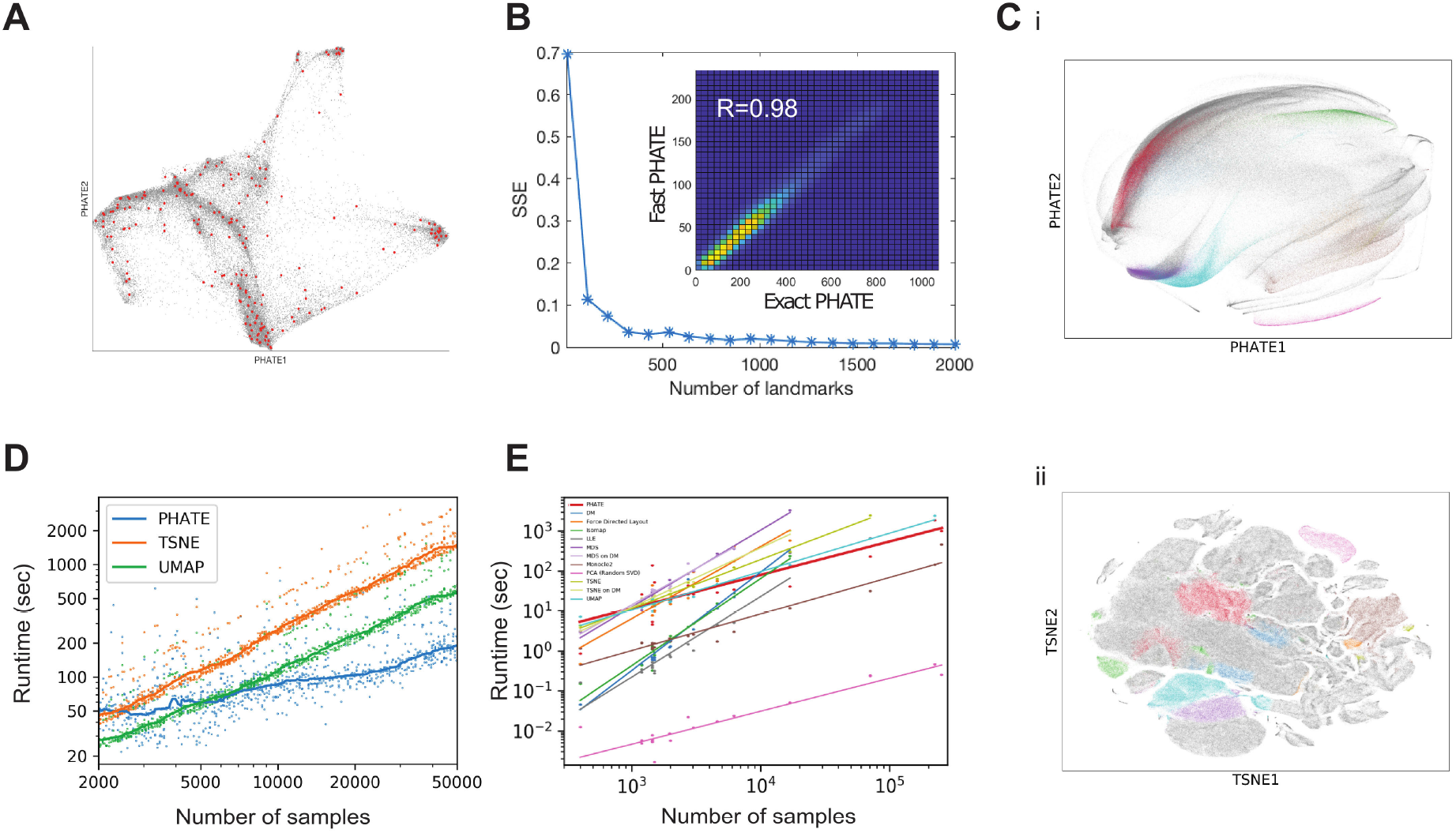
Scalability tests of PHATE. (**A**) Scalable PHATE embedding of iPSC CyTOF data with a subset of the landmarks shown in red (200 out of 2000). (**B**) Robustness of PHATE to the number of landmarks chosen. PHATE on the EB data computed using increasing numbers of landmarks (X-axis) was compared to exact PHATE, i.e. without landmarks. Comparison was done using Procrustes analysis (optimal linear transformation) and the sum of squared error (SSE, Y-axis) is shown. To ensure a stable embedding that accurately approximates exact PHATE we choose 2000 landmarks as default. The inset shows the histogram of pairwise distances in the visualization computed using fast PHATE (2000 landmarks) on the EB data vs. the pairwise distances from exact PHATE. The correspondence and correlation coefficient are very high. (**C**) PHATE and t-SNE embeddings of a 1.3 million mouse brain cell dataset from 10X genomics [85]. The PHATE embedding was calculated with 2000 landmarks and completed in three hours. A subset (10 of 60) of the clusters provided by 10X are shown in color, the rest in gray. t-SNE shatters the cluster structure, while PHATE retains clusters as contiguous groups of cells. (**D**) Runtime of PHATE, t-SNE and UMAP on increasingly large subsamples of the EB data. Runtime was averaged across four runs. (**E**) Runtime of 12 visualization methods shown in Figures S3 and S7 across 19 datasets and corresponding line of best fit for each method. Where a method ran out of memory or took longer than one hour, the runtime is not shown and linear fits are cut off accordingly.

**Supplemental Figure S6.**
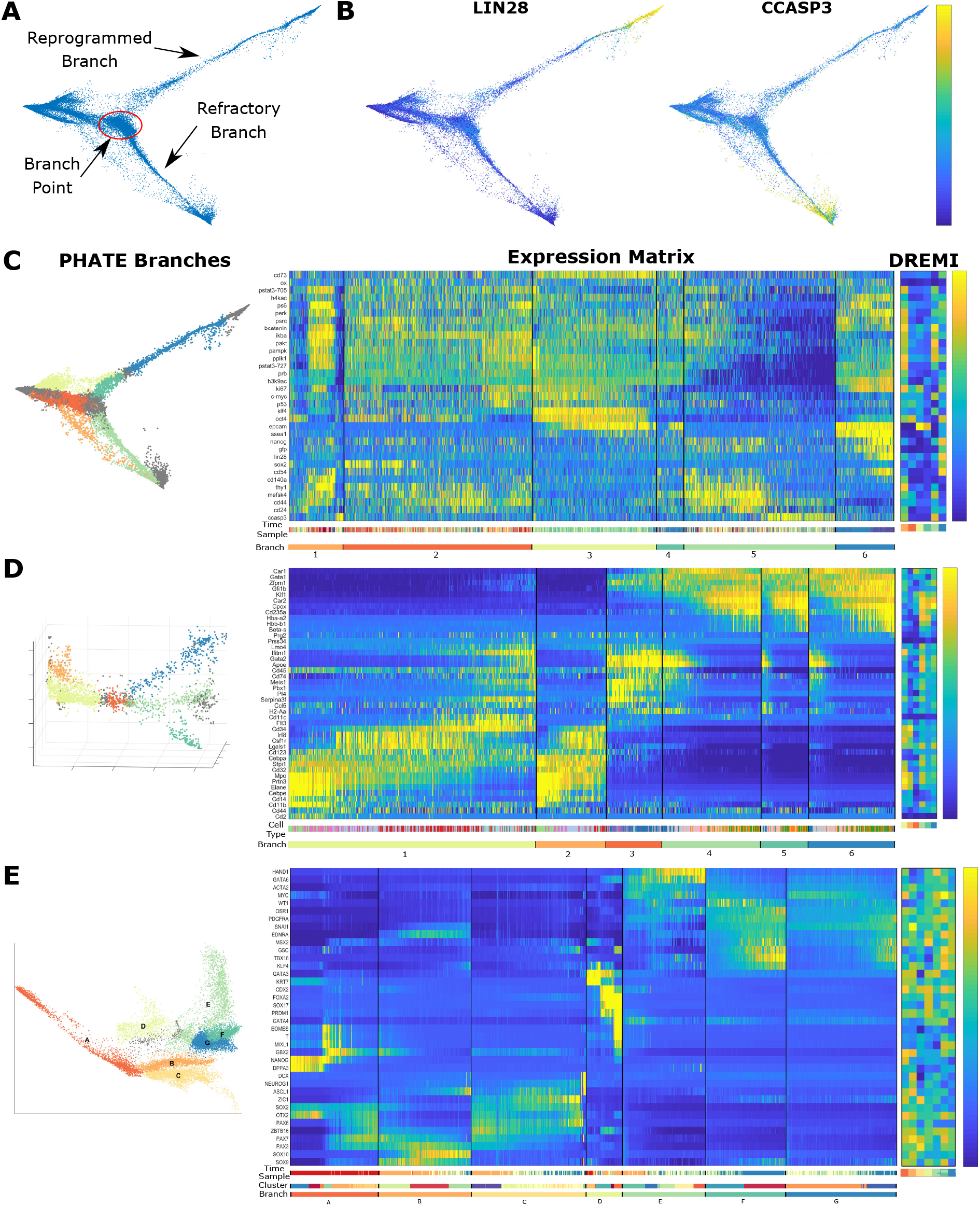
Annotated PHATE visualizations of CyTOF iPSC data [16] and branch expression analysis. (**A**) The primary branch point between the two major branches (reprogrammed and refractory) of the data is highlighted. (**B**) The PHATE visualization colored by LIN28 (a marker associated with the transition to pluripotency [32]) and CCASP3 (associated with cell apoptosis). LIN28 expression is limited to the reprogrammed branch while CCASP3 is primarily expressed in the refractory branch, indicating that the failure to reprogram may initiate apoptosis in these cells. (**C**) Analysis of branches on the PHATE embedding for the iPSC CyTOF data [16], (**D**) bone marrow scRNA-seq dataset from [21], and (**E**) newly generated embryoid body scRNA-seq data. (Left) The PHATE visualization with identified branches. (Middle) Expression level for each cell ordered by branch and ordering within the branch. Cell ordering is calculated using Wanderlust [26] starting on the left-most point of each branch. Expression levels are z-scored for each gene. A colorbar is given below the expression matrices that identifies each branch and (in the case of the bone marrow scRNA-seq data) cell type. (Right) DREMI scores [31] between gene expression levels and cell order within each branch. MAGIC [92] is applied first in (**D**) and (**E**) to impute missing values using the same kernel used for PHATE and smaller *t*. For branch analysis of the bone marrow data in (**D**), we used 3 PHATE dimensions to obtain clearer branch separation.

**Supplemental Figure S7.**
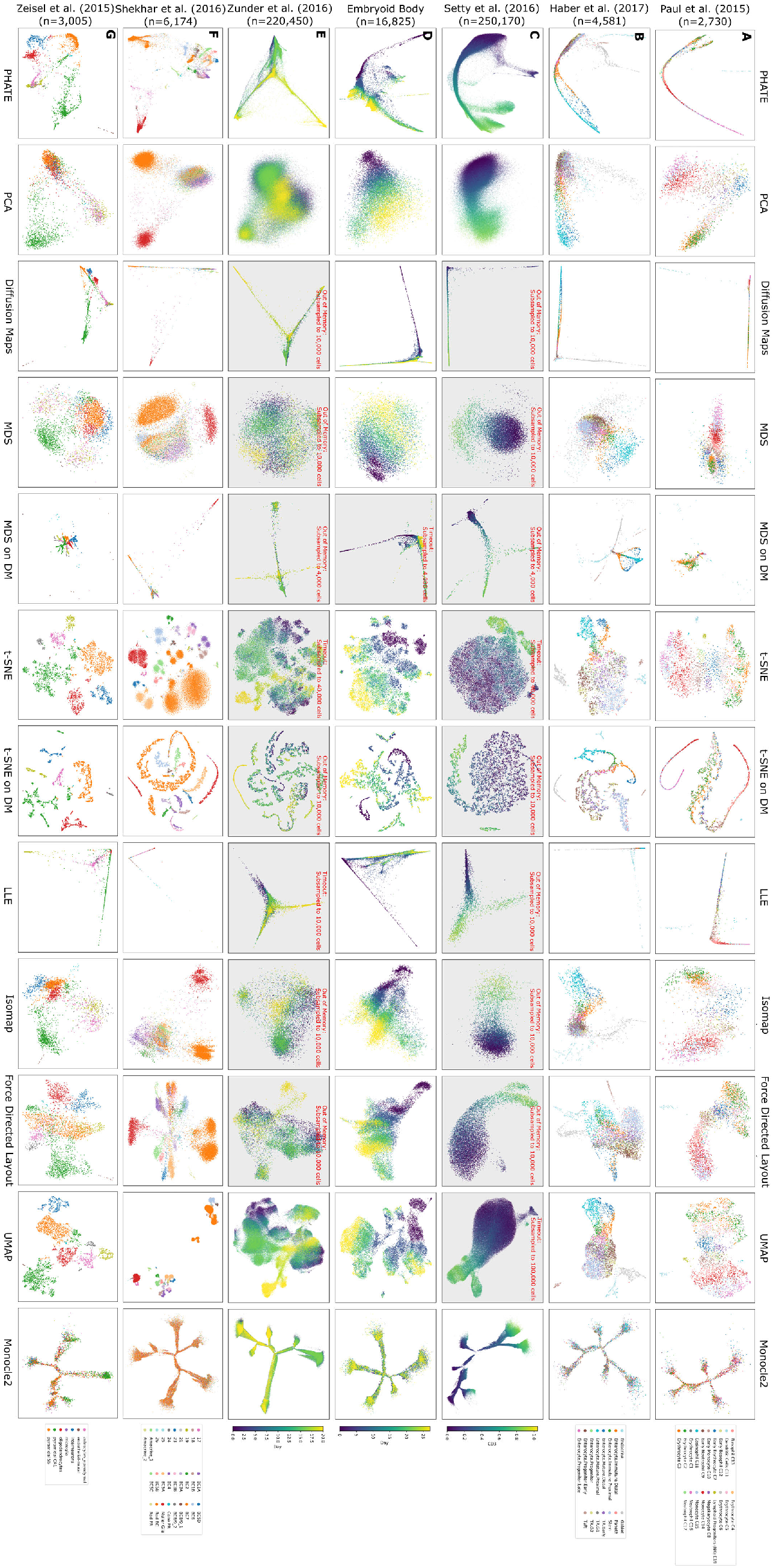
Comparison of PHATE to various methods on multiple biological datasets. See B for discussion.

**Supplemental Figure S8.**
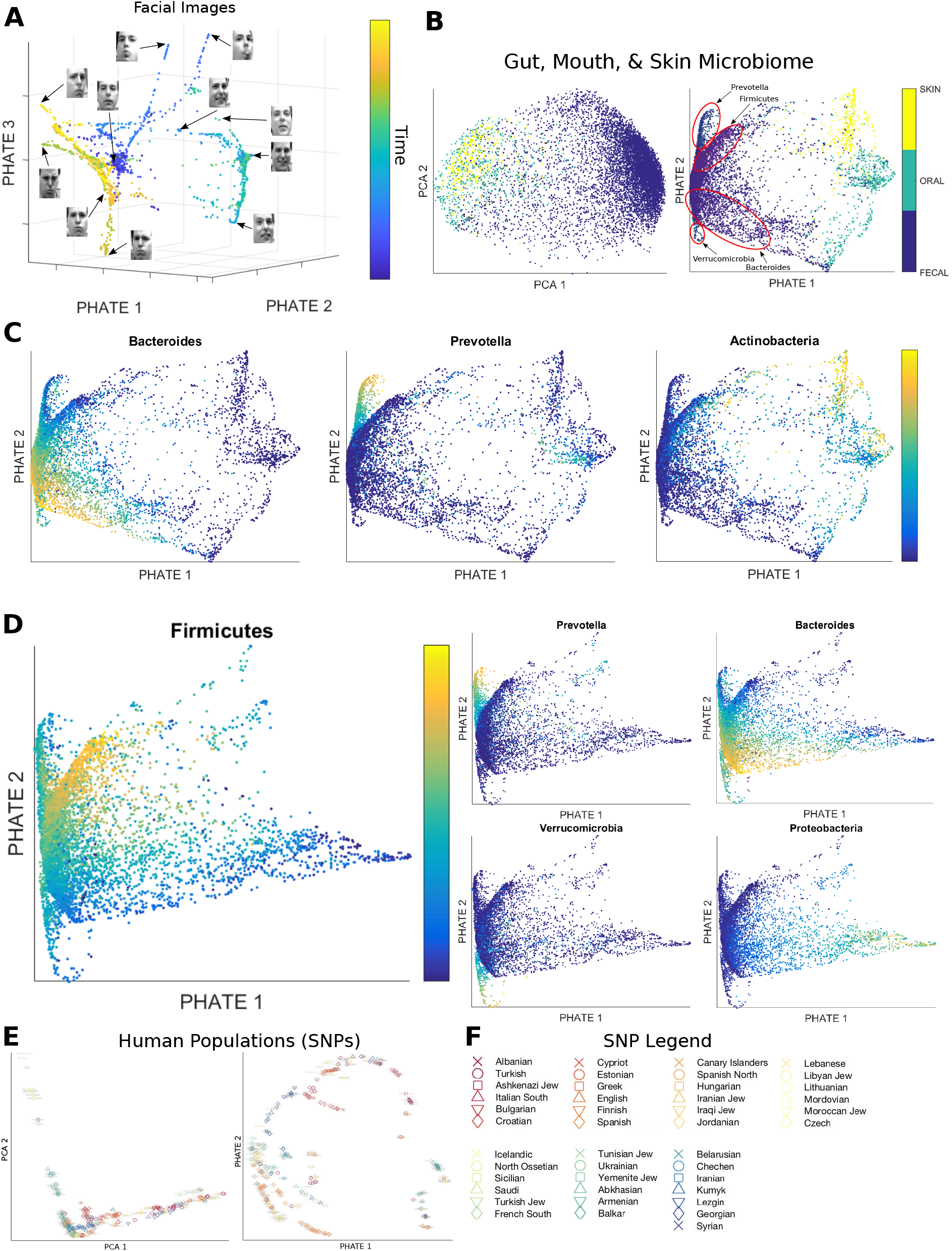
PHATE reveals structure in a variety of high-dimensional datasets. (**A**) A 3D PHATE visualization of the Frey Faces dataset used in [6]. Points are colored by time within the video. Multiple branches corresponding to different poses are clearly visible. (**B**) PCA and PHATE embeddings of microbiome data from the American Gut project, colored by body site, and branches annotated by their dominant genera or phyla. (**C**) The PHATE embedding of data from the American Gut project colored by 2 genera (bacteroides and prevotella) and a phylum (actinobacteria) of bacteria. (**D**) The PHATE embedding of only the fecal samples from the American Gut project colored by various genera (bacteroides and prevotella) and phyla (firmicutes, verrucomicrobia, and proteobacteria) of bacteria. Each PHATE branch is associated with one of these bacteria groups. (**E**) PCA and PHATE embeddings of SNP data from the Human Origins dataset showing genotyped present-day humans from 203 populations [125] with the population legend in (**F**).

**Supplemental Figure S9.**
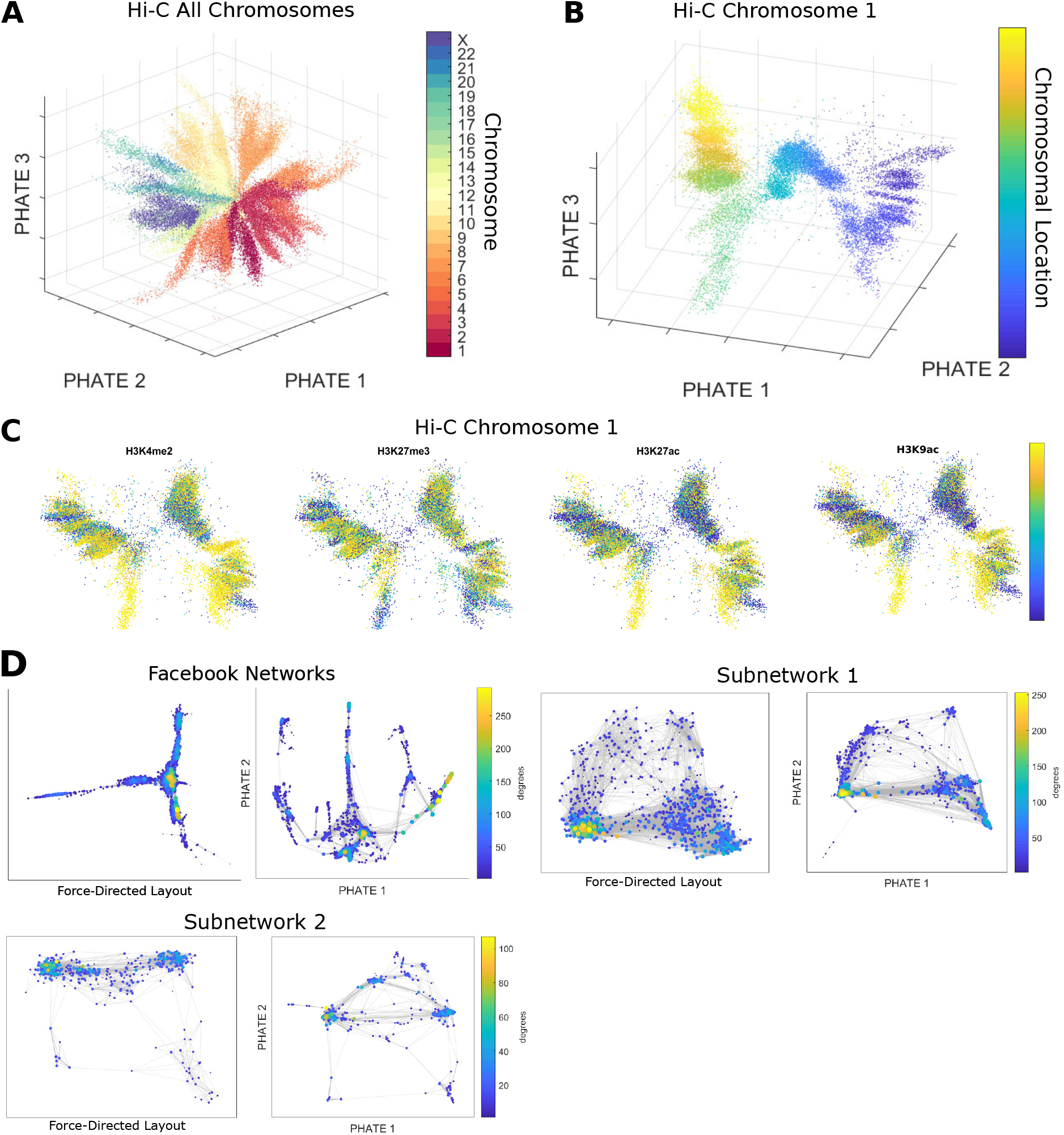
PHATE reveals structure in a variety of connectivity datasets. (**A**) 3D PHATE visualization of human Hi-C data [15] using all 23 chromosomes at 50 kb resolution, colored by chromosome. Each point corresponds to a genomic fragment. (**B**) PHATE visualizations of the same human Hi-C data in A for chromosome 1 at 10 kb resolution colored by chromosome location. (**C**) 2D PHATE visualization of human Hi-C data [15] for chromosome 1 at 10 kb resolution, colored by selected chromatin modification markers from ChIP-seq data. (**D**) Force-directed layout and PHATE visualizations of Facebook network data with data points colored by their degree (number of connections). The subnetworks are taken from the friend networks of selected individuals within the entire network. In all cases, PHATE reveals more structure.

**Supplemental Figure S10.**
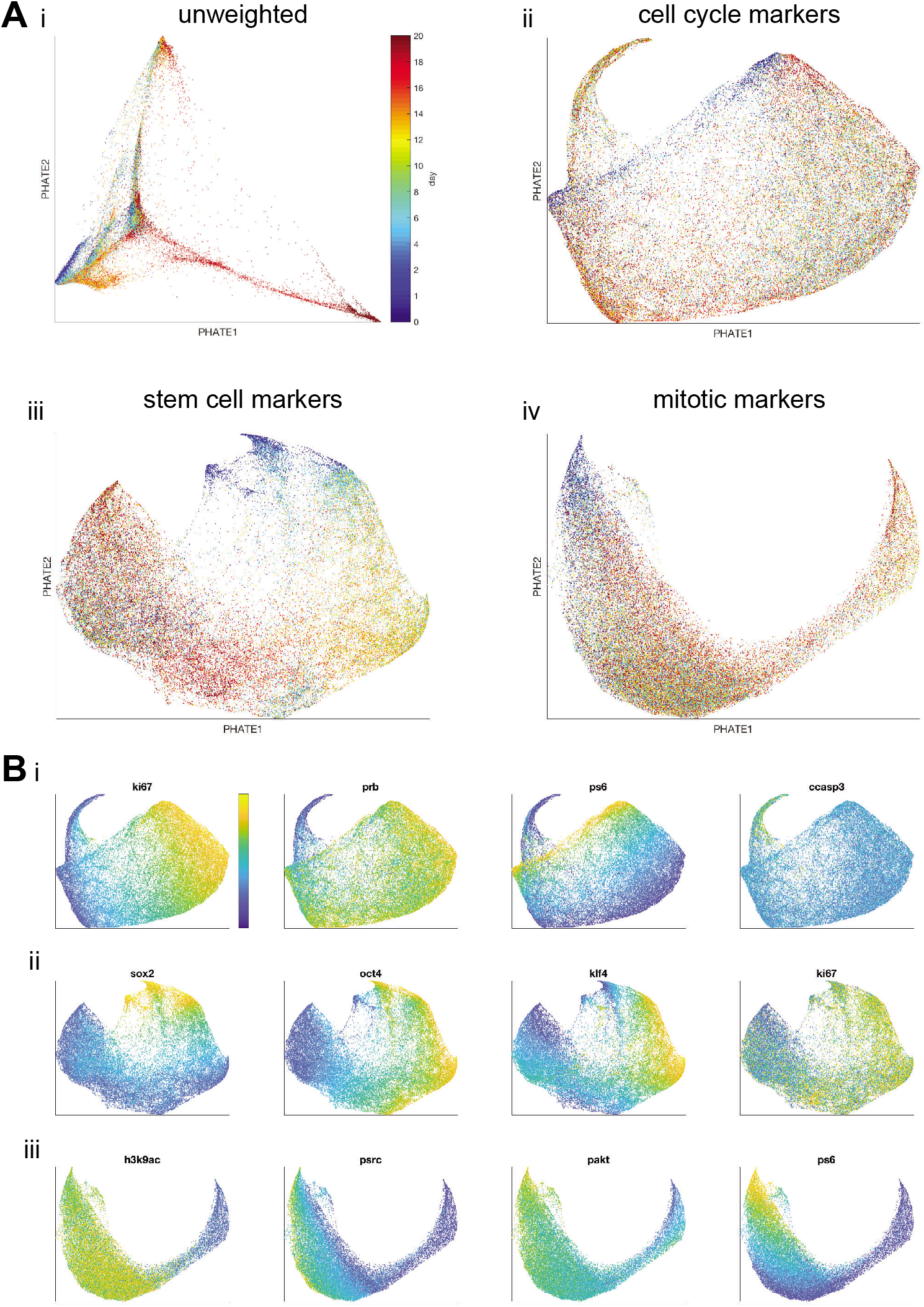
PHATE using reweighted distances to highlight specific biological processes or “views” of the data. (**A**) PHATE embedding of CyTOF iPSC data using (i) unweighted distances, (ii) distances after upweighting cell cycle markers, (iii) distances after upweighting stem cell markers, (iv) distances after upweighting mitotic markers. (**B**) PHATE colored by different markers (columns). From top to bottom: (i) PHATE cell cycle “view”, (ii) PHATE stem cell “view” (iii) PHATE mitotic “view”.

**Supplemental Figure S11.**
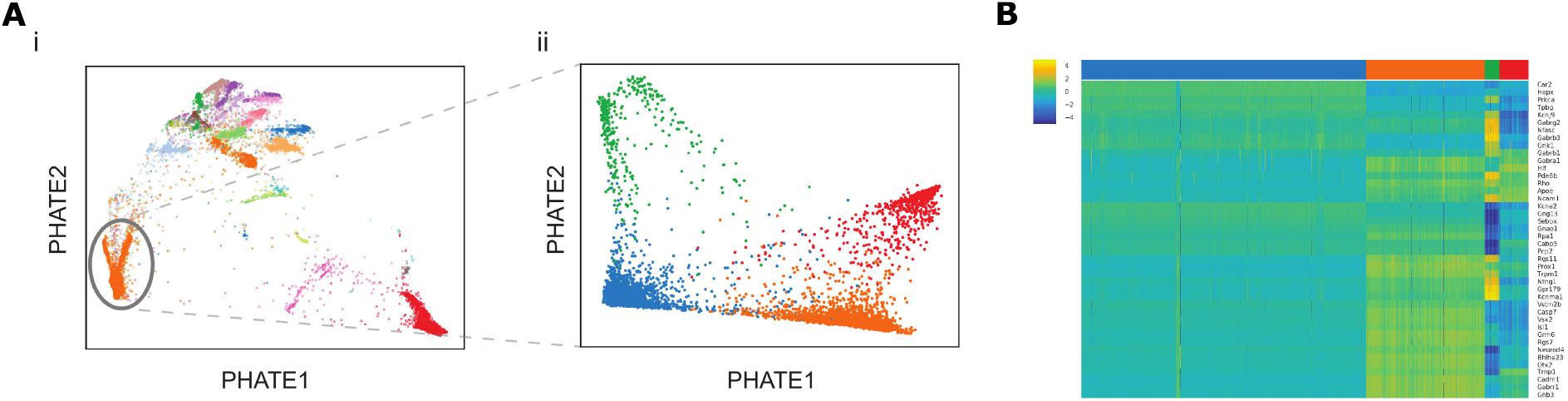
Additional analysis with PHATE. ((**A**) i. Initial PHATE embedding of scRNAseq on mouse retinal bipolar neurons. The rod bipolar cells cluster (cluster 1) is circled. ii. Subsequent PHATE embedding of cluster 1, colored by K means clustering to show heterogeneity within rod bipolar cells. (**B**) Transcriptional characterization of subtypes of rod bipolar cells, using known bipolar cell markers.

**Supplemental Figure S12.**
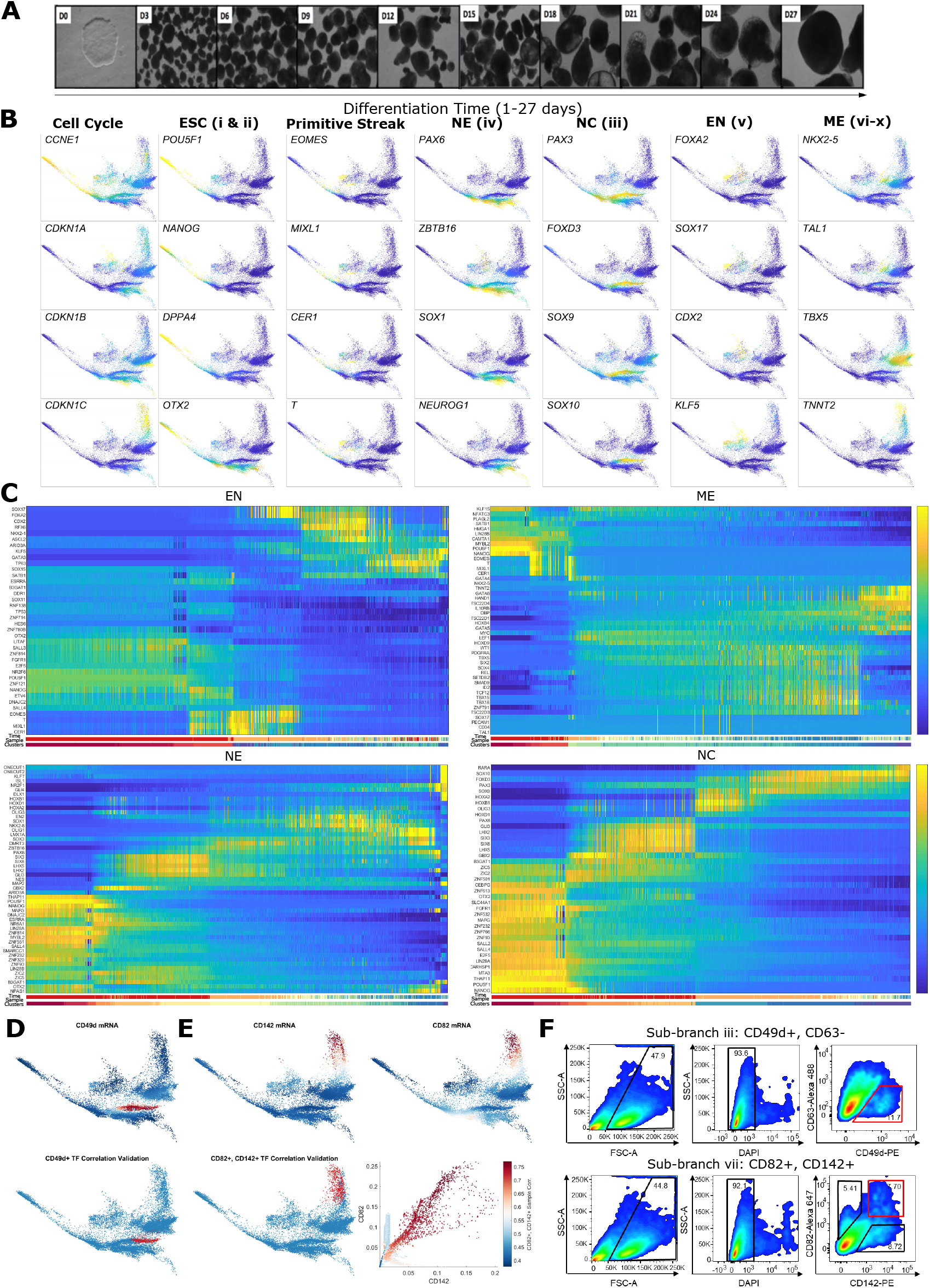
Further analysis of the EB scRNA-seq data. (**A**) Inverted images of hESCs and EBs at each timepoint of data collection. Structures of different densities are clearly visible late in the time course (D15-D27) indicating the formation of distinct cell types. (**B**) PHATE colored by expression levels of selected markers. (**C**) Heatmap showing gene expression level in each cell in four of the branches starting with ESC. Cell ordering is determined using Wanderlust [26]. Genes were selected either manually or by high DREMI scores [31] between gene expression and cell ordering. (**D**) PHATE colored by CD49d expression level from the scRNA-seq data (top) and by correlation between the scRNA-seq transcription factor expression and the CD49d-sorted bulk RNA-seq transcription factor expression per cell (bottom). (**E**) Same as **D**, with CD142 and CD82. The correlation coefficient is highest in branch vii, which is the branch with the highest CD142 and CD82 expression. Bottom right: Scatter plot of single cell expression levels between CD82 and CD142. Color corresponds to the correlation between the scRNA-seq expression and the CD142+CD82+ sorted bulk RNA-seq expression. The branch with highest correlation corresponds to cells that are positive in both CD142 and CD82. (**F**) Scatter plots showing the gating procedure for FACS sorting cell populations of sub-branch iii (CD49d and CD63) and sub-branch vii (CD82 and CD142).

## Footnotes

1 Recall the diffusion distance is simply the Euclidean distance in these coordinates

